# Glutamate (E299) is a key residue in the evolutionarily divergence of the SAM-dependent methyltransferases DnrK and RdmB in anthracycline biosynthesis

**DOI:** 10.1101/2024.05.03.592356

**Authors:** Moli Sang, Qingyu Yang, Jiawei Guo, Peiyuan Feng, Wencheng Ma, Shengying Li, Wei Zhang

## Abstract

A novel sub-class of *S*-adenosyl-L-methionine (SAM)-dependent methyltransferases catalyze atypical chemical transformations in the biosynthesis of anthracyclines, which include antineoplastic compounds. Intriguingly, the closely related methyltransferases DnrK and RdmB have markedly divergent functions. We investigated their catalytic activities and discovered a previously unknown 10-hydroxylation activity for DnrK and 4-*O*-methylation activity for RdmB. Isotope-labeling demonstrated that the 10-hydroxy group introduced by DnrK is derived from water molecules while RdmB utilizes O_2_. A single residue, E299, was the key modulator in the differing catalytic functions of DnrK and RdmB, especially the decarboxylative oxidation activity. Furthermore, the multifunctionality of DnrK was demonstrated to be SAM-tunable and concerted, whereas RdmB activity was cofactor-dependent and stepwise. Our findings expand the versatility and importance of methyltransferases and should aid studies to enrich the structural diversity and bioactivities of anthracyclines.

## Introduction

Anthracycline-type natural products are used in cancer treatment.^[1, 2, 3, 4]^ Unfortunately, side effects hamper the clinical application of anthracycline-type drugs.^[5]^ Thus, there is considerable interest in synthesizing chemically modified anthracyclines with an improved therapeutic index.^[6]^ Anthracyclines are derived from type II polyketide synthases and feature a tetracyclic 7,8,9,10-tetrahydro-5,12-naphtacenoquinone scaffold (Figure S1).^[1a, 7]^ Tailoring enzymes, including hydroxylases, carboxylases, methyltransferases, and oxidases, contribute to the structural complexity.^[1a, 6]^ However, many of the post-modification enzymes can catalyze unconventional reactions, adding to anthracycline diversity.^[8]^ Two methyltransferase homologues, DnrK and RdmB, have significantly divergent and atypical functions in anthracycline biosynthesis.^[8b, 9]^ DnrK from *Streptomyces peucetius* is a 4-*O*-methyltransferase with flexible substrate specificity, which can catalyze the 4-*O*-methylation and 10-decarboxylation of several anthracycline-type natural products (Figure S2A).^[8b, 10]^ In contrast, RdmB from *Streptomyces purpurascens* has 10-hydroxylase activity and performs a decarboxylative hydroxylation reaction on anthracyclines without methyltransferase activity (Figure S2B).^[8a, 8b, 9, 11]^ Interestingly, DnrK and RdmB share 53.4% amino acid sequence similarity and the hallmark *C*-terminal *S*-adenosyl-L-methionine (SAM-binding) motif GxGxG of methyltransferases (Figure S3).^[12]^ The two enzymes have 3D structural similarities with an RMSD of 1.14 Å and a Class I Rossmann-like fold highly similar to that of classical SAM-dependent methyltransferases (Figure S4).^[12, 13]^

In investigating the basis for the divergent functions of DnrK and RdmB, we discovered for the first time of the 10-hydroxylation activity of DnrK and the 4-*O*-methylation activity of RdmB. DnrK catalyzed the concerted methylation, decarboxylation, and hydroxylation reactions in a SAM-tunable manner whereas RdmB catalyzed methylation and decarboxylative hydroxylation in a cofactor-dependent and stepwise mechanism. Notably, the 10-hydroxy group introduced by DnrK was derived from H_2_O while RdmB utilized O_2_. Mutation of E299 of DnrK switched the catalytic characteristics of these two methyltransferases.

## Results and Discussion

### Investigation of DnrK catalytic activity

To study DnrK functions, the DnrK-encoding gene from the doxorubicin producer *Streptomyces coeruleorubidus* was cloned into pET28a (Figure S5),^[14]^ and soluble, recombinant *N*-terminal His^6^-tagged DnrK was obtained in *Escherichia coli* (Figure S6). DnrK-catalyzed reactions were conducted *in vitro* in the presence of SAM using the natural substrate 10-carboxy-13-deoxycarminomycin (**1**) (Figure S7 and Figures S30-S34). As expected, the 10-decarboxylated product 13-deoxycarminomycin (**2**) (*m/z* = 500.1921, [M+H]^+^, *calcd*. 500.1915) and the 10-decarboxylated and 4-*O*-methylated product 13-deoxydaunorubicin (**3**) (*m/z* = 514.2078, [M+H]^+^, *calcd*. 514.2072) were observed by ultra-performance liquid chromatography (UPLC) (Figure 1, 2A-i and ii); high-resolution electrospray ionization-liquid chromatography mass spectrometry (HRESI-LCMS) (Figure S8); and confirmed by nuclear magnetic resonance (NMR) analysis (Figure S9 and Figures S35-S44). Unexpectedly, a new product with a molecular mass of 530.2028 (*m/z*, [M+H]^+^, *calcd*. 530.2021) was also detected by HRESI-LCMS (Figure S8), indicating generation of an oxidized product. This product was purified and identified as 10-hydroxy-13-deoxydaunorubicin (**4**) by 1D and 2D NMR analysis (Figures S9 and S45-49). However, when **2** and **3** were used as DnrK substrates, the corresponding C-10 hydroxylated product was not detected by UPLC (Figure S10), suggesting that the inert CH_2_ group cannot be activated by DnrK. Thus, the 10-hydroxylation reaction likely occurs during the 10-decarboxylation reaction within the active site (Figure 1). The discovery of **4** demonstrated that DnrK naturally possesses C-10 hydroxylation activity.

**Figure 1.**
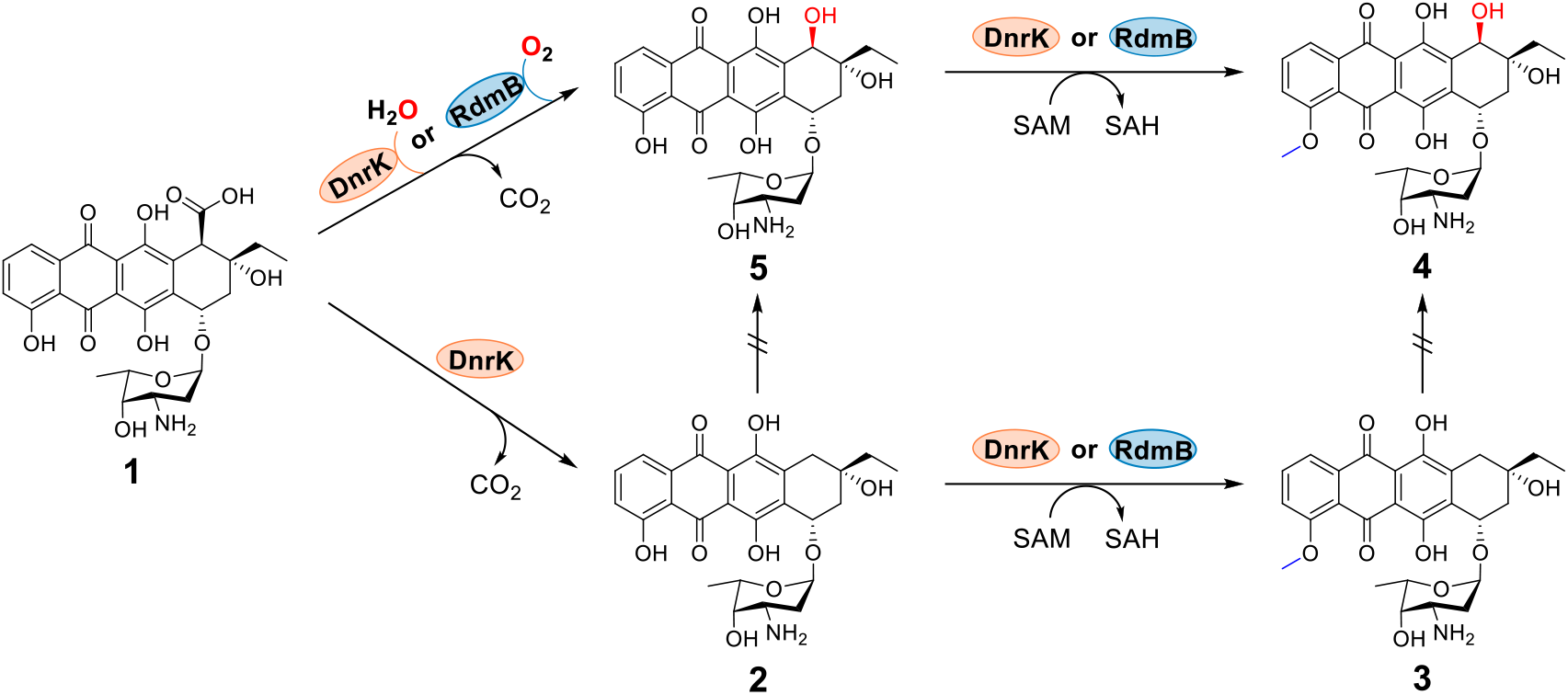
Reactions mediated by DnrK and RdmB. Substrates and catalytic products: 10-carboxy-13-deoxycarminomycin (**1**), 13-deoxycarminomycin (**2**), 13-deoxydaunorubicin (**3**), 10-hydroxy-13-deoxydaunorubicin (**4**), and 10-hydroxy-13-deoxycarminomycin (**5**).

**Figure 2.**
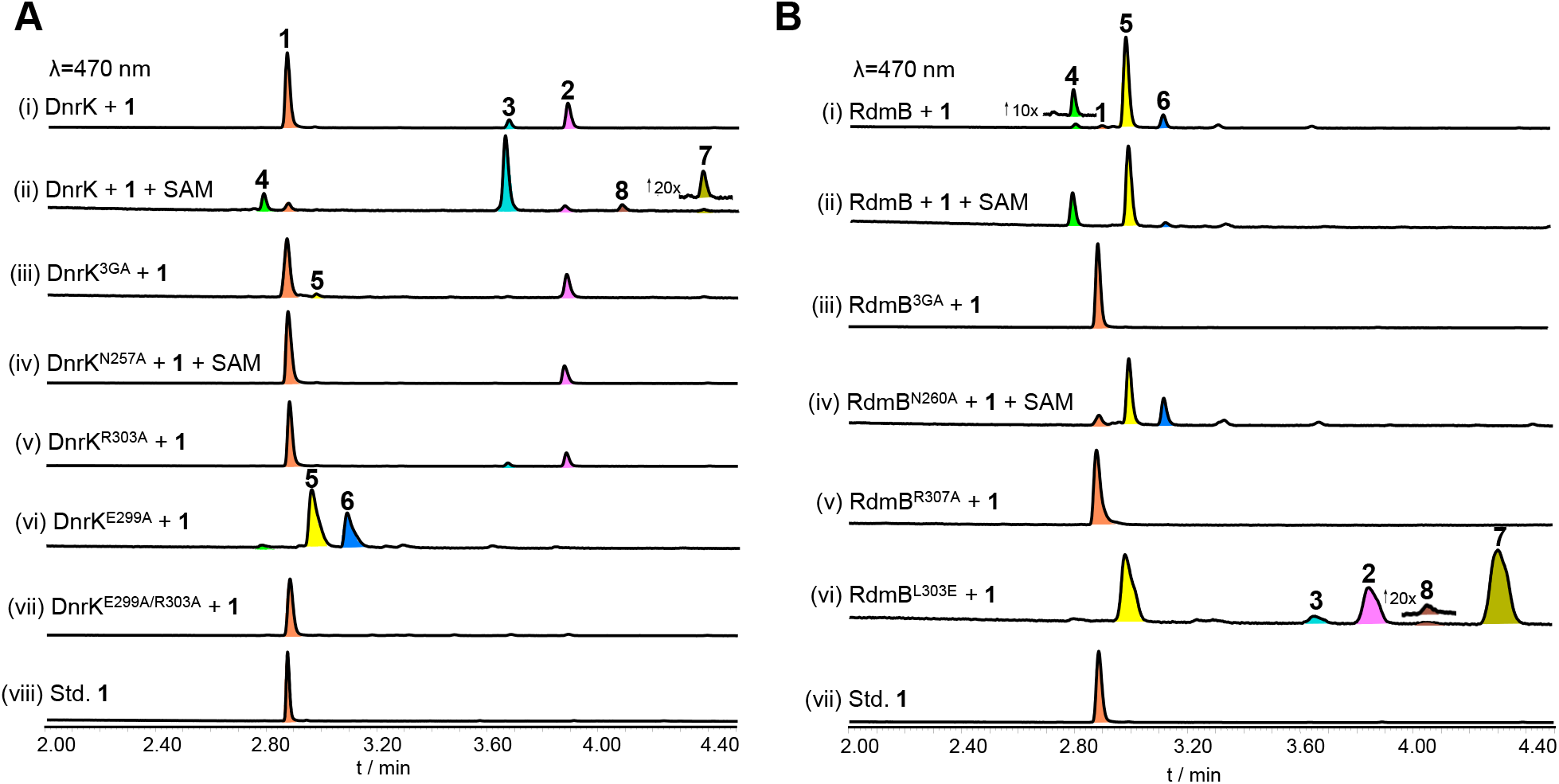
Comparative analysis of the enzymatic conversion of **1** catalyzed by DnrK and RdmB *in vitro*. (A) UPLC analysis of the conversion of **1** catalyzed by DnrK and its mutants DnrK^3GA^, DnrK^N257A^, DnrK^R303A^, DnrK^E299A^ and DnrK^E299A/R303A^. (i-vii) The conversion of **1** by (i) DnrK without additional cofactor SAM; (ii) DnrK in the presence of 1 mM SAM; (iii) DnrK^3GA^ in the presence of 1 mM SAM; (iv) DnrK^N257A^ in the presence of 1 mM SAM; (v) DnrK^R303A^; (vi) DnrK^E299A^; (vii) DnrK^E299A/R303A^; (viii) Std. **1**: The standard of **1**. (B) UPLC analysis of the conversion of **1** catalyzed by RdmB and mutants RdmB^3GA^, RdmB^N260A^, RdmB^R307A^ and RdmB^L303E^. (i-vi) The conversion of **1** by (i) RdmB without additional cofactor SAM; (ii) RdmB in the presence of 1 mM SAM; (iii) RdmB^3GA^ in the presence of 1 mM SAM; (iv) RdmB^N260A^ in the presence of 1 mM SAM; (v) RdmB^R307A^; (vi) RdmB^L303E^; (vii) Std. **1**: The standard of **1**.

### Functional comparison of DnrK and RdmB

Previous studies revealed that RdmB and DnrK can recognize 15-demethoxy-aclacinomycin T as substrate (Figure S2), with RdmB reported to catalyze only the SAM-dependent hydroxylation of the C-10 while DnrK catalyzes C-10 decarboxylation and C-4 methylation.^[8b, 9]^ We discovered that DnrK can also perform C-10 hydroxylation. To more fully compare these two protein homologs, RdmB was expressed as a soluble *N*-terminal His^6^-tagged protein (Figure S6). When RdmB was incubated with **1** in the presence of SAM for 2 hours at 30 °C, the expected 10-hydroxylation product 10-hydroxy-13-deoxycarminomycin (**5**, [M+H]^+^: 516.1873, *calcd*. 516.1864) was detected by UPLC (Figure 1 and Figure 2B-i and ii) and HRESI-LCMS (Figures S8); **5** was then purified from a large-scale reaction and confirmed by 1D and 2D NMR analysis (Figures S9 and S50-54). Interestingly, a product with the same retention time and *m/z* value to those of **4** was detected by UPLC and HRESI-LCMS (Figure 2B-i and ii, and Figure S8). RdmB converted **5** and **2** to, respectively, the 4-*O*-methylated product **4** and product **3** with a high conversion rate (Figure S11). These results clearly demonstrated that RdmB is able to catalyze the methylation reaction as a conventional SAM-dependent *O*-methyltransferase, in addition to its hydroxylation activity when the daunosamine glycosides of aklavinone (**1, 2** or **5**) were used as substrate.

Time-dependent production of **5** and **4** was discovered during time-course analysis of RdmB with **1** in the presence of SAM. Specifically, conversion of **1** to **6** followed by the production of **5** was accomplished in 5 min with more than 90% conversion rate by RdmB. With longer reaction time, conversion of **6** to **5** was observed. However, no methylated product **4** was detected until the reaction time was extended to 60 min (Figure S12). Compound **6** was deduced to be a peroxide intermediate according to its *m/z* value (532.1822 [M+H]^+^, *calcd*. 532.1813), which is 16 Da larger than that of **5** (Figure S8). Production of **5** and **6** is consistent with the previously reported function of RdmB. ^[11b]^ These results indicated that RdmB performs the two consecutive reactions in a stepwise mechanism, with the decarboxylative hydroxylation of **1** that produces the peroxide intermediate **6** occurring first, and then **6** undergoing homolytic cleavage of the O-O bond generating **5**, followed by the 4-*O*-methylation reaction forming **4**. DnrK had a different catalytic timing during production of the decarboxylation and methylation products, which suggested a concerted pattern since the products were delivered without an apparent sequence of reactions (Figure S13).

When the triglycosylated aclacinomycin A with the C-10 alkoxycarbonyl group was used as substrate, no methylated product of DnrK and RdmB was detected, consistent with previous studies (Figure S14).^[8b, 9]^ To further determine whether the presence of the C-10 alkoxycarbonyl or the triglycosyl group affects methylation activity, the monoglycosylated substrate rhodomycin D was incubated with DnrK or RdmB. Production of methylated rhodomycin D from these two reactions (Figure S15) indicated that decarboxylation of the C-10 alkoxycarbonyl group is not a necessary step and that the triglycosyl group of anthracyclines prevents the methylation reaction.

### Role of SAM for DnrK and RdmB

Methyl transfer is one of many biochemical processes requiring SAM as cofactor, and the positive charge of SAM may also be involved in the delocalization of electrons into the anthraquinone core of the substrate.^[11b, 15]^ When the methyl donor SAM was not added, the decarboxylated **2** was the main product of DnrK from substrate **1** with minor production of methylated **3** (Figure 2A-i). When the DnrK concentration was doubled to 4 μM, a new product was detected in small amounts, which had the same *m/z* value as that of **5** (*m/z* 516.1873 [M+H]^+^, Figure S16). Additionally, SAM omission from RdmB reaction mixtures led mainly to the production of **5** with a small ratio of the methylated product **4** (Figure 2B-i). We reasoned that the small yield of **3** from **2** and production of **4** from **5** may be due to the co-purification of SAM with DnrK and RdmB, which was demonstrated by the detection of SAM in the supernatant of heat-denatured DnrK and RdmB by HRESI-LCMS analysis (Figure S17).^[16]^

Accordingly, the production of non-methylated products **2** and **5** may be derived from the enzyme-mediated 10-decarboxylation and hydroxylation in a SAM-independent manner. To evaluate this hypothesis, the three conserved glycine residues in the SAM-binding motif of DnrK and RdmB were mutated to alanine by site-directed mutagenesis. When **1** was reacted with the resultant mutant protein DnrK^3GA^, **1** was converted to **2** with a time-dependent accumulation (Figure 2A-iii and S18). Thus, unlike known methyltransferases such as TleD,^[17]^ LepI ^[18]^ and TnmJ ^[19]^ that require SAM as a cofactor to catalyze non-methylation reactions, the decarboxylation and hydroxylation activities of DnrK are not necessarily dependent on the presence of SAM. Thus, DnrK can convert **1** into four products, including the C-10 decarboxylated product **2**, the C-10 hydroxylated product **5**, and the respective methylated products **3** and **4**. The 10-hydroxylation products **4** and **5** were also identified in *S. coeruleorubidus* cultures by HRESI-LCMS analysis, albeit in low abundance (Figure S19), indicating these two products are natural intermediates during the biosynthesis of doxorubicin-type anthracyclines.

However, the mutated SAM-free protein RdmB^3GA^ completely abolished both the methylation and hydroxylation activities with **1** (Figure 2B-iii). Furthermore, when the RdmB enzymatic assays were performed in the presence of the SAM competitive inhibitor *S*-adenosyl-_L_-homocysteine (SAH, 200 μM) and sinefungin (200 μM),^[16, 20]^ the conversion rate of **1** into the decarboxylated product was affected slightly (Figure 3A). In particular, the initial substrate conversion rate of **1** by RdmB was about 60% and 50% at 2 min when SAH or sinefungin was added, respectively, to the reaction mixture compared to the rate in the presence of SAM. Over time, substrate **1** was consumed by about 90% at 10 min and completed at 20 min (Figure 3A and S20). These results indicate that the presence of SAM or its analogues helps maintain the structural integrity of RdmB. In contrast, when SAH or sinefungin was incubated with DnrK, the consumption rate of **1** and the production of **2** and **5** were only slightly affected when compared to that of DnrK without addition of SAM or the DnrK^3GA^ mutant (Figure 3A and S21).

**Figure 3.**
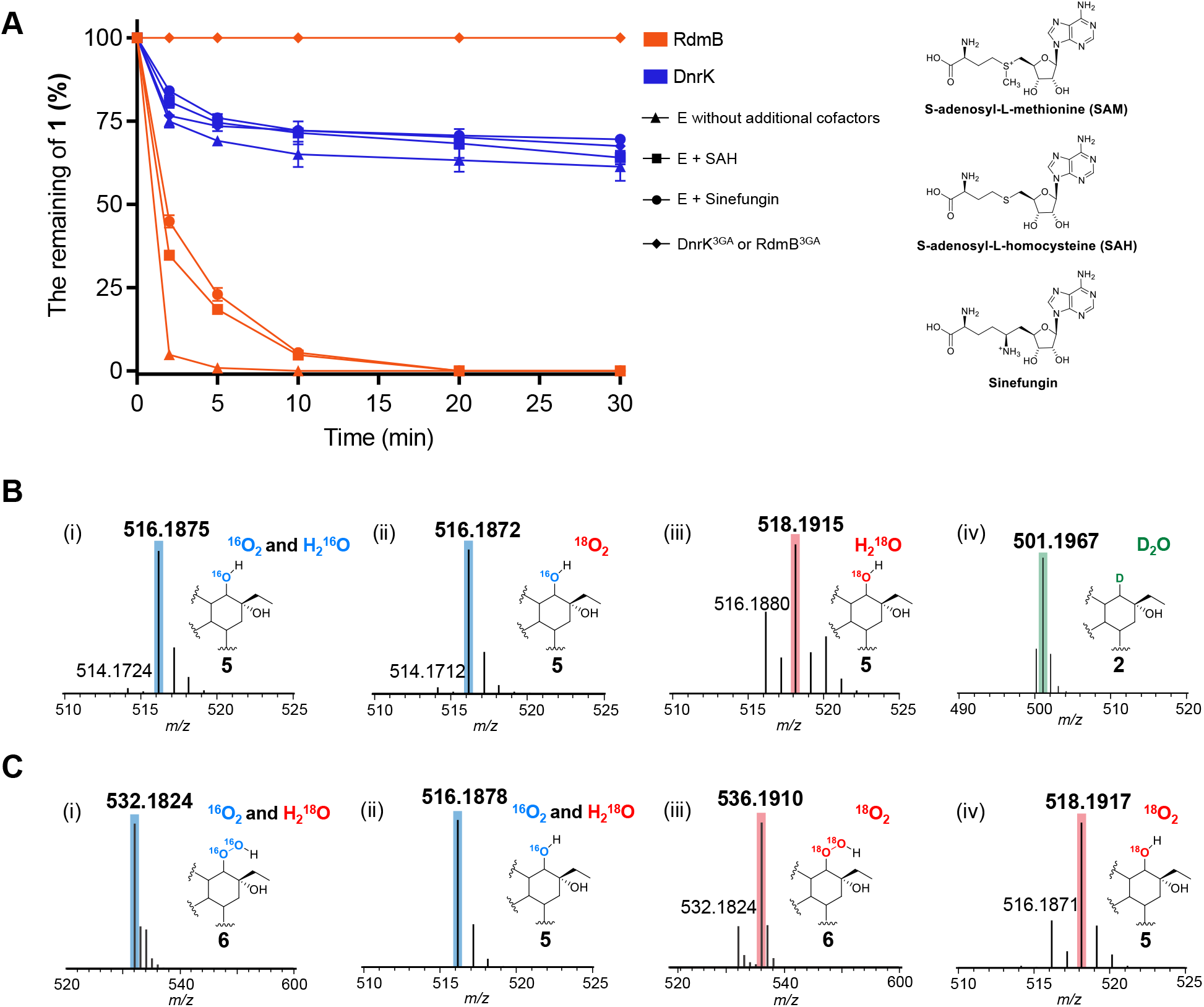
The effect of SAM analogs on the catalytic reactions and isotopic-labeling analysis of DnrK and RdmB (A) Time-course analysis of the consumption of **1** (50 μM) in the presence of enzyme (1 μM) with or without cofactors: SAM (200 μM), SAH (200 μM), sinefungin (200 μM). E, wild-type enzyme DnrK (blue) or RdmB (orange). The data show one representative experiment from at least three independent replicates. (B, C) Isotopic-labeling analysis of DnrK and RdmB mediated reactions toward **1**. (B) HRESI-LCMS analysis of **5** produced by DnrK in a mixture of (i) ^16^O_2_ and H_2_^16^O, (ii) ^18^O_2_ and H_2_^16^O, (iii) H_2_^18^O and (iv) HRESI-LCMS analysis of **2** in D_2_O. (C) HRESI-LCMS of **5** and **6** delivered by RdmB in a mixture of ^16^O_2_ and H_2_^18^O (i and ii), ^18^O_2_ and H_2_^16^O (iii and iv).

These results confirmed that the methylation and hydroxylation activities of RdmB are strictly dependent on the binding of cofactors. Given that the catalytic activities of RdmB^3GA^ were completely abolished, SAM and its analogs are considered an essential structural ligand to maintain ternary structural integrity and the proper binding mode and orientation of electron-rich substrates during decarboxylative hydroxylation of C-10 by RdmB. In contrast, DnrK acts as a member of the methyltransferase superfamily to mediate a SAM-independent decarboxylation and hydroxylation reactions, possibly in a substrate-assisted catalytic mechanism.

### Oxygen origin for the C-10 hydroxy group

To probe the origin of the oxygen atom in the C-10 hydroxy group introduced by DnrK and RdmB, we conducted isotope-labeling experiments under ^18^O_2_ atmosphere. The molecular mass of **5** was not affected in HRESI-LCMS analysis of the DnrK-mediated reactions (Figure 3B-i and ii). However, when the reaction was performed in H_2_^18^O buffer, the molecular mass of **5** was 2 Da more than that observed in H_2_^16^O buffer (518.1915 *vs* 516.1864, [M+H]^+^) (Figure 3B-i and iii). Thus, the C-10 hydroxyl of **5** introduced by DnrK originated from water molecules rather than O_2_. Conversion of **1** by DnrK with D_2_O revealed the molecular mass of **2** was 1 Da larger than that in normal water (m/z = 501.1967 *vs* 500.1978, [M+H]^+^) (Figure 3B-iv).

When the RdmB-mediated reaction was performed in ^18^O_2_, the molecular mass of **6** and **5** was increased by 4 Da and 2 Da, respectively (Figure 3C-iii and iv). With H_2_^18^O buffer, the molecular mass of **5** or **6** was not affected in HRESI-LCMS analysis (Figure 3C-i and ii). Thus, the oxygen atoms of **5** and **6** originated from O_2_, and **6** is converted to **5** during the reaction in the absence of an exogenous reducing agent, consistent with the previously proposed mechanism of a cofactor-less oxygenase that consecutively utilizes electron-rich substrates to activate O_2_.^[11b, 18]^ Accordingly, the 10-decarboxylative hydroxylase mechanism of DnrK potentially differs from that of RdmB.

### Methyl transfer mechanisms of DnrK and RdmB

To accurately compare the catalytic mechanisms of the two enzymes, the three-dimensional crystal structure with a natural substrate is required, but has not been solved, although crystal structures with a non-natural substrate have been reported. ^[8b, 9, 12b]^ Accordingly, DnrK and RdmB were crystallized with **1**. The X-ray crystal structures of DnrK-SAH-**3** and RdmB-SAH-**3** were solved as homodimers at 1.3 Å and 2.2 Å resolution, respectively (PDB IDs: 8KHI and 8KHJ), indicating that the decarboxylation reaction has proceeded during crystallization (Figure 4A and 4B, Table S3). The overall 3D structures of DnrK and RdmB were exceedingly similar, with an RMSD of 1.0 Å over 295 equivalent Cα atoms and similar SAH-product positioning (Figure S22). Each subunit of the single ternary structure displayed the typical fold of Class I methyltransferases, consisting of an *N*-terminal domain with a mainly helical structure for substrate recognition and dimerization and a *C*-terminus containing a Rossman-like fold for substrate binding and catalysis.^[21]^

**Figure 4.**
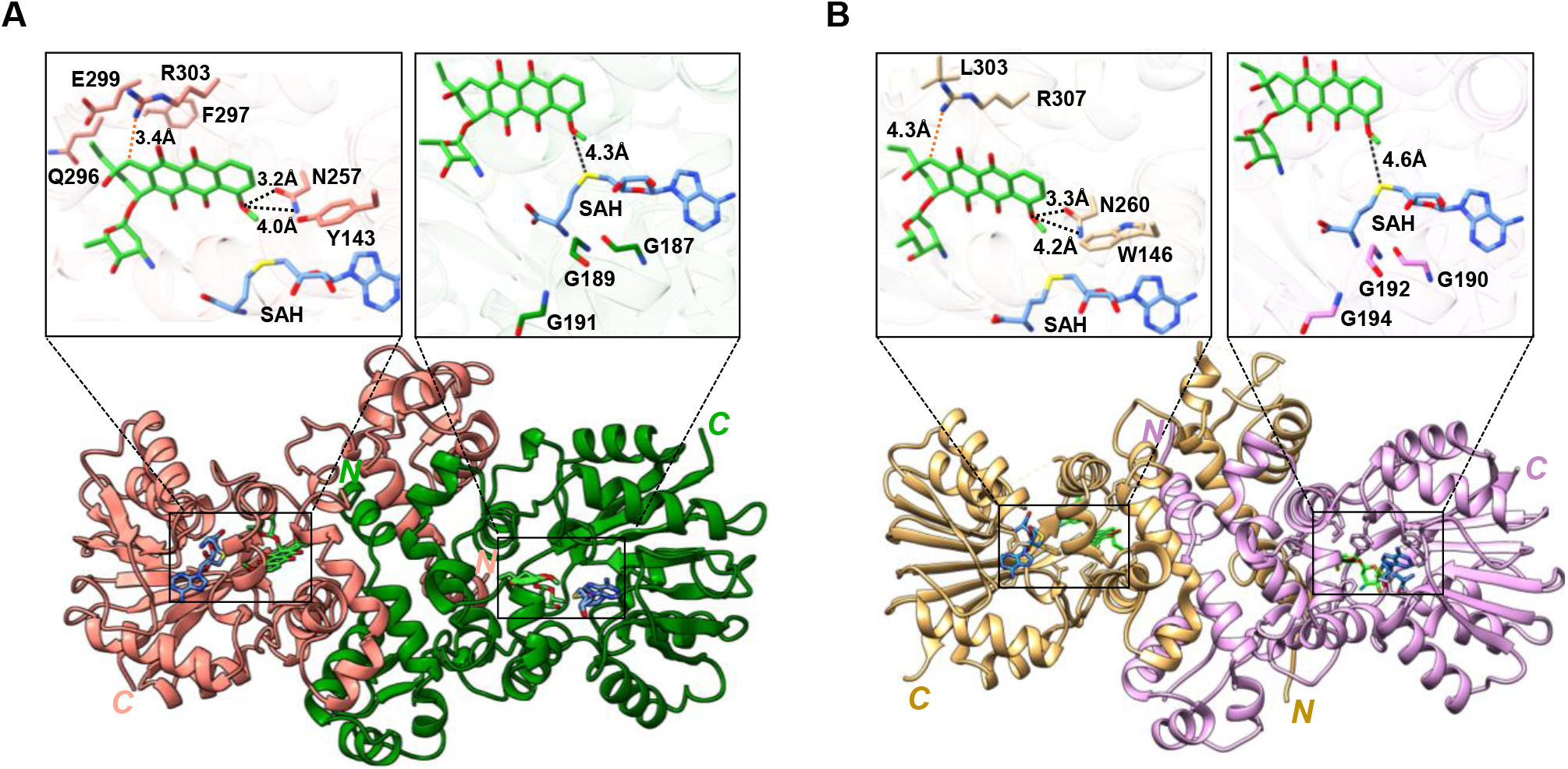
Ternary structure complexes of DnrK and RdmB with functional verification by sited-directed mutagenesis. (A) 3D structure of DnrK bound with product **3** and SAH. The two monomers of DnrK are colored in pink and cyan. The bound ligands of SAH and **3** are shown in the stick models by blue and green. The top box shows a zoom-in view of the active site of DnrK with **3**. (B) 3D structure of RdmB bound with product **3** and SAH. The two monomers of RdmB are colored in tan and lilac. The top box shows a zoom-in view of the active site of RdmB with **3**.

Specifically, product **3** was found in the catalytic pocket, and SAH was bound to the consensus GxGxG located in a loop connecting the first *β*-sheet and the *α*-helix in the Rossmann fold motif, close to **3** (Figure 4A and 4B). RdmB showed a distance of 4.6 Å between the C4 oxygen atom of **3** and the sulfur atom of SAH, while DnrK showed 4.3 Å in the ternary complex structures (Figure 4A and 4B, top). This distance range is suitable for hydrogen abstraction to form an *O*-methyl group.^[15]^ When substrate **1** was docked into the catalytic sites of DnrK and RdmB by AutoDock Vina,^[22]^ two hydrogen bonds were formed by DnrK^N257^ and DnrK^Y143^ to the C4-hydroxyl group of **1**, while only one hydrogen bond formed between RdmB^N260^ and the C4-hydroxyl group (Figure S23), which may play important roles in the methylation activity. When DnrK^Y143A^ was incubated with **2** in the presence of SAM, little effect was found on the 4-methylation activity of DnrK as indicated by the successful formation of **3** (Figure S24). However, the methylation activity of DnrK^N257A^ or RdmB^N260A^ was completely abolished toward **2** in the presence of SAM (Figure S25). With compound **1** as substrate, the production of decarboxylated **2** and hydroxylated **5**, but not the methylated products, was detected with DnrK^N257A^ or RdmB^N260A^ (Figure 2A-iv and 2B-iv). Accordingly, instead of the previously proposed “proximity and desolvation” mechanism ^[12b]^, an Asn-mediated S_N_2-like type methylation is suggested for both DnrK and RdmB, using an active N257 as a catalytic residue to deprotonate the C4 hydroxyl group of **1**. Deprotonation further enhances the nucleophilicity of the hydroxyl group to attack the SAM donor methyl group and contribute to rate acceleration of methylation. Our results also suggest that the C-7 triglycosylated substrate (*i*.*e*., aclacinomycin A) prevents the proper binding mode needed to accommodate the potential proton extraction of the C4 hydroxyl group from the conserved Asn residue of DnrK and RdmB, preventing methyl transfer activity.

### Switching catalytic activities between DnrK and RdmB

Additionally, we conducted mutation studies of key residues involved in DnrK and RdmB decarboxylation and hydroxylation activities. The ternary complex structures of DnrK and RdmB revealed formation of a salt bridge between the side chains of DnrK R303 and RdmB R307 with the carbonyl group of **1** by substrate docking analysis (Figure S23). R307 of RdmB and R303 of DnrK were proposed to be important for initiating the decarboxylation reaction. ^[11b, 12b]^ As expected, no conversion of **1** was detected with RdmB^R307A^ in the presence of SAM after 2 h (Figure 2B-v). Unexpectedly, **2** and **3** were still detected when **1** was incubated with DnrK^R303A^ (Figure 2A-v), indicating that DnrK R303 is not the key residue for the decarboxylation reaction, therefore differing from RdmB R307.

Considering hydrophilicity and potential interaction, E299, which has a distance of 5.8 Å to the carboxyl group of **1**, was selected as a candidate (Figure S26A and S26B) and mutated to generate DnrK^E299A^ (Figure S27). Importantly, incubation of DnrK^E299A^ and **1** abolished **2** and **3** productions (Figure 2A-vi). Unexpectedly, complete conversion of **1** into **5** and **6** by 1h was detected by UPLC and HRESI-LCMS (Figure 2-vi). Additionally, the oxygen atom in the C-10 hydroxy group of **5** generated by DnrK^E299A^ originated from O_2_ instead of the water origin for DnrK^WT^ (Figure S28B-i). As sequence alignments and structure analysis indicated that E299 was substituted by the nonpolar L303 in RdmB (Figure S26C), DnrK^E299L^ was constructed, which had the same catalytic characteristics as DnrK^E299A^ (Figure S28A-i and B-ii). Interestingly, when E was replaced with D, DnrK catalytic performance was not affected (Figure S28A-ii, and B-iii). However, incubation of DnrK^E299Q^ and DnrK^E299K^ with **1** produced **5** (Figure S28A-iii and iv), and the C-10 hydroxy group oxygen of **5** was derived both from oxygen and water (Figure S28B-iv and v). These site-directed mutagenesis studies indicated that the residue at position 299 of DnrK is vital for controlling catalytic functions.

For comparison, the corresponding L303 in RdmB was mutated into amino acid E. Interestingly, **2** was detected when RdmB^L303E^ was incubated with **1**, together with **5** and **6** (Figure 2B-vi), and the oxygen atom in the C-10 hydroxy group of **5** was derived from O_2_ and water molecules (Figure S28B-vi). Additionally, two more products, **7** and **8**, were detected and deduced to be anthracycline derivatives by UV characteristic spectrum comparisons (Figure S29A). HRESI-LCMS suggested that the structure of **7** (*m/z* = 482.1807, [M+H]^+^, *calcd*. 482.1809) is formed by a C9-C10 elimination reaction from **2** and that **8** (*m/z* = 496.1957, [M+H]^+^, *calcd*. 496.1966) is the methylated product of **7** (Figure S29B). Notably, **7** and **8** were identified by detailed examination of the product profile of DnrK, albeit in low yields (Figure 2A-ii). Formation of the 9,10-elimination products possibly resulted from interaction of DnrK^R285^/RdmB^R288^ with the C9 hydroxyl group of the carbanion intermediate in the hydrophilic pocket conferred by DnrK^E299-R303^/RdmB^E303-R307^ further contributing to the generation of a better leaving group.^[9]^

Our analyses indicate that DnrK^E299^/RdmB^L303^ play important roles in the catalytic function of these two atypical methyltransferases. The hydrophilic residue E is the key modulator, controlling the oxygen origin of the installed 10-hydroxyl group. When E299 and R303 are both close to the substrate carboxyl group, they help coordinate the water molecules and form a polar environment to facilitate the water-mediated oxidation by DnrK. When E299 is replaced by nonpolar residue, as observed with RdmB^L303^, a bulkier hydrophobic environment is formed, allowing O_2_ greater access to the reaction center. Since the decarboxylation is considered the initial step for further oxidation, the double mutant DnrK^R303A/E299A^ was constructed, and the resultant protein completely lost decarboxylation activity toward **1** (Figure 2A-vii). Based on these results, E299 and R303 in DnrK are both involved in the C-10 decarboxylation, and one of them is sufficient to initiate this reaction.

Accordingly, substrate-assisted activation mechanisms of DnrK and RdmB utilizing a highly-conjugated electron-rich tetracyclic 7,8,9,10-tetrahydro-5,12-naphtacenoquinone moiety were proposed.^[15, 23]^ After **1** binds to the RdmB catalytic pocket, decarboxylation of **1** is initiated by R307, resulting in the formation of the carbanion intermediate **9**. Subsequently, substrate-assisted activation of O_2_ oxidizes **9** to yield the hydroperoxide intermediate **6**, which is then converted into **5** by spontaneous homolytic O-O bond cleavage. Exclusion of water molecule from the active site is important for this substrate-assisted oxidation step, possibly because the substrate-binding pocket is formed by hydrophobic residues as demonstrated by mutagenesis of L303 (Figure S26). Contrastingly, after the decarboxylation reaction of DnrK mediated by E299 and R303, the C-10 carbanion of intermediate **9** is quenched by protonation of water molecules to deliver **2** as the main product. Alternatively, water molecules could attack intermediate **10** to produce**11**, which undergoes dehydrogenative oxidation, possibly due to the double annelation effect, to form **5** (Figure 5).

**Figure 5.**
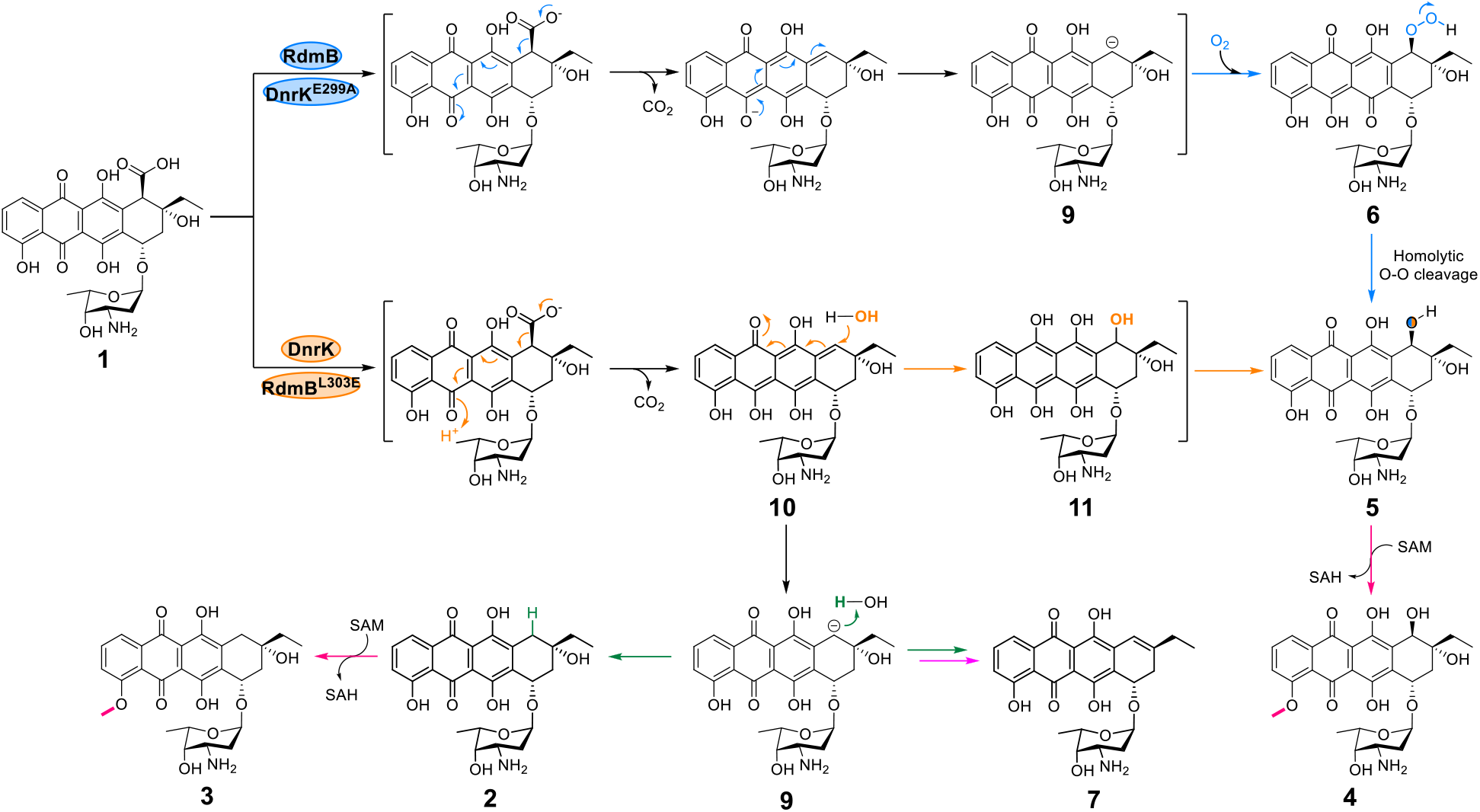
Proposed catalytic mechanisms mediated by DnrK and RdmB toward substrate **1**. Product **5** is derived by RdmB from homolytic O-O bond cleavage of the peroxyl intermediate **6** since no reductant was added into the reaction mixture, while a dehydrogenative oxidation reaction takes place due to the double annelation effect of intermediate **10** with DnrK. The methylation activity of DnrK and RdmB shows flexible substrate specificity.

## Conclusion

We demonstrated the inherent 10-hydroxylation activity of DnrK and 4-*O*-methylation activity of RdmB in anthracycline biosynthesis. Based on our comparative analysis, a new Asn-mediated acid/base catalyzed S_N_2-type nucleophilic substitution methylation by DnrK and RdmB was proposed instead of the previously reoprted “proximity and desolvation” mechanism.^[12b]^ Additionally, DnrK and RdmB decarboxylation activity occurs via different catalytic residues, with the basic R307 as the key residue in RdmB, whereas R303 and the acidic E299 are both involved in DnrK. Despite functionally convergent evolution, DnrK activity is SAM-tunable and concerted, while RdmB activity is cofactor dependent and stepwise. Furthermore, the 10-OH group introduced by RdmB originates from O_2_, while that installed by DnrK originates from water molecules. To our knowledge, DnrK is the first example of a cofactor-less oxygenase in the methyltransferase family that uses water molecules for hydroxylation activity. Intriguingly, E299 was found to be the key switch controlling the catalytic function of DnrK and RdmB, as demonstrated by the gain of RdmB functions by the mutant DnrK^E299A^.

Several SAM-dependent methyltransferases can catalyze diverse types of non-methylating reactions.^[17, 18, 24]^ Interestingly, methyltransferase TnmJ, involved in anthraquinone-fused enediyne biosynthesis, functions as a cofactor-less oxygenase in catalyzing deformylation and epoxidation reactions.^[19]^ Our functional characterization of DnrK and RdmB expands our understanding of the versatility and potential of cofactor-less methyltransferases.

## Methods

### Materials

10-carboxy-13-deoxycarminomycin (**1**) was isolated from *Streptomyces coeruleorubidus* mutant *S. coeruleorubidus::*ΔDnrK and compounds **2** -**5** were obtained through enzymatic reactions. SAM (*S*-adenosyl-L-methionine) and SAH (*S*-adenosine-L-homocysteine) were purchased from Aladdin (Shanghai, China), and sinefungin was purchased from Psaitong (Beijing, China). All antibiotics used in this study were obtained from Solarbio (Beijing, China). All restriction enzymes were purchased from Thermo Scientific (PA, USA). 2 × Phanta Max Master Mix Vazyme (Nanjing, China) was employed to amplify DNA fragments. Kits for plasmid mini-preparations and DNA gel extractions were acquired from Omega Bio-tek, Inc. (GA, USA). Ni-NTA SefinoseTM Resin (Settled Resin) for protein purification was purchased from Sangon Biotech (Shanghai, China). The FlexiRun premixed gel solution for SDS-PAGE was obtained from MDBio (Qingdao, China). Oligonucleotide primers and DNA sequencing were ordered from TsingKe Biotech (Shanghai, China). Gene synthesis was ordered from Beijing Genomics Institution (Shenzhen, China). Organic solvents for compound isolation and purification were bought from sinopharm (Shanghai, China).

### Analytical procedures

The protein sequence alignments were performed using T-Coffee^[25]^ and ESPript3.^[26]^ The biosynthetic gene clusters of *S. coeruleorubidus* were predicted using the bacterial version of antiSMASH (https://secondarymetabolites.org/).^[27]^ NMR data were processed using MestReNova 9.0. All HPLC (DIONEX Ultimate 3000 instrument) analyses were performed on a YMC Triart C18 column (4.6 × 250 mm, 5 μm, UV detection at 470 nm) with a biphasic solvent system of acetonitrile-0.1% formic acid (solvent B) and water (solvent A) at a flow rate of 1.0 mL/min. All UPLC (Acquit-H-class, Waters) analyses were performed using a YMC Triart C18 (2.1 × 100 mm, 1.9 μm, UV detection at 470 nm) column with a biphasic solvent system of acetonitrile-0.1% formic acid (solvent B) and water (solvent A) at a flow rate of 0.4 mL/min. HRESI-LCMS analyses were carried out on a Bruker impact HD High Resolution Q-TOF mass spectrometer.

### Strain and Culture Conditions

*S. coeruleorubidus* and mutant were grown in NDYE liquid medium (maltose 22.5 g, yeast extract 5.04 g, NaNO_3_ 4.28 g, K_2_HPO_4_ 0.23 g, HEPES 4.77 g, MgSO_4_·7H_2_O 0.12 g, NaOH 0.4 g, 1 L; adding trace element solution: ZnCl_2_ 40 mg, FeCl_3_·6H_2_O 200 mg, CuCl_2_·2H_2_O 10 mg, MnCl_2_·4H_2_O 10 mg, Na_2_B_4_O_7_·10H_2_O 10 mg, (NH_4_)_6_Mo_7_O_24_·4H_2_O 10 mg, 1 L) or on MS agar (soybean flour 20 g, mannitol 20 g and agar 20 g, 1 L) plates at 30 °C for 5 days. *Escherichia coli* DH5α strain was used for vector construction and plasmid preparation. *E. coli* BL21 (DE3) (for *N*-His^6^-tagged proteins) was used for protein expression, and cultured in LB media.

### Construction of gene inactivation mutants

The CRISPR/Cas9 genome editing technology was used for gene *dnrK* disruption. To construct the plasmid for the target gene inactivation, the upstream and downstream homology arms were amplified with dnrK-LF/dnrK-LR primers and dnrK-RF/dnrK-RR primers using genomic DNA of *S. coeruleorubidus* as template. The *dnrK* deletion sgRNA expression cassette was amplified from pKCcas9dO with the dnrK-sgRNA-F/dnrK-sgRNA-R primers. The above three fragments were then ligated to the plasmid pKCcas9dO, which was linearized by SpeI and HindIII using the ClonExpress Ultra One Step Cloning Kit to construct the knockout plasmid pKCcas9d-dnrK. The plasmid was then conjugated into S. *coeruleorubidus* according to the standard procedure.^[28]^ After cultured for 3-5 days at 30 °C, the colonies with apramycin-resistant were transferred to MS plates supplied with apramycin antibiotics at final concentration of 100 μg/mL. The apramycin-sensitive colonies were picked and confirmed by diagnostic PCR analysis using the primers dnrK-VF/dnrK-VR. Wild type *S. coeruleorubidus* st was used as a control and the expected lengths of the PCR products were 2706 bp and 1635 bp, respectively. Subsequently, mutant *S. coeruleorubidus::*ΔDnrK was cultured on MS plates without any antibiotics at 37 °C to lose the CRISPR/Cas9 plasmids.

### Isolation of 10-carboxy-13-deoxycarminomycin (1)

To chracterize the accumulated compounds of the mutant strain *S. coeruleorubidus::*ΔDnrK, a 1 L large-scale fermentation was carried out for the mutant strain ΔDnrK using the fermentation condition described above. After 5 days of cultivation, the fermentation broth was collected and dried under vacuum to obtain crude extracts. The crude extracts were subjected to HPLC for purification **1** using a YMC Triart C18 column (20 mm × 250 mm, 7 μM). Water (solvent A) and acetonitrile-0.1% FA (solvent B) were used as the mobile phases at a flow rate of 15 mL/min. The HPLC program was as follows: 0-2 min, 20% B in A; 2-20 min, 20-80% B in A; 20-20.5 min 80-100% B in A; 20.5-22 min 100% B; 22-22.5 min 100-20% B in A; and 22.5-25 min 20% B in A. The obtained fractions were dried under vacuum to give compounds **1** (63 mg), whose structures were identified by HRESI-LCMS and NMR analysis.

### Construction of plasmids for protein expression

The DNA sequence encoding *dnrK* was amplified from the gDNA of *S. coeruleorubidus* using the primer pair of dnrK-F/dnrK-R. The purified DNA fragments were ligated into *Nde*I*/Hind*III digested pET28a (+) with His^6^ at the N-terminus using the ClonExpress Ultra One Step Cloning Kit, and transformed into *E. coli* DH5α for plasmid amplification. The recombinant plasmid was verified by DNA sequencing, and the correct plasmids were transformed into *E. coli* BL21(DE3) for protein overexpression. The gene sequences of RdmB were synthesized according to the codon preference of *E. coli* BL21(DE3) by Beijing Genomics Institution (Shenzhen, China). The site-specific mutations of DnrK were constructed by site-directed PCR using PET28a-*dnrK* as a template with primer pairs of dnrK^3GA^-F/dnrK^3GA^-R, dnrK^Y143A^-F/dnrK^Y143A^-R, dnrK^N257A^-F/dnrK^N257A^-R, dnrK^R303A^-F/dnrK^R303A^-R, dnrK^E299A^-F/dnrK^E299A^-R, dnrK^E299A/R303A^-F/dnrK^E299A/R303A^-R, dnrK^E299L^-F/dnrK^E299L^-R, dnrK^E299D^-F/dnrK^E299D^-R, dnrK^E299Q^-F/dnrK^E299Q^-R, dnrK^E299K^-F/dnrK^E299K^-R. Similarly, the site-specific mutations of RdmB were constructed by site-directed PCR using PET28a-*rdmB* as template using primer pairs of RdmB^3GA^-F/RdmB^3GA^-R, RdmB^N260A^-F/dnrK^N260A^-R, and RdmB^R307A^-F/RdmB^R307A^-R, RdmB^L303E^-F/RdmB^L303E^-R. The individual amplified product was purified using the Omega Gel Extraction Kit according to the manufacturer’s instructions and then transformed into *E. coli* DH5α for plasmid amplification. The above recombinant plasmids were verified by DNA sequencing and the correct plasmids were transformed into *E. coli* BL21(DE3) for protein overexpression.

### In vitro enzymatic assays and products detection

Unless otherwise stated, all enzymatic assays were carried out in a total volume of 100 μL in the desalting buffer (50 mM NaH_2_PO_4_, 300 mM NaCl, 10% glycerol, pH 7.4) at 30 °C, and the boiled enzymes were used as negative controls. For DnrK catalyzed reaction, 0.1 mM **1** or **2** was incubated with 2 μM DnrK with or without 1 mM SAM at 30 °C for 2 h. For RdmB catalyzed reaction, 0.1 mM **1** or **2** or **5** was incubated with 2 μM RdmB and 1 mM SAM at 30 °C for 2 h. To analyze the activity of DnrK mutants, DnrK was replaced by its mutants (DnrK^3GA^ / DnrK^N257A^ /DnrK^R303A^ / DnrK^Y143A^ / DnrK^W106A^ / DnrK^W106A/R303A^) in the above reaction system. To analyze the activity of RdmB mutants, RdmB was replaced by its mutants (RdmB^3GA^ / RdmB^N260A^ / RdmB^R307A^) in the above reaction system. The reactions were quenched by thorough mixing with 200 μL methanol to precipitate the proteins. The metamorphosed proteins were removed by high-speed centrifugation at 14,000 rpm for 10 min, and the supernatants were analyzed by UPLC or HRESI-LCMS using gradient elution programs. Water-0.1% FA (solvent A) and acetonitrile-0.1% FA (solvent B) were used as the mobile phases. The UPLC program: 0-0.5 min, 20% B in A; 0.5-5 min, 20-90% B in A; 5-5.1 min 90-100% B in A; 5.1-5.6 min 100% B; 5.6-5.7 min 100-20% B in A; and 5.7-6.5 min 20% B in A at a flow rate of 0.4 mL/min, UV 470 nm. HRESI-LCMS program: 0-2 min, 20% B in A; 2-20 min, 20-80% B in A; 20-20.5 min 80-100% B in A; 20.5-22 min 100% B; 22-22.5 min 100-20% B in A; and 22.5-25 min 20% B in A at a flow rate of 1 mL/min, UV 470 nm.

### Time course analysis of the conversion of 1 catalyzed by RdmB and DnrK

For the conversion of **1** by DnrK or RdmB, a 100 μL reaction system containing 100 μM **1** with 2 μM DnrK in desalting buffer was incubated at 30 °C. The time points for the reactions were set to 0, 2, 5, 10, 30, 60, 120 min, respectively. When each reaction time point was reached, the reaction mixture was quenched with 200 μL methanol. After centrifugation at 14,000 rpm for 10 min, the supernatants were analyzed by UPLC, UV 470 nm. To investigate the conversion of **1** by DnrK or RdmB with cofactor SAH or sinefungin, the reaction mixture containing 1 μM DnrK or RdmB and 200 μM SAH or 200 μM sinefungin was incubated at room temperature for 10 min. Then, 50 μM **1** was added to the reaction mixture to initiate the enzymatic reaction (total volume was 300 μL). The time point for the reactions was set at 0, 2, 5, 10, 20, 30 min, respectively. When the reaction time point reached, a 50 μL aliquot of the reaction mixture was taken and quenched by mixing with 100 μL of methanol. After centrifugation at 14,000 rpm for 10 min, the supernatant was subjected to UPLC analysis. The reaction systems catalyzed by DnrK or RdmB without additional cofactors were used as controls, following the same procedure as described above with the addition of cofactors (SAH and sinefungin). The consumption of **1** was estimated based on the standards. Error bars represent the standard deviation of three independent replicates.

### Isolation of compound 2-5

To identify the structure of **2**, an 80 mL total volume large-scale enzymatic reaction was performed in desalting buffer containing 0.25 mM **1** and 10 μM DnrK^3GA^. After incubation at 30 °C for 4 hours, 160 mL of methanol was added to quench the reaction. Compound **2** (2 mg) was purified from the concentrated reaction mixture by semi-preparative HPLC using 50% ACN in H_2_O (0.1% FA) at a flow rate of 3 mL/min.

To identify the structure of **3**, a 50 mL total volume large-scale enzymatic reaction was performed in desalting buffer containing 0.5 mM **1**, 1 mM SAM and 10 μM DnrK. After incubation at 30 °C for 4 hours, 100 mL of methanol was added to quench the reaction. Compound **3** (3 mg) was purified from the concentrated reaction mixture by semi-preparative HPLC using 45% ACN in H_2_O (0.1% FA) at a flow rate of 3 mL/min.

To determine the structure of **4**, a 50 mL total volume large-scale enzymatic reaction was performed in desalting buffer containing 0.5 mM **5**, 10 μM RdmB, and 1 mM SAM. After incubation at 30 °C for 2 hours, 100 mL of methanol was added to quench the reaction. Compound **4** (2 mg) was purified from the concentrated reaction mixture by semi-preparative HPLC using 26% ACN in H_2_O (0.1% FA) at a flow rate of 3 mL/min.

To obtain sufficient of **5**, a 200 mL total volume large-scale enzymatic reaction were performed in desalting buffer containing 1 mM **1** and 5 μM RdmB. After incubation at 30 °C for 2 hours, 400 mL of methanol was added to quench the reaction. Compound **5** (5 mg) was purified from the concentrated reaction mixture by semi-preparative HPLC using 24% ACN in H_2_O (0.1% FA) at a flow rate of 3 mL/min.

### NMR data of the compound 1-5

10-carboxy-13-deoxycarminomycin (**1**): ^1^H NMR (600 MHz, MeOD) δ 7.82 - 7.63 (m, 1H), 7.60 (s, 1H), 7.28 - 7.07 (m, 1H), 5.43 (d, *J* = 9.5 Hz, 1H), 5.02 (d, *J* = 18.5 Hz, 1H), 4.27 (dd, *J* = 31.9, 6.4 Hz, 1H), 4.02 (d, *J* = 14.7 Hz, 1H), 3.65 (dd, *J* = 20.5, 9.9 Hz, 1H), 3.60 - 3.54 (m, 1H), 3.50 (dd, *J* = 11.2, 6.0 Hz, 1H), 2.48 - 2.35 (m, 1H), 2.27 (d, *J* = 13.5 Hz, 1H), 2.06 - 1.88 (m, 2H), 1.31 - 1.27 (m, 3H), 1.23 - 1.17 (m, 1H), 1.14 - 1.03 (m, 3H). ^13^C NMR (151 MHz, MeOD) δ 187.67, 175.42, 168.82, 162.20, 160.31, 156.08, 136.79, 136.57, 133.29, 118.97, 118.81, 115.77, 115.69, 110.76, 110.50, 70.46, 66.56, 65.18, 63.01, 35.51, 34.23, 33.28, 30.51, 15.60, 6.21.

13-deoxycarminomycin (**2**): ^1^H NMR (600 MHz, MeOD) δ 7.84 (s, 1H), 7.78 (s, 1H), 7.33 (s, 1H), 5.48 (s, 1H), 4.30 (dd, *J* = 23.7, 6.3 Hz, 1H), 3.67 (dd, *J* = 11.1, 5.9 Hz, 1H), 3.09 (d, *J* = 17.4 Hz, 1H), 2.67 (d, *J* = 21.3 Hz, 1H), 2.31 (d, *J* = 13.6 Hz, 1H), 2.05 (d, *J* = 7.2 Hz, 1H), 1.93 (s, 1H), 1.70 (dd, *J* = 18.7, 6.9 Hz, 2H), 1.33 (s, 2H), 1.32 (d, *J* = 6.4 Hz, 3H), 1.09 (t, *J* = 6.9 Hz, 3H). ^13^C NMR (151 MHz, MeOD) δ 189.32, 171.16, 165.48, 164.26, 162.26, 136.82, 136.72, 128.31, 126.96, 121.88, 121.18, 120.05, 110.79, 110.01, 99.71, 71.23, 68.92, 66.55, 63.01, 37.69, 35.49, 34.26, 28.22, 15.57, 6.37.

13-deoxydaunorubicin (**3**): ^1^H NMR (600 MHz, MeOD) δ 7.96 (s, 1H), 7.84 (s, 1H), 7.57 (d, *J* = 8.3 Hz, 1H), 4.39 (d, *J* = 9.3 Hz, 1H), 4.25 (d, *J* = 6.7 Hz, 1H), 3.83 (t, *J* = 5.6 Hz, 1H), 3.72 - 3.68 (m, 1H), 3.52 (d, *J* = 6.0 Hz, 3H), 3.15 (t, *J* = 5.6 Hz, 1H), 3.03 (d, *J* = 17.7 Hz, 1H), 2.27 (d, *J* = 14.0 Hz, 1H), 2.06 - 2.01 (m, 1H), 1.88 (dd, *J* = 13.0, 4.6 Hz, 1H), 1.64 (dq, *J* = 14.4, 7.5 Hz, 2H), 1.33 - 1.29 (m, 2H), 1.28 (t, *J* = 3.1 Hz, 3H), 1.05 (t, *J* = 7.3 Hz, 3H). ^13^C NMR (151 MHz, MeOD) δ 189.10, 171.20, 163.26, 160.66, 159.72, 137.96, 137.27, 129.36, 127.68, 122.46, 121.25, 120.98, 113.07, 112.86, 87.83, 71.81, 68.64, 63.29, 62.04, 50.32, 39.94, 36.31, 31.47, 30.19, 17.66, 8.46.

10-hydroxy-13-deoxycarminomycin (**4**): ^1^H NMR (600 MHz, MeOD) δ 8.06 (t, *J* = 11.0 Hz, 1H), 7.88 (t, *J* = 8.0 Hz, 1H), 7.61 (d, *J* = 8.3 Hz, 1H), 5.48 (d, *J* = 3.3 Hz, 1H), 5.07 (d, *J* = 2.8 Hz, 1H), 4.90 (s, 1H), 4.31 - 4.18 (m, 1H), 4.05 (s, 3H), 3.66 - 3.62 (m, 1H), 3.58 (dd, *J* = 11.2, 4.9 Hz, 1H), 3.50 (dd, *J* = 11.2, 6.0 Hz, 1H), 2.19 - 2.15 (m, 2H), 2.07 - 1.98 (m, 1H), 1.89 - 1.83 (m, 1H), 1.81 (dd, *J* = 14.0, 6.5 Hz, 1H), 1.74 (dd, *J* = 14.1, 7.3 Hz, 1H), 1.32 (dd, *J* = 14.3, 7.1 Hz, 2H), 1.28 (s, 3H), 1.09 (t, *J* = 7.4 Hz, 3H). ^13^C NMR (151 MHz, MeOD) δ 187.75, 174.73, 163.32, 162.74, 158.19, 137.56, 136.72, 130.69, 129.34, 126.32, 121.32, 121.06, 114.41, 113.79, 102.39, 74.57, 73.44, 68.82, 68.62, 67.08, 65.13, 57.84, 34.80, 32.34, 30.39, 17.64, 7.59.

10-hydroxy-13-deoxycarminomycin (**5**): ^1^H NMR (600 MHz, MeOD) δ 7.84 (s, 1H), 7.78 (s, 1H), 7.32 (s, 1H), 5.48 (s, 1H), 5.05 (s, 1H), 4.83 (s, 1H), 4.30 - 4.22 (m, 1H), 3.65 (d, *J* = 9.3 Hz, 1H), 3.58 (dd, *J* = 11.2, 4.9 Hz, 1H), 3.50 (dd, *J* = 11.2, 6.0 Hz, 1H), 2.17 (q, *J* = 14.8 Hz, 2H), 2.03 (t, *J* = 11.6 Hz, 1H), 1.89 (d, *J* = 9.2 Hz, 1H), 1.82 (dt, *J* = 14.6, 7.2 Hz, 1H), 1.79 - 1.71 (m, 1H), 1.29 (d, *J* = 6.5 Hz, 3H), 1.09 (t, *J* = 7.4 Hz, 3H). ^13^C NMR (151 MHz, MeOD) δ 188.28, 171.68, 164.49, 163.52, 161.95, 139.17, 135.66, 130.47, 129.37, 126.46, 121.45, 121.20, 113.80, 113.42, 102.32, 74.55, 73.21, 68.66, 68.63, 67.06, 65.10, 34.78, 32.24, 30.29, 17.64, 7.60.

### Confirmation of the presence of SAM in DnrK and RdmB

300 μM DnrK and RdmB in 30 μL storage buffer were denatured by heating at 100 °C for 10 min each. 100 μM SAM in water was heated at 100 °C for 10 min. Then, the solutions were centrifuged at 14,000 rpm for 10 min. The supernatants were analyzed by HRESI-LCMS with a linear gradient of 5-60% ACN-H_2_O with 5 mM ammonium acetate in 25 min at a flow rate of 1 mL/min. UV detection was performed at 260 nm. The same procedures were performed for DnrK^3GA^ and RdmB^3GA^.

### Time-dependent conversion experiments of 1 by DnrK^3GA^

A 500 μL reaction system containing 100 μM of 1 with 2 μM DnrK^3GA^ in desalting buffer was incubated at 30 °C. The time points for the reactions were set at 0, 1, 2, 4, and 6 h, respectively. When each reaction reached to the reaction time point, the reaction mixture was terminated with with 200 μL methanol. After centrifugation at 14,000 rpm for 10 min, the supernatants were analyzed by UPLC, UV 470 nm.

### Crystallization, data collection and structure determination

Purified DnrK or RdmB proteins were mixed with both the respective **1** (100 mM stock in DMSO) and SAM (1 M stock in buffer) to a molar stoichiometry of 1:5:5 protein:1:SAM. Crystals were obtained by utilizing the sitting drop vapor diffusion method in a 1:1 ratio with the crystallization condition. The co-complex crystals of DnrK with **3** were grown in the precipitating solution of 0.1 M Bis-Tris (pH 5.5), 2.0 M ammonium sulfate and the co-complex crystals of RdmB with **3** were grown in the precipitating solution of 0.1 M Bis-Tris (pH 6.5), 2.0 M ammonium sulfate. Crystals were supplemented with cryoprotectants containing the reservoir contents plus 20% ethylene glycol and flash-frozen in liquid nitrogen. Diffraction data of both DnrK-**3** and RdmB-**3** complex crystals were collected at 100 K on beamline BL18U at the Shanghai Synchrotron Radiation Facility (SSRF) and processed using the HKL3000 program.^[29]^

### Structure determination and refinement

The structure of DnrK-**3** was solved by molecular replacement with the program Phaser,^[30]^ using the DnrK structure (PDB code: 5EEG) structure as the search model, and the structure of RdmB-**3** was solved by molecular replacement, using the RdmB structure (PDB code: 1R00) structure as the search model. Further manual model building was facilitated by using Coot,^[31]^ combined with the structure refinement using Phenix.^[32]^ The statistics of data collection and structure refinement are summarized in table S3. The Ramachandran statistics, as calculated by Molprobity,^[33]^ are 98.36%/0.75%, 97.70%/1.02% (favored/outliers) for structures of DnrK and RdmB, respectively. All the structure diagrams were prepared using the ChimeraX 1.6.1.^[34]^

### Molecular docking of substrate 1 to DnrK and RdmB

To place the substrate into the active site of the DnrK and RdmB, molecular docking of substrate **1** with DnrK and RdmB was performed using Autodock vina 1.2.0,^[35]^ resulting in protein-substrate complex structures. Chem3D 19.0 was used to generate ligand **1**. The final structure of DnrK or RdmB bound with **1** was selected based on the binding state of **3** in the DnrK or RdmB co-crystal structure. After the protein-substrate complex structure was prepared, the substrate-protein interactions were analyzed using PLIP (the protein-ligand interaction profiler) (https://plip-tool.biotec.tu-dresden.de) to determine key catalytic residues. Pymol and ChimeraX 1.6.1 were used for viewing the molecular interactions and image processing.

### ^18^O_2_ labeling experiments

To further verify the catalytic mechanism of RdmB and DnrK, ^18^O labeling water (≥98% labeled, Tenglong Weibo, Qingdao) and ^18^O_2_ (≥98% labeled, Delin, Shanghai) were used to replace water or O_2_ in the reaction system.

For ^18^O_2_ labeling experiments, the reaction was performed in a 100 μL reaction buffer (H_2_O) using liquid bottle (2 mL). 2 μM RdmB or RdmB^L303E^ or 2 μM DnrK or DnrK^E299^ was added to the reaction system, respectively. Then the bottle was purged with nitrogen and sealed with liquid bottle cap. ^18^O_2_ was introduced into the bottle from the compressed gas bag via a syringe needle. Finally, 100 μM **1** was added to the bottle using a microsyringe. After incubation for each enzyme for 60 min at 30 °C, 100 μL methanol was added to stop the reactions.

For H_2_^18^O labeling experiments, the enzymatic reactions of 2 μM RdmB or 2 μM DnrK with 100 μM **1** were performed in a 100 μL reaction buffer (H_2_^18^O) using microcentrifuge tube (1.5 mL). After incubation for RdmB at 30 °C for 10 min or for DnrK at 30 °C for 2 h, 100 μL methanol was added to quench the reactions, respectively. HRESI-LCMS analysis was carried out using the same method as above.

## Acknowledgments

This work was supported by the National Key Research and Development Program of China (2019YFA0905400 and 2019YFA0905700), the National Natural Science Foundation of China (U2106227 and 82022066), the Shandong Provincial Natural Science Foundation (ZR2021ZD28), and the Shenzhen Fundamental Research Program (20220523121619003). We also thank Guannan Lin, Jing Zhu, Zhifeng Li, Jingyao Qu, and Haiyan Sui of the Core Facilities for Life and Environmental Sciences, State Key laboratory of Microbial Technology of Shandong University for HRESI-LCMS and NMR analysis.

## Data Availability Statement

The data that support the findings of this study are available in the supplementary material of this article.

## Reference

[1] a)K. Krohn, Anthracycline chemistry and biology I: biological occurence and biosynthesis, synthesis and chemistry, Vol. 282, Springer, 2009; b)B. Pang, X. Qiao, L. Janssen, A. Velds, T. Groothuis, R. Kerkhoven, M. Nieuwland, H. Ovaa, S. Rottenberg, O. V. Tellingen, J. Janssen, P. Huijgens, W. Zwart, J. Neefjes, Nat. Commun., 2013, 4, 1908.

[2] G. Aubel. Sadron, D. Londos. Gagliardi, Biochimie., 1984, 66, 333–352.

[3] R. P. Warrell Jr, Drugs Exp. Clin. Res., 1986, 12, 275–282.

[4] R. J. Cersosimo, W. K. Hong, J. Clin. Oncol., 1986, 4, 425–439.

[5] E. Barry, J. A. Alvarez, R. E. Scully, T. L. Miller, S. E. Lipshultz, Expert Opin. Pharmacother., 2007, 8, 1039–1058.

[6] M. B. Hulst, T. Grocholski, J. J. C. Neefjes, G. P. van. Wezel, M. Metsä. Ketelä, Nat. Prod. Rep., 2022, 39, 814–841.

[7] C. Hertweck, A. Luzhetskyy, Y. Rebets, A. Bechthold, Nat. Prod. Rep., 2007, 24, 162–190.

[8] a)Y. Wang, J. Niemi, K. Airas, K. Ylihonko, J. Hakala, P. Mäntsälä, Biochim. Biophys. Acta., 2000, 1480, 191–200; b)T. Grocholski, P. Dinis, L. Niiranen, J. Niemi, M. Metsä. Ketelä, Proc. Natl. Acad. Sci. U. S. A., 2015, 112, 9866–9871; c)V. Siitonen, B. Selvaraj, L. Niiranen, Y. Lindqvist, G. Schneider, M. Metsä. Ketelä, Proc. Natl. Acad. Sci. U. S. A., 2016, 113, 5251–5256.

[9] P. Dinis, H. Tirkkonen, B. Nji. Wandi, V. Siitonen, J. Niemi, T. Grocholski, M. Metsä. Ketelä, C. Dupont, Proc. Natl. Acad. Sci. Nexus., 2023, 2. 1–10.

[10] a)K. Madduri, F. Torti, A. L. Colombo, C. Hutchinson, J. Bacteriol., 1993, 175, 3900–3904; b)M. L. Dickens, N. D. Priestley, W. R. Strohl, J. Bacteriol., 1997, 179, 2641–2650.

[11] a)J. Niemi, P. Mäntsälä, J Bacteriol., 1995, 177, 2942–2945; b)A. Jansson, H. Koskiniemi, A. Erola, J. Wang, P. Mäntsälä, G. Schneider, J. Niemi, J. Biol. Chem., 2005. 280, 3636–3644.

[12] a)A. Jansson, J. Niemi, Y. Lindqvist, P. Mäntsälä, G. Schneider, J. Mol. Biol., 2003, 334, 269–280; b)A. Jansson, H. Koskiniemi, P. Mäntsälä, J. Niemi, G. Schneider, J. Biol. Chem., 2004, 279, 41149–41156.

[13] A. Winona. Struck, M. L. Thompson, L. Shin. Wong, J. Micklefield, ChemBioChem, 2012, 13, 2642–2655.

[14] M. Blumauerová, J. Matějů, K. Stajner, Z. Vaněk, Folia. Microbiol., 1977, 22, 275–285.

[15] a)Q. Sun, M. Huang, Y. Wei, Acta Pharm. Sin. B, 2021. 11, 632–650. b)Y. Hsuan. Lee, D. Ren, B. Jeon, H. Wen. Liu, Nat. Prod. Rep., 2023, 40, 1521–1549.

[16] J. K. Coward, E. P. Slisz, J. Med. Chem., 1973, 16, 460–463.

[17] a)T. Awakawa, L. Zhang, T. Wakimoto, S. Hoshino, T. Mori, T. Ito, J. Ishikawa, M. E. Tanner, I. Abe, J. Am. Chem. Soc. 2014, 136, 9910–9913; b)F. Yu, M. Li, C. Xu, B. Sun, H. Zhou, Z. Wang, Q. Xu, M. Xie, G. Zuo, P. Huang, H. Guo, Q. Wang, J. He, Biochem. J., 2016, 473, 4385–4397.

[18] a)M. Ohashi, F. Liu, Y. Hai, M. Chen, M. C. Tang, Z. Yang, M. Sato, K. Watanabe, K. N. Houk, Y. Tang, Nature, 2017, 549, 502–506; b)Z. Chang, T. Ansbacher, L. Zhang, Y. Yang, T. Ping. Ko, G. Zhang, W. Liu, J. Wen. Huang, L. Dai, R. Ting. Guo, D. Thomas. Major, C. Chi Chen, Org. Biomol. Chem., 2019, 17, 2070–2076.

[19] C. Gui, E. Kalkreuter, Y. Chen. Liu, G. Li, A. D. Steele, D. Yang, C. Chang, B. Shen, Nat. Chem. Biol., 2023, 1–8.

[20] N. J. Bauer, A. J. Kreuzman, J. E. Dotzlaf and W. K. Yeh, J. Biol. Chem., 1988, 263, 15619–15625.

[21] D. K. Liscombe, G. V. Louie, J. P. Noel, Nat. Prod. Rep., 2012, 29, 1238–1250.

[22] G. M. Morris, R. Huey, W. Lindstrom, M. F. Sanner, R. K. Belew, D. S. Goodsell, A. J. Olson, J. Comput. Chem., 2009, 30, 2785–2791.

[23] a)V. Siitonen, B. Blauenburg, P. Kallio, P. Mäntsälä, M. Metsä. Ketelä, J. Biol. Chem., 2012, 19, 638–646; b)M. M. Machovina, R. J. Usselman, J. L. DuBois, J. Biol. Chem., 2016, 291, 17816–17828.

[24] C. Huang, C. V. Smith, M. S. Glickman, W. R. Jacobs Jr, J. C. Sacchettini, J. Biol. Chem., 2002, 277, 11559–11569.

[25] X. Robert, P. Gouet, Nucleic Acids Res., 2014, 42, 320–324.

[26] C. Notredame, D. G. Higgins, J. Heringa, J. Mol. biol., 2000, 302, 205–217.

[27] K. Blin, S. Shaw, A. M. Kloosterman, Z. Charlop. Powers, G. P. van Wezel, M. H. Medema, T. Weber, Nucleic Acids Res., 2021, 49, W29-W35.

[28] T. Kieser, M. J. Bibb, M. J. Buttner, K. F. Chater, D. A. Hopwood, Practical streptomyces genetics, Vol. 291, John Innes Foundation Norwich, 2000.

[29] W. Minor, M. Cymborowski, Z. Otwinowski, M. Chruszcz, Acta Crystallogr., Sect. D: Biol. Crystallogr., 2006, 62, 859–866.

[30] A. J. McCoy, R. W. Grosse. Kunstleve, P. D. Adams, M. D Winn, L. C Storoni, R. J Read, J. Appl. Crystallogr., 2007, 40, 658–674.

[31] P. Emsley, K. Cowtan, Acta Crystallogr., Sect. D: Biol. Crystallogr., 2004, 60, 2126–2132.

[32] P. D. Adams, P. V. Afonine, G. Bunkóczi, V. B. Chen, I. W. Davis, N. Echols, J. J. Headd, L. Wei. Hung, G. J. Kapral, R. W. Grosse. Kunstleve, A. J. McCoy, N. W. Moriarty, R. Oeffner, R. J. Read, D. C. Richardson, J. S. Richardson, T. C. Terwilliger, P. H. Zwart, Acta Crystallogr., Sect. D: Biol. Crystallogr., 2010, 66, 213–221.

[33] I. W. Davis, A. Leaver. Fay, V. B Chen, J. N. Block, G. J. Kapral, X. Wang, L. W. Murray, W. B. Arendall, J. Snoeyink, J. S. Richardson, D. C. Richardson, Nucleic Acids Res., 2007, 35, 375–383.

[34] E. F. Pettersen, T. D. Goddard, C. C. Huang, E. C. Meng, G. S. Couch, T. I. Croll, J. H. Morris, T. E. Ferrin, Protein Sci., 2021, 30, 70–82.

[35] G. M. Morris, R. Huey, W. Lindstrom, M. F. Sanner, R. K. Belew, D. S. Goodsell, A. J. Olson, J. Comput. Chem., 2009, 30, 2785–2791.

