## Supplementary Information for "Glutamate (E299) is a key residue in the evolutionarily divergence of the SAM-dependent methyltransferases DnrK and RdmB in anthracycline biosynthesis"

[b] Prof. Dr. W. Zhang
Laboratory for Marine Biology and Biotechnology，Qingdao Marine Science and Technology Center, Qingdao, Shandong, 266237, China

### Supplementary Tables

#### Table S1 Primers used for *dnrK* gene knockout.

| Primers | Sequence (5’-3’) |
|---|---|
| dnrK-sgRNA-F | agtcctaggtataatactagtCCCACGTACGAGTCCATCTAgttttagagctagaaata |
| dnrK-sgRNA-R | ctcAAAAAAagcaccgactcggt |
| dnrK-LF | cgagtcggtgctttttttgagctgggtgtgggaccagtccac |
| dnrK-LR | ggtgtgtccgtgctcctgtcg |
| dnrK-RF | acaggagcacggacacaccggagcacgacgatgcccacac |
| dnrK-RR | cgacggccagtgccaagctt ctccctgggtgatcaggctg |
| dnrK-VF | tgccacagcagacgcatgcc |
| dnrK-VR | cgaagtcgcaccgggaggagc |

The guide sequence sgRNA were shown in capital letter.

#### Table S2 Primers used in site-directed mutagenesis for *dnrK* gene.

| Primers | Sequence (5’-3’) |
|---|---|
| dnrK$^{WT}$-F | tgccgcgcggcagccatatgacagccgaaccgacg |
| dnrK$^{WT}$-R | tggtggtggtgctcgagtcaggcgccggtggccgc |
| dnrK$^{3GA}$-F | gtgctggatgtg**gca**ggc**gca**aaa**gca**ggttttgcggcggcgatt |
| dnrK$^{3GA}$-R | cgcAAAacc**tgc**ttt**tgc**gcc**tgc**cacatccagcacatgacgaac |
| DnrK$^{N257A}$-F | tttgtgctgctg**gca**tggccggatcatgatgcggtg |
| DnrK$^{N257A}$-R | atgatccggcca**tgc**cagcagcacaaagctcagaat |
| DnrK$^{R303A}$-F | ctggacctg**gca**atgctggtgtttctgggcggc |
| dnrK$^{R303A}$ -R | caccagcat**tgc**caggtccagctcggtgaactg |
| dnrK$^{Y143A}$-F | ggcaaaccgttt**gca**gaggacctggcgggccgc |
| dnrK$^{Y143A}$-R | caggtcctc**tgc**aaacggtttgccataaatgct |
| dnrK$^{E299A}$-F | cagttcacc**gca**ctggacctgcgcatgctg |
| dnrK$^{E299A}$-R | caggtccag**tgc**ggtgaactgttcattaaagct |
| dnrK$^{E299L}$-F | cagttcacc**ctt**ctggacctgcgcatgctg |
| dnrK$^{E299L}$-R | caggtccag**aag**ggtgaactgttcattaaagct |
| dnrK$^{E299D}$-F | cagttcacc**gat**ctggacctgcgcatgctg |
| dnrK$^{E299D}$-R | caggtccag**atc**ggtgaactgttcattaaagct |
| dnrK$^{E299Q}$-F | cagttcacc**caa**ctggacctgcgcatgctg |
| dnrK$^{E299Q}$-R | caggtccag**ttg**ggtgaactgttcattaaagct |
| dnrK$^{E299K}$-F | cagttcacc**aag**ctggacctgcgcatgctg |
| dnrK$^{E299K}$-R | caggtccag**ctt**ggtgaactgttcattaaagct |
| dnrK$^{E299A/R303A}$-F | cagttcacc**gca**ctggacctg**gca**atgctggtgtttctg |
| dnrK$^{E299A/R303A}$-R | ccagcat**tgc**caggtccag**tgc**ggtgaactgttcattaaa |

The site-specific mutation sites of recombinant proteins in bold.

### Table S3 Primers used in site-directed mutagenesis for *rdmB* gene.

| Primers | Sequence (5'-3') |
|---|---|
| RdmB$^{3GA}$-F | ttggatgtg**gca**gcg**gca**aat**gca**ggcatgtta |
| RdmB$^{3GA}$-R | taacatgcc**tgc**att**tgc**cgc**tgc**cacatccaacacatg |
| RdmB$^{N260A}$-F | gtgctgctg**gca**tggagcgatgaagatgcgctg |
| RdmB$^{N260A}$-R | atcgctcca**tgc**cagcagcacAAAgcttaataa |
| RdmB$^{R307A}$-F | ctggatctg**gca**atgctgacctttatgggcggccgc |
| RdmB$^{R307A}$-R | AAAggtcagcat**tgc**cagatccagcagggtgcta |
| RdmB$^{L303E}$-F | tttagcacc**gaa**ctggatctgcgcatgctg |
| RdmB$^{L303E}$-R | cagatccag**ttc**ggtgctaaaaaaacgatccgc |

The site-specific mutation sites of recombinant proteins in bold.

### Table S4 Data collection and refinement statistics

| | DnrK + **1** | RdmB + **1** |
|---|---|---|
| **PDB code** | 8KHI | 8KHJ |
| **Data collection** | | |
| Space Group | P 1 21 1 | P 32 2 1 |
| Cell dimensions | | |
| *a, b, c* (Å) | 60.40,101.71,62.43 | 79.997,79.997,234.203 |
| a, b, g (°) | 90.00,102.55,90.00 | 90.00,90.00,120.00 |
| Resolution (Å) | 58.96-1.29 (1.36-1.29) * | 69.28-2.10 (2.21-2.10) |
| *R*sym or *R*merge | 11.1(27.1) | 9.5 (121.7) |
| CC1/2 | 0.993 (0.873) | 0.999 (0.910) |
| $I/\delta I$ | 11.1 (2.8) | 19.2 (3.0) |
| Completeness (%) | 81.5 (27.9) | 95.6 (99.9) |
| Redundancy | 5.8 (2.7) | 17.5 (18.3) |
| **Refinement** | | |
| Resolution (Å) | 38.508-1.560 (1.616-1.560) | 44.719 - 2.098 (2.173 - 2.098) |
| Completeness (%) | 99.96 (100.00) | 95.54 (99.61) |
| No. reflections | 104403 (10376) | 49468 (5052) |
| $R_{work}/R_{free}$ | 15.9/18.6 | 21.3/25.6 |
| No. atoms | | |
| Protein | 5245 | 4976 |
| Ligand/ion | 74 | 74 |
| Water | 1018 | 229 |
| B-factors | | |
| Protein | 14.9 | 46.57 |
| Ligand/Ion | 13.87 | 56.05 |
| Water | 26.97 | 46.76 |
| R.m.s. deviations | | |
| Bond lengths (Å) | 0.0061 | 0.0065 |
| Bond angles (°) | 0.97 | 0.92 |

One crystal was used for each structure.

*Values in parentheses are for highest-resolution shell.

### Supplementary Figures

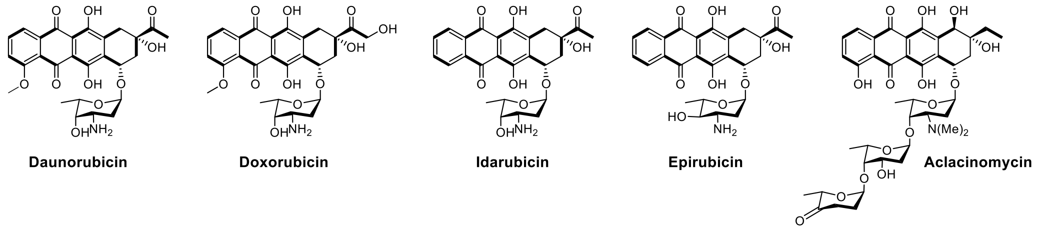

**Figure S1**. Chemical structures of anthracyclines.

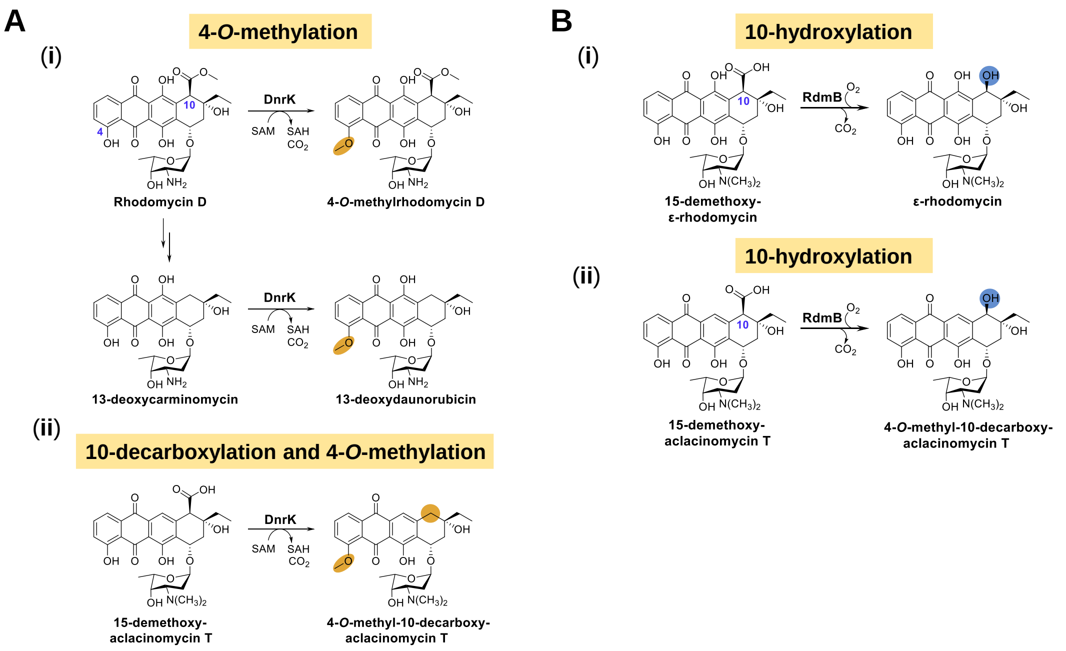

**Figure S2**. Chemical reactions catalyzed by methyltransferases DnrK (A) and RdmB (B).

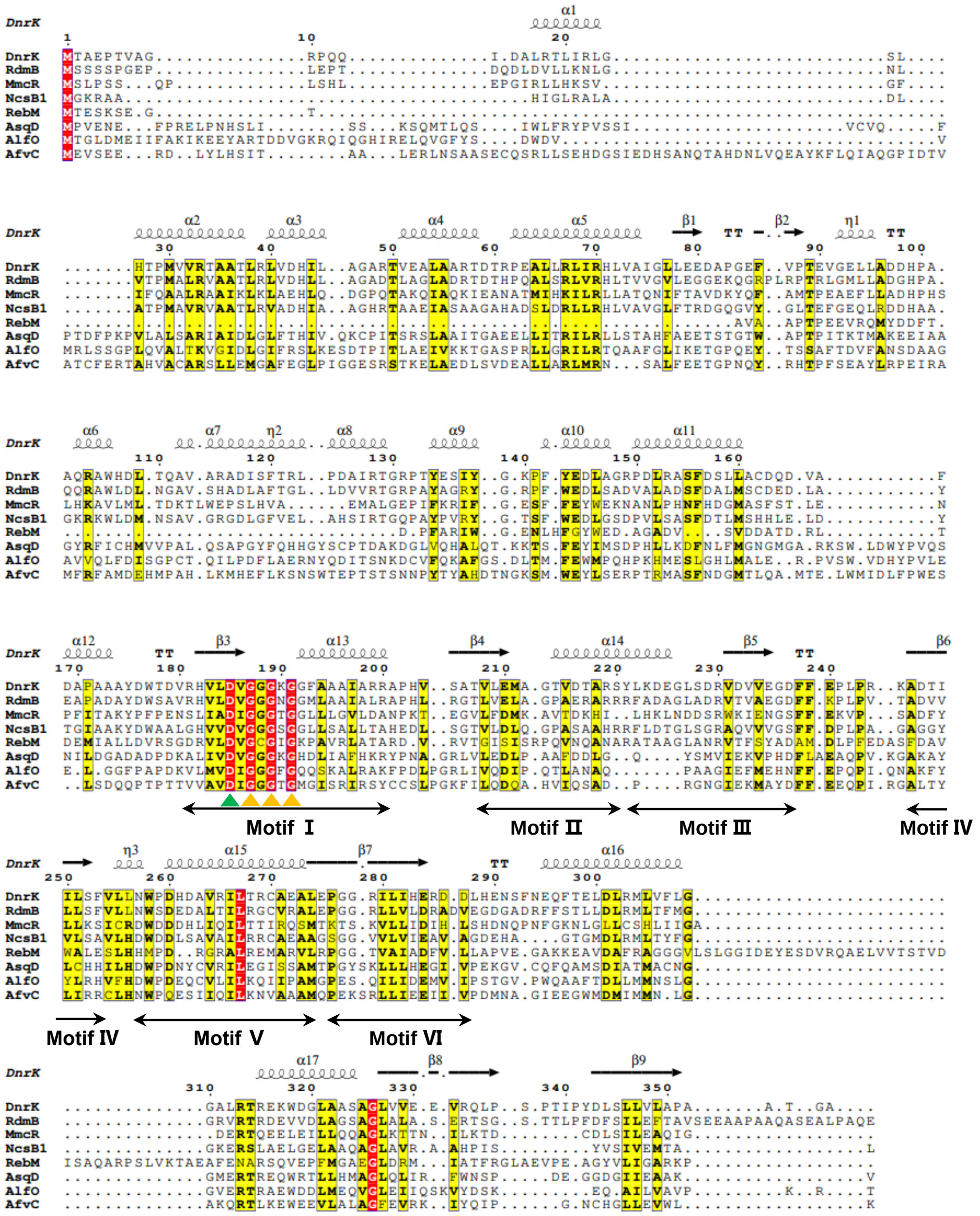

**Figure S3.** Multiple sequence alignments of DnrK and RdmB and typical *O*-methyltransferases including MmcR (accession No. CAG9426959.1); NcsB1 (accession No. Q84HC8.1); RebM (accession No. Q8KZ94.1); AsqD (accession No. Q5AR47.1); AlfO (accession No. Q12120.1); AfvC (accession No. B8N8R1.1). The sequence analysis shows the high sequence similarity of DnrK and RdmB to typical *O*-methyltransferases. Color triangles indicate the conserved residues in motifs I for SAM binding.

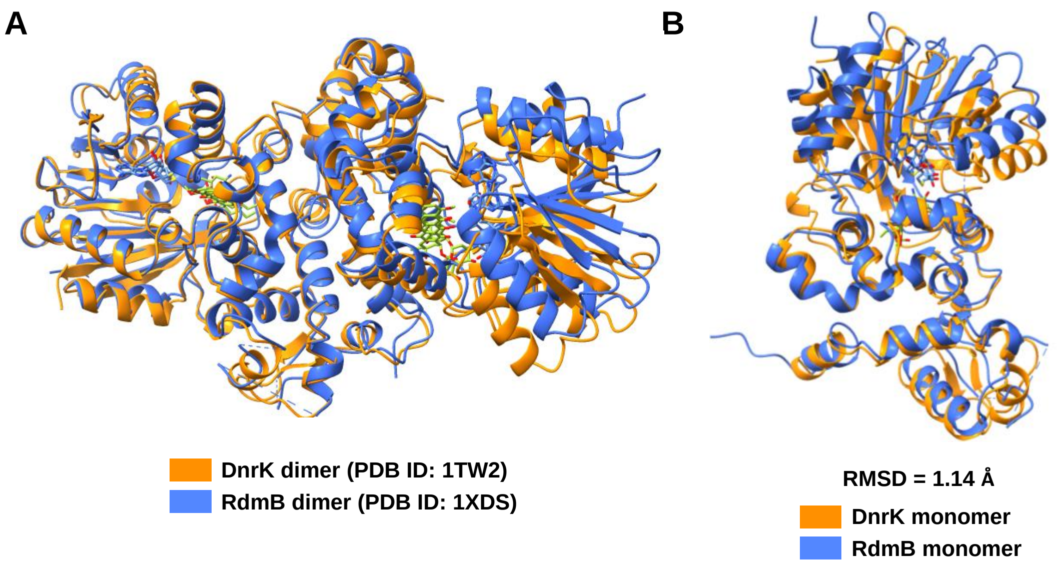

**Figure S4.** 3D structural alignment of DnrK (PDB: 1TW2) and RdmB (PDB: 1XDS). (A) Ribbon diagram shows the overall dimeric structural comparison of DnrK-SAH-4-methoxy-*ε*-rhodomycin T and RdmB-SAM-11-deoxy-*β*-rhodomycin. (B) Ribbon representation shows the structural comparison of the monomeric DnrK with SAH and the monomeric RdmB with SAM, respectively. DnrK is shown in orange color, while RdmB is shown in violet color.

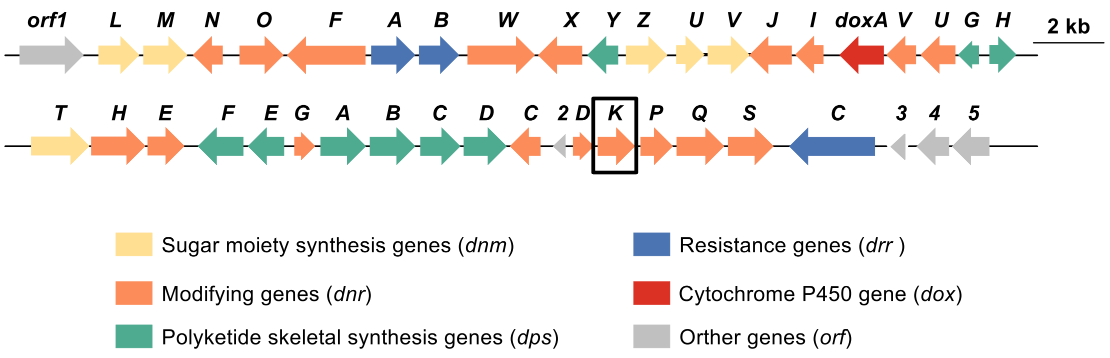

**Figure S5.** The biosynthetic gene cluster of doxorubicin from *Streptomyces coerleorubidus*. The black box indicates the SAM-dependent methyltransferase coding gene *dnrK*.

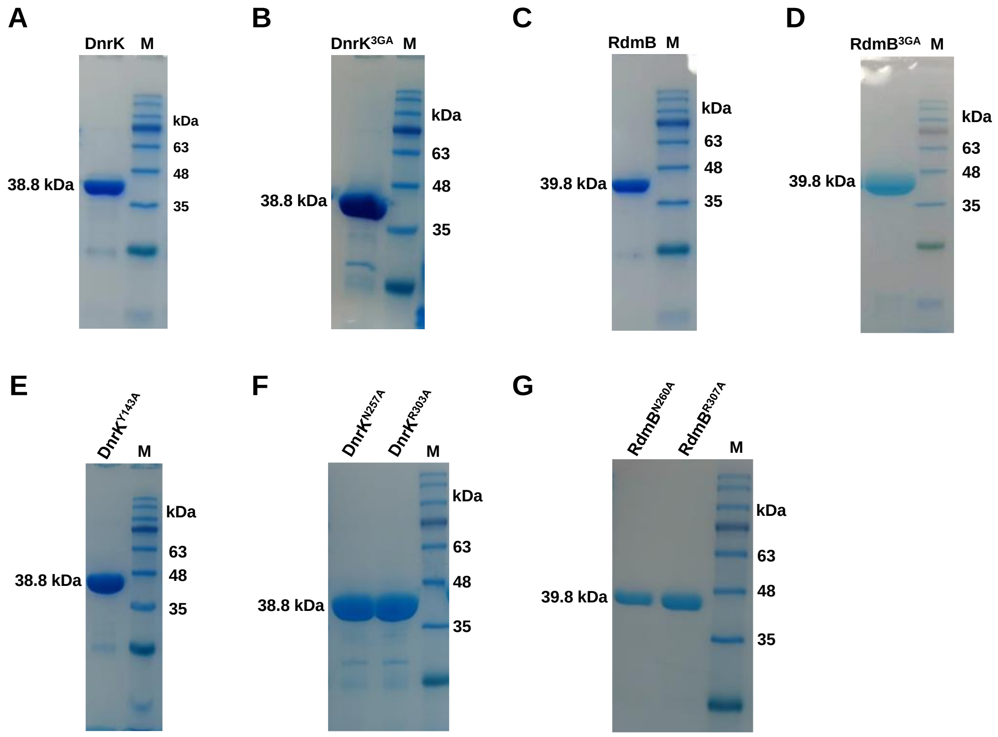

**Figure S6.** SDS-PAGEs analysis for the purified proteins. (A) DnrK; (B) DnrK$^{3GA}$; (C) RdmB; (D) RdmB$^{3GA}$; (E) DnrK$^{Y143A}$; (F) DnrK$^{N257A}$ and DnrK$^{R303A}$; (G) RdmB$^{N260A}$ and RdmB$^{R307A}$; M (Marker 180).

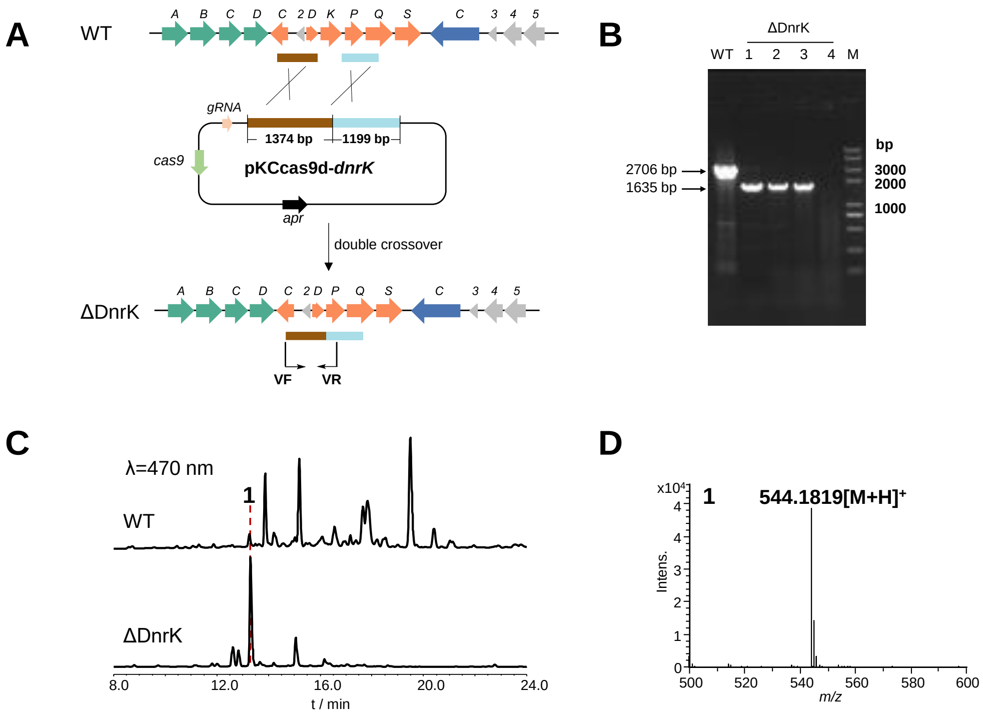

**Figure S7.** The construction and fermentation products detection of mutant ΔDnrK. (A) Gene *dnrK* in-frame deletion in *S. coeruleorubidus*. (B) PCR verification of *S. coeruleorubidus::*ΔDnrK mutant. The arrows indicate the expected size of the fragments from the wild-type (WT) and mutant DNA, respectively. M, Trans2K® Plus II DNA marker. (C) HPLC profiles of metabolites produced by *S. coeruleorubidus* and *S. coeruleorubidus::*ΔDnrK mutant. (D) HRESI-LCMS spectrum of **1** in positive ion mode**.**

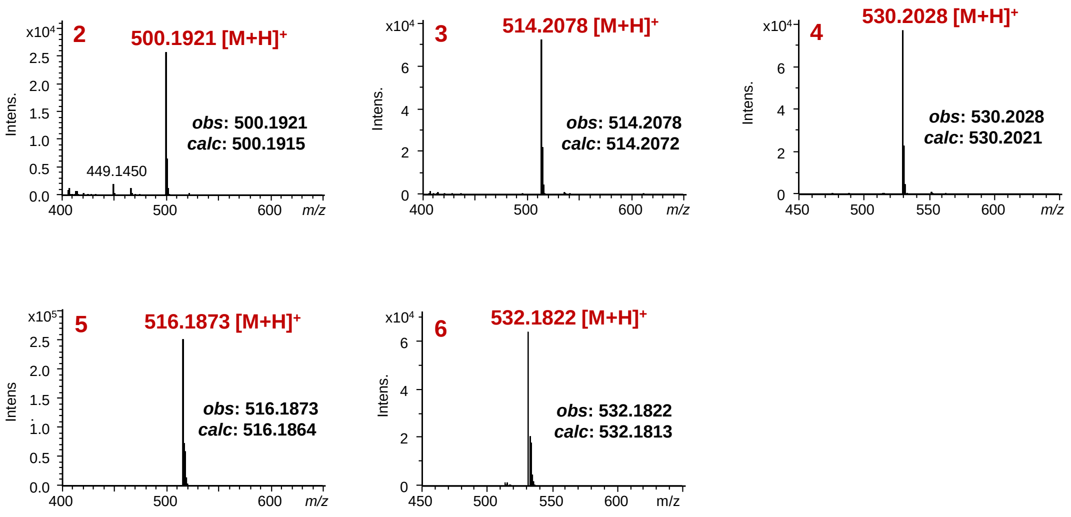

**Figure S8.** HRESI-LCMS analysis of **2-6** in positive ion mode.

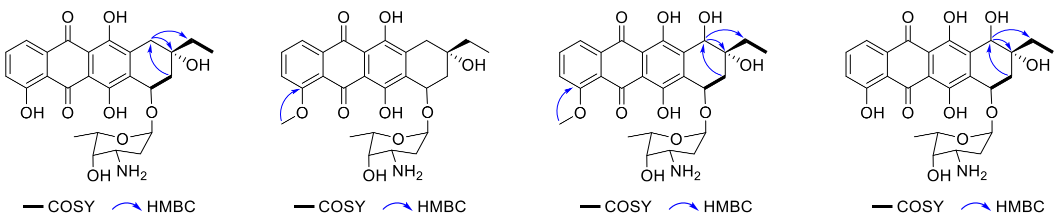

**2** **3** **4** **5**

**Figure S9.** Key 2D NMR correlations of compounds **2-5**.

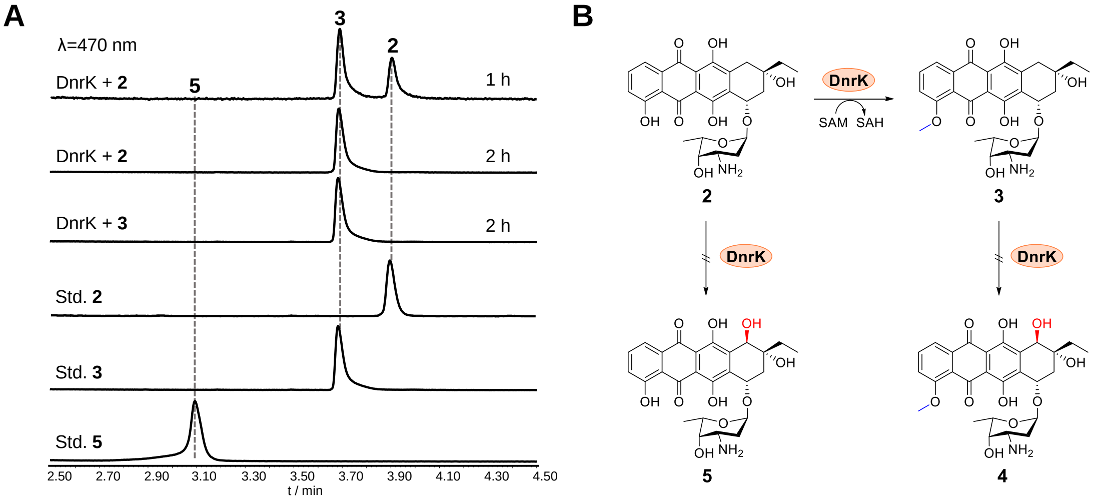

**Figure S10**. UPLC analysis of the conversion of **2** and **3** by DnrK (A) and the reaction scheme (B).

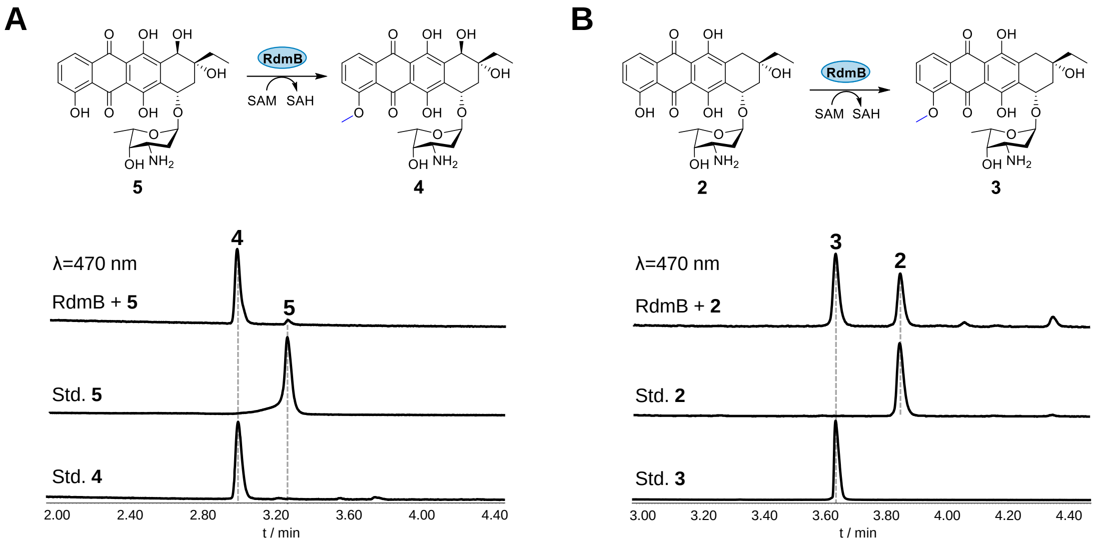

**Figure S11.** UPLC analysis of the *in vitro* enzymatic conversion of **5** (A) and **2** (B) catalyzed by RdmB, respectively.

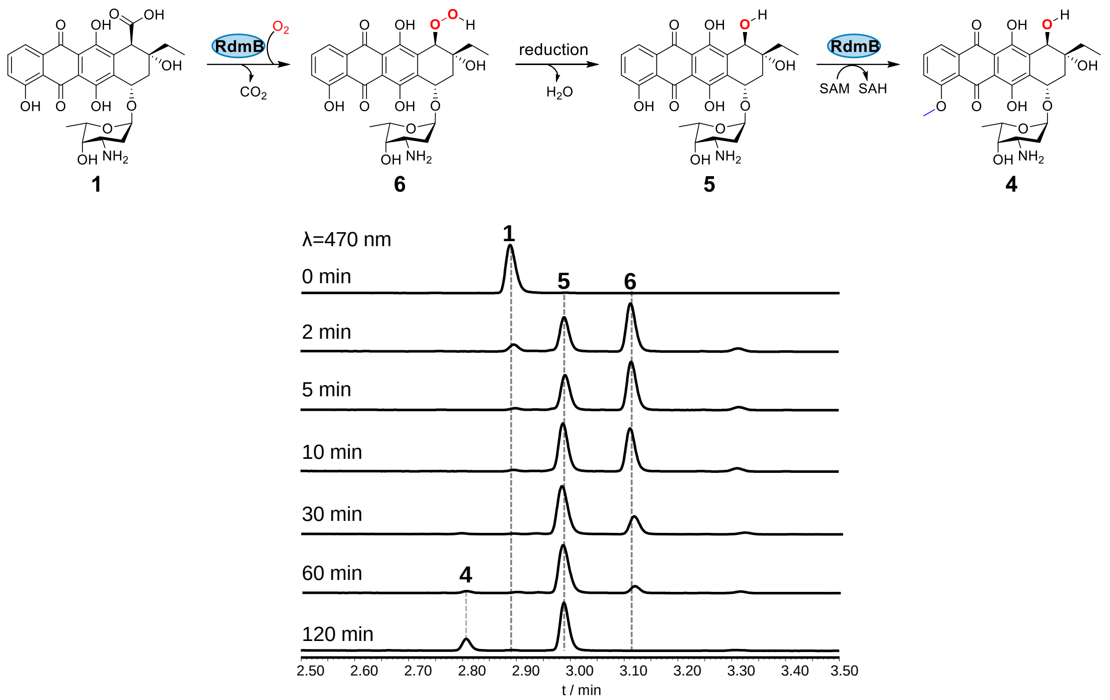

**Figure S12**. Time-course analysis of the stepwise conversion of **1**→**6**→**5**→**4** by RdmB in the presence of SAM.

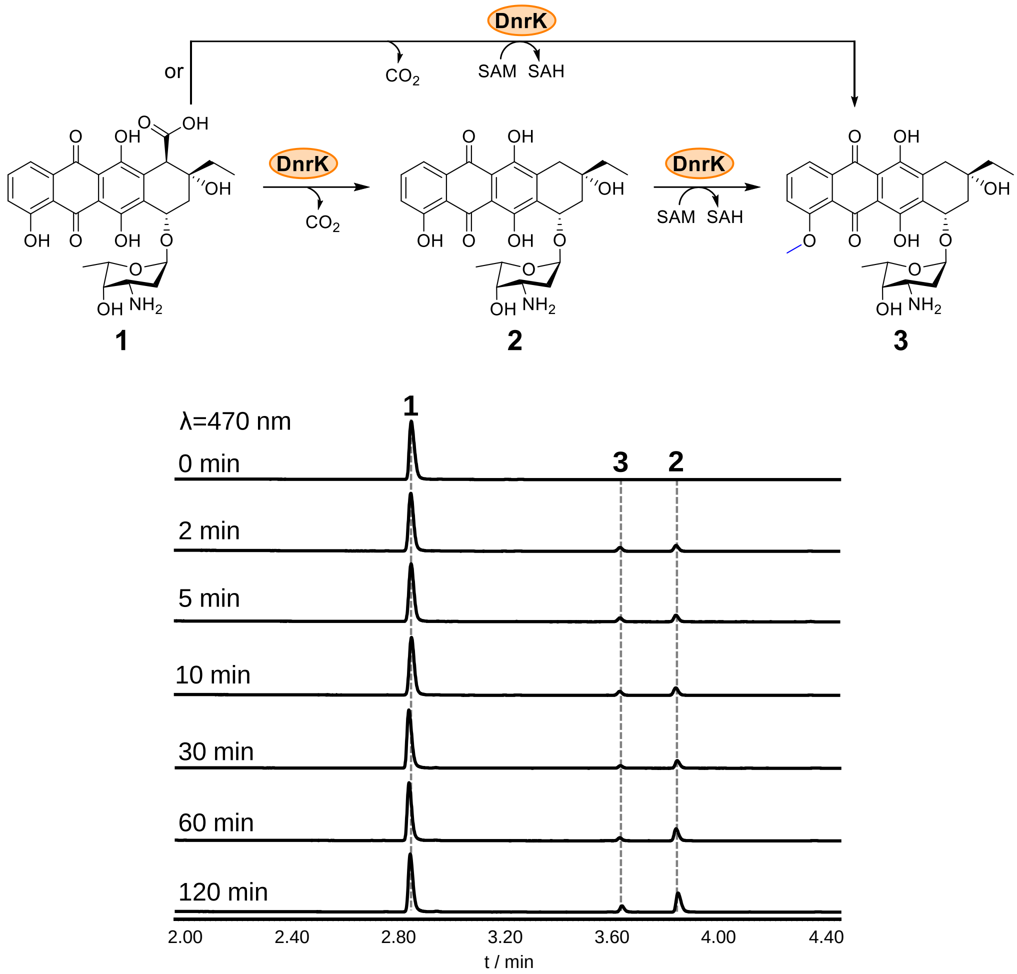

**Figure S13**. Time-course analysis of the concerted conversion of **1** by DnrK without SAM.

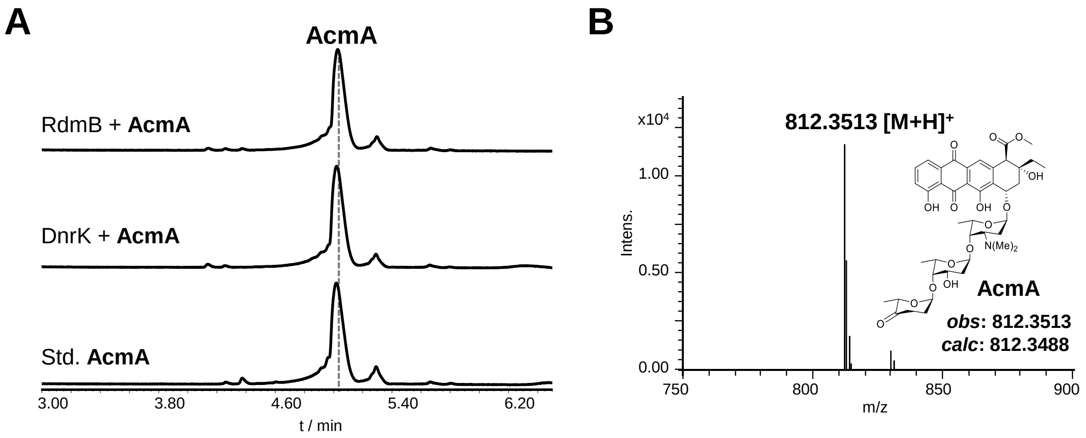

**Figure S14.** (A) UPLC analysis of the conversion of triglycosylated anthracycline aclacinomycin A (AcmA) catalyzed by DnrK and RdmB. (B) HRESI-LCMS spectrum of aclacinomycin A in positive ion mode.

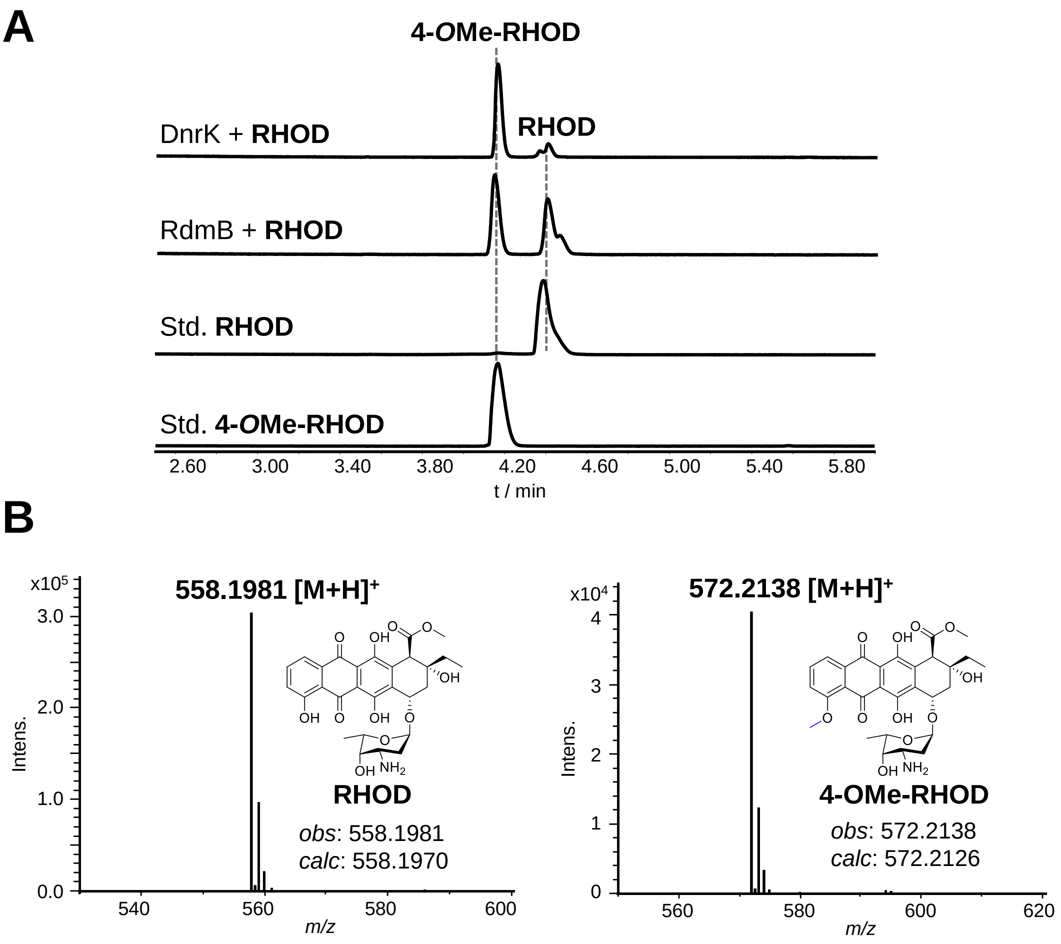

**Figure S15.** (A) UPLC analysis of the *in vitro* enzymatic conversion of rhodomycin D (RHOD) to 4-*O*Me-RHOD by DnrK or RdmB. (B) HRESI-LCMS analysis of RHOD and 4-*O*Me-RHOD. Control: negative control with boiled RdmB.

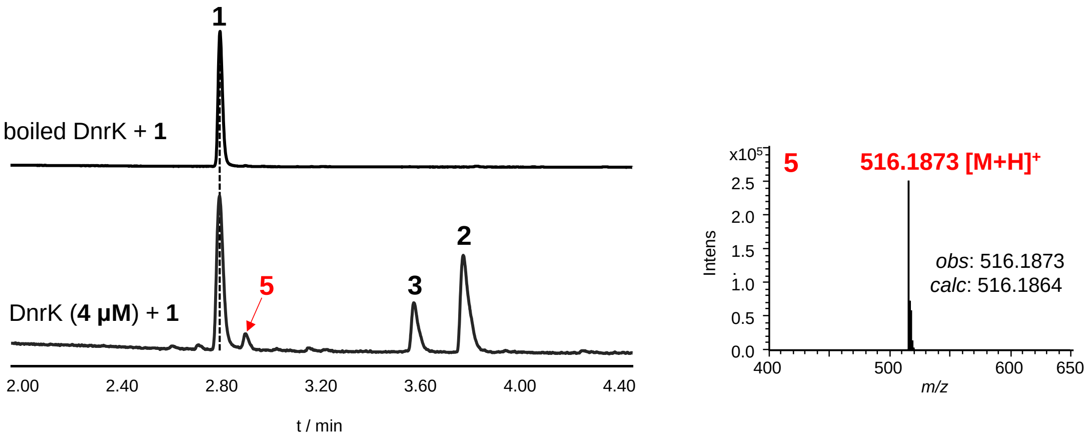

**Figure S16**. UPLC analysis of the formation of **5** catalyzed by DnrK (A) and HRESI-LCMS data of **5** in positive ion mode (B).

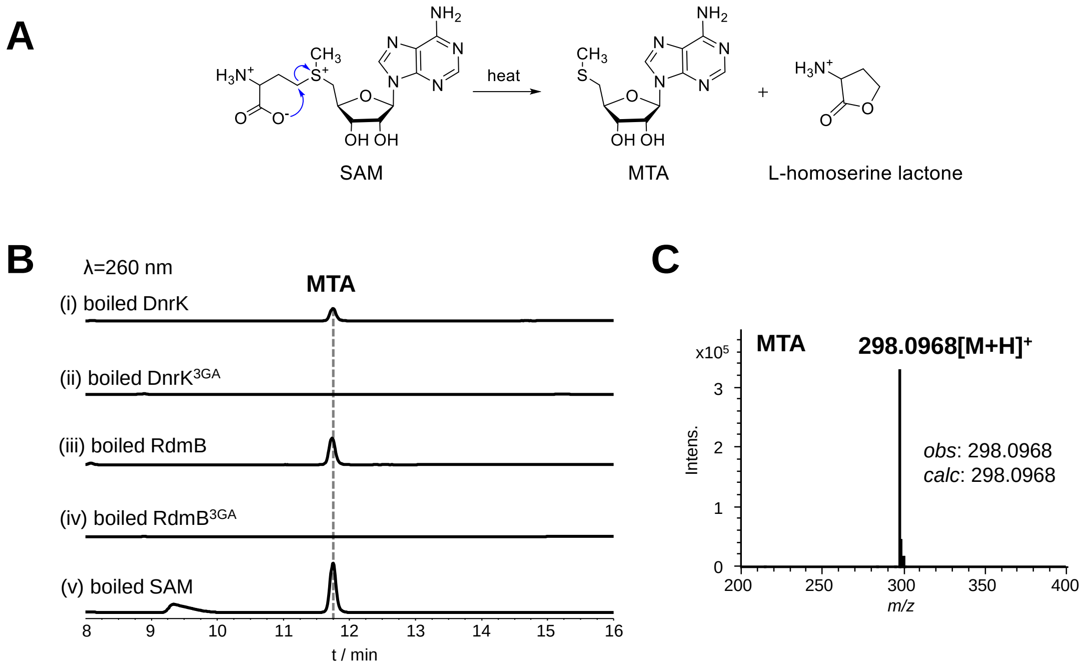

**Figure S17**. HPLC analysis of co-purified SAM after DnrK, RdmB, DnrK$^{3GA}$. (A) Reaction of SAM degradation to produce 5′-deoxy-5′-(methylthio) adenosine (MTA). (B) When DnrK and RdmB were denatured by heating 100°C for 10 min, MTA as the major degradation product of SAM was detected. (i) DnrK heated at 100°C for 10 min; (ii) DnrK$^{3GA}$ heated at 100°C for 10 min; (iii) RdmB heated at 100°C for 10 min; (iv) RdmB$^{3GA}$ heated at 100°C for 10 min; (v) Standard SAM heated at 100°C for 10 min. (C) HRESI-LCMS spectrum of MTA in positive ion mode.

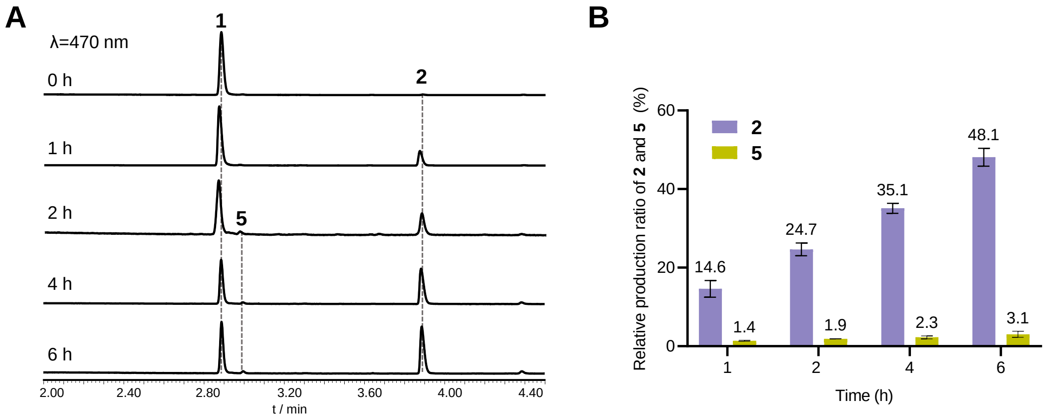

**Figure S18**. Time-course analysis conversion of **1** catalyzed by mutant DnrK$^{3GA}$ in the absence of SAM.

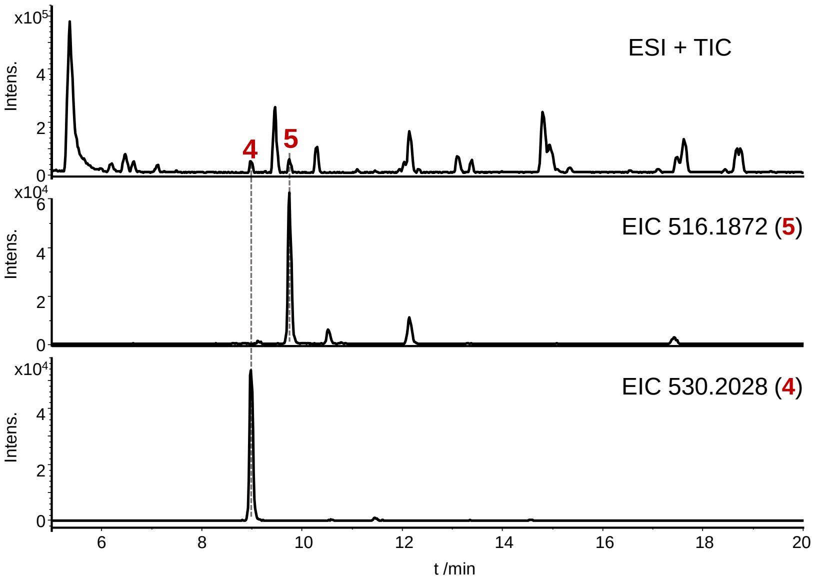

**Figure S19.** The ion chromatograms of compounds **4** and **5** in the extracts of *S. coeruleorubidus* by HRESI-LCMS analysis.

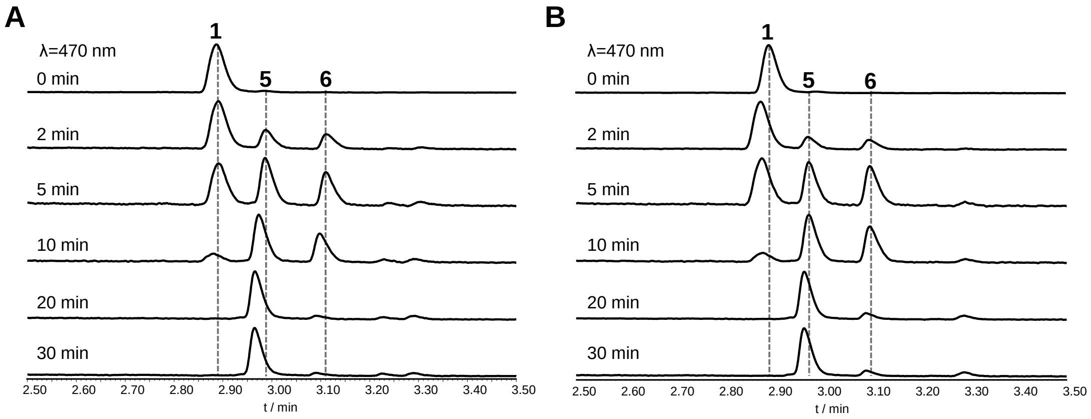

**Figure S20.** Time-course analysis of the *in vitro* reaction of **1** (50 μM) with 1 μM RdmB in the presence of 200 μM SAH (A) or 200 μM sinefungin (B).

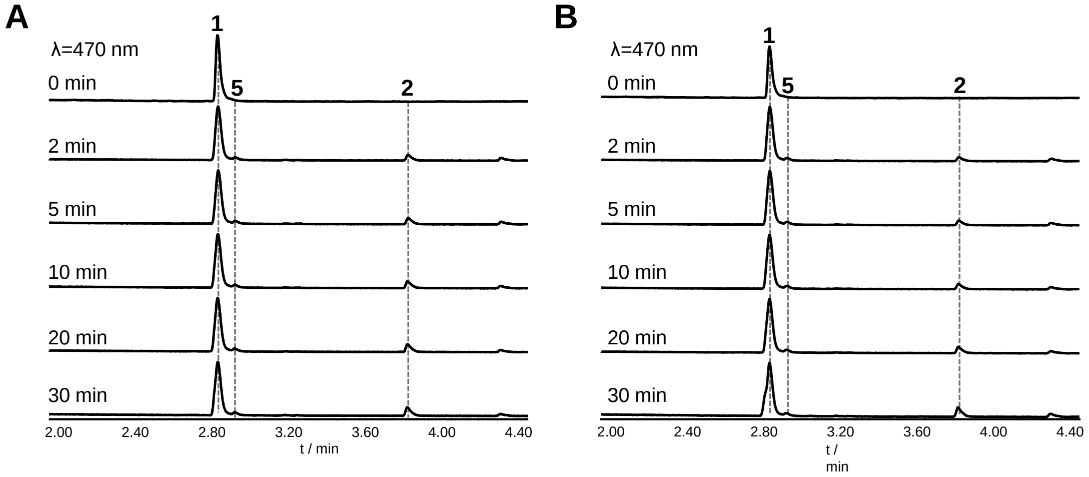

**Figure S21.** Time-course analysis of the *in vitro* reaction of **1** (50 μM) with 1 μM DnrK in the presence of 200 μM SAH (A) or 200 μM sinefungin (B).

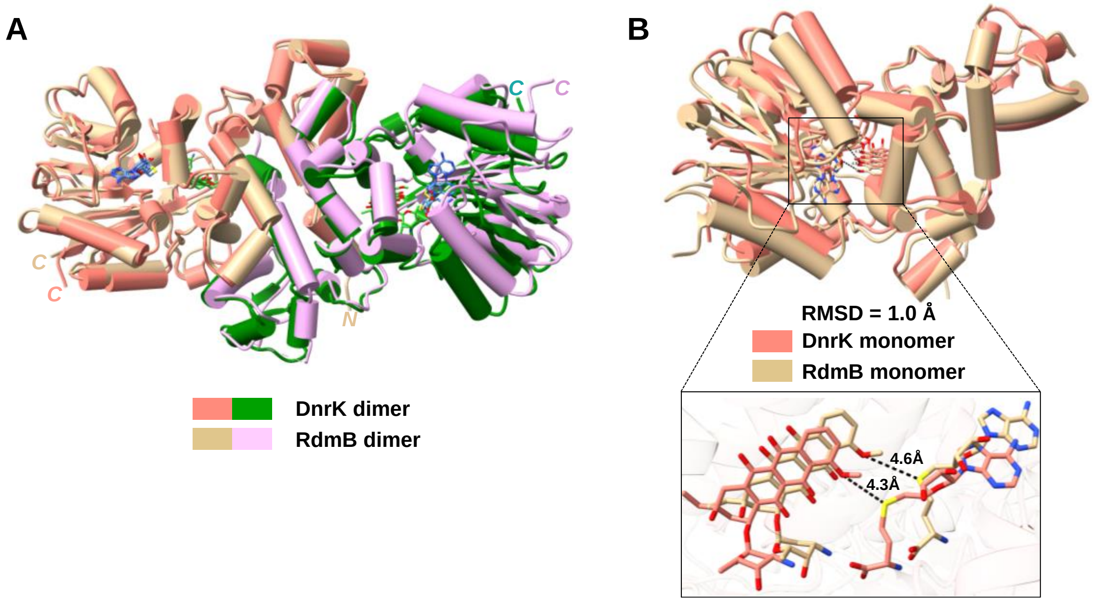

**Figure S22**. Structural alignment of the overall structure of DnrK (PDB: 8KHI) and RdmB (PDB: 8KHJ) obtained in this study. (A) Ribbon diagram showing the overall structural comparison of the dimeric DnrK-SAH-**3** and RdmB-SAM-**3**. The two monomeric of DnrK is shown in pink and green color, while RdmB is shown in tan and purple color, respectively. (B) Ribbon representation showing the structural comparison of the monomeric DnrK-SAH-**3** and the monomeric RdmB-SAH-**3**, respectively. The box shows a zoom-in view of the binding of SAH/**3** in the active pockets of DnrK and RdmB.

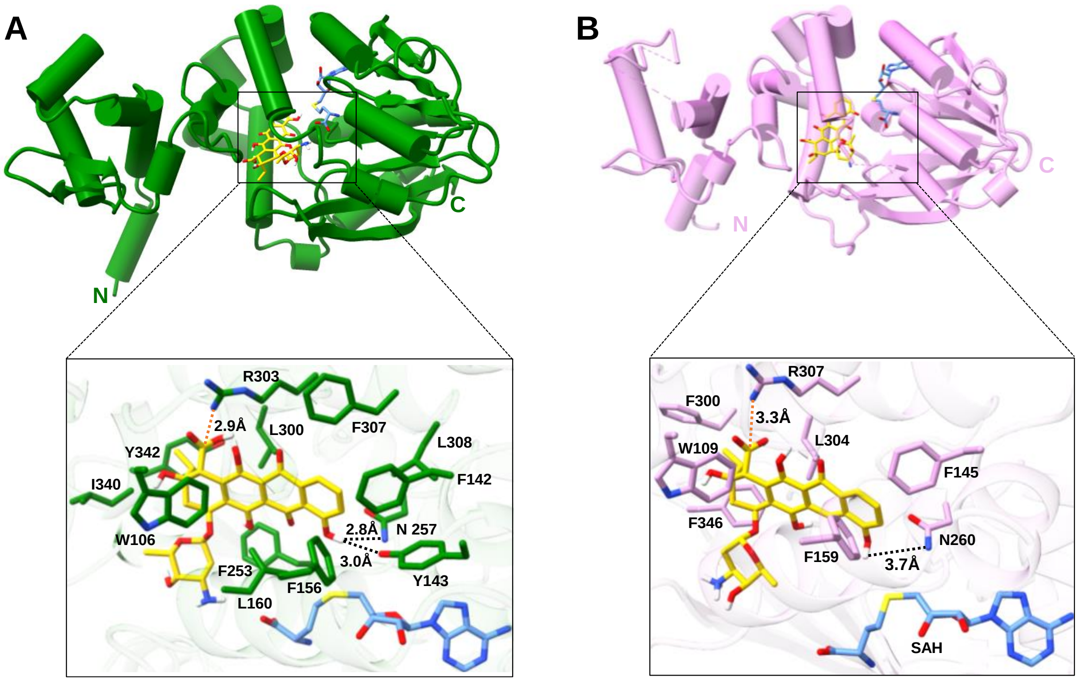

**Figure S23**. The docking model of DnrK and RdmB in complex with the substrate **1**. (A) Molecular docking of **1** with DnrK. The box shows a zoom-in view of the active site of DnrK with docked **1**. In this view, the combined ribbon diagram and the stick-ball model show the detailed interactions between DnrK with docked **1**. The hydrogen bonds, salt bridges, and hydrophobic interactions are shown as black, orange, and purple dotted lines, respectively. (B) Molecular docking of **1** with RdmB. The box shows a zoom-in view of the active site of RdmB with docked **1**. In this view, the combined ribbon diagram and the stick-ball model show the detailed interactions between RdmB with docked **1**. The hydrogen bonds and salt bridges are shown as black and orange dotted lines, respectively.

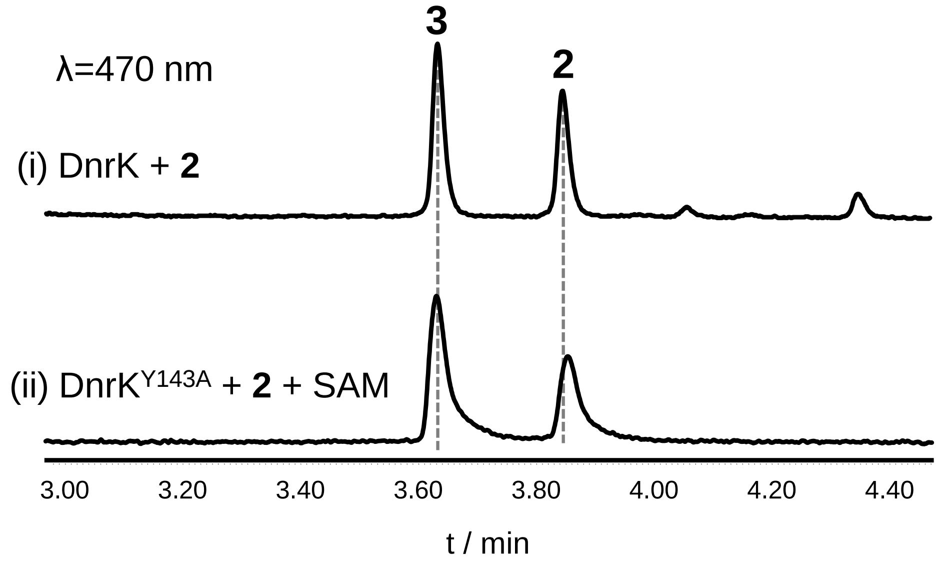

**Figure S24.** UPLC profiles of the conversion of **2** catalyzed by wild-type DnrK (i) and its mutant DnrK$^{Y143A}$ (ii).

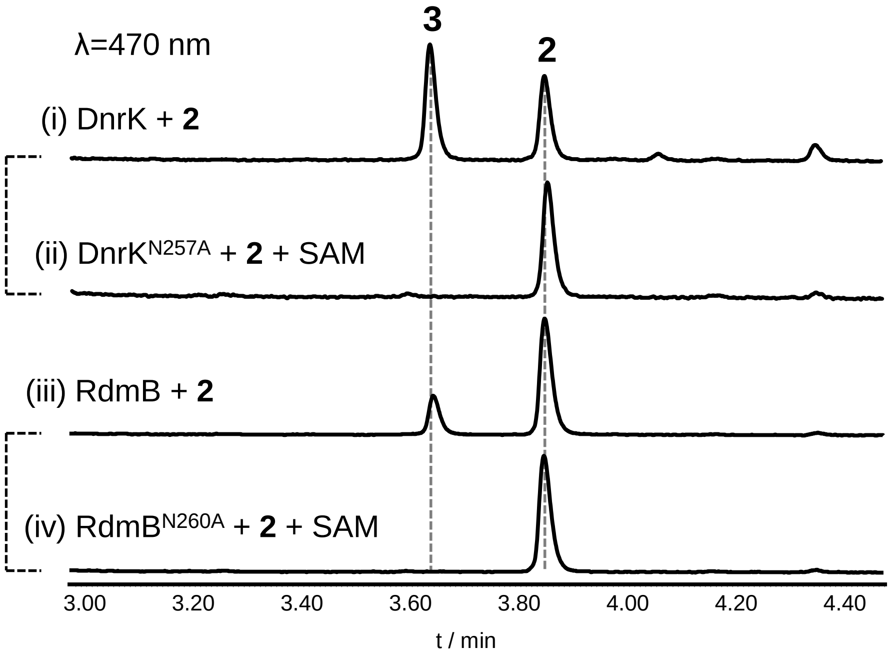

**Figure S25.** UPLC analysis of the conversion of **2** by DnrK (i), DnrK$^{N257A}$ (ii), RdmB (iii), and RdmB$^{N260A}$ (iv).

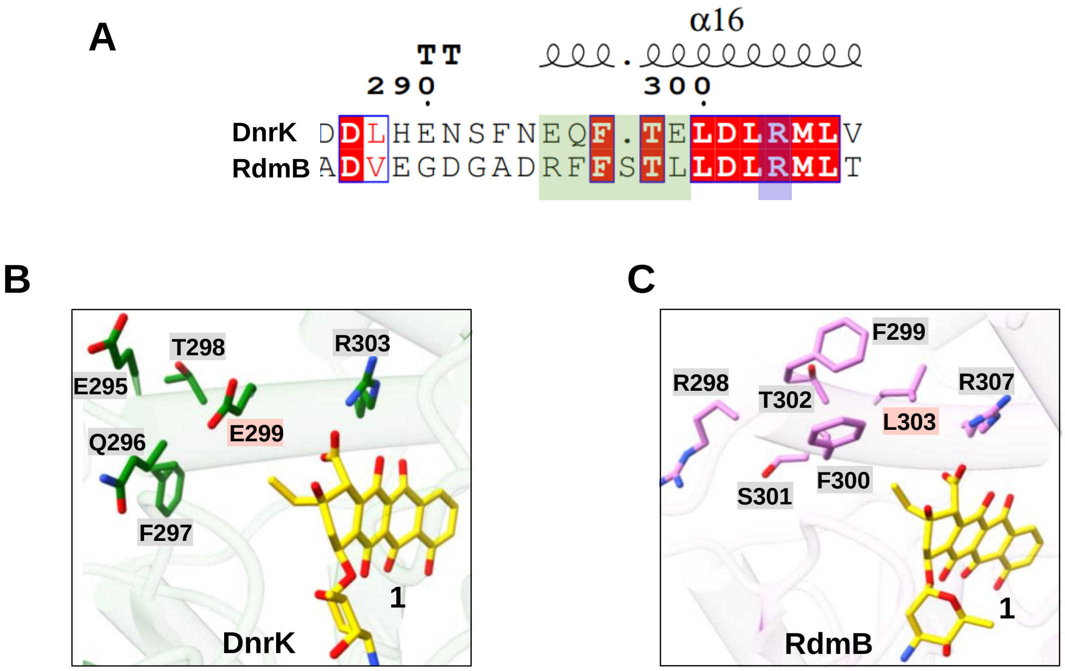

**Figure S26.** Sequence alignment and molecular docking analysis of DnrK and RdmB with **1**. (A) Sequence alignment of residues near DnrK R303 and RdmB R307. (B) The display of key residues near the C-10 position of **1** in DnrK. (C) The display of key residues near the C-10 position of **1** in RdmB.

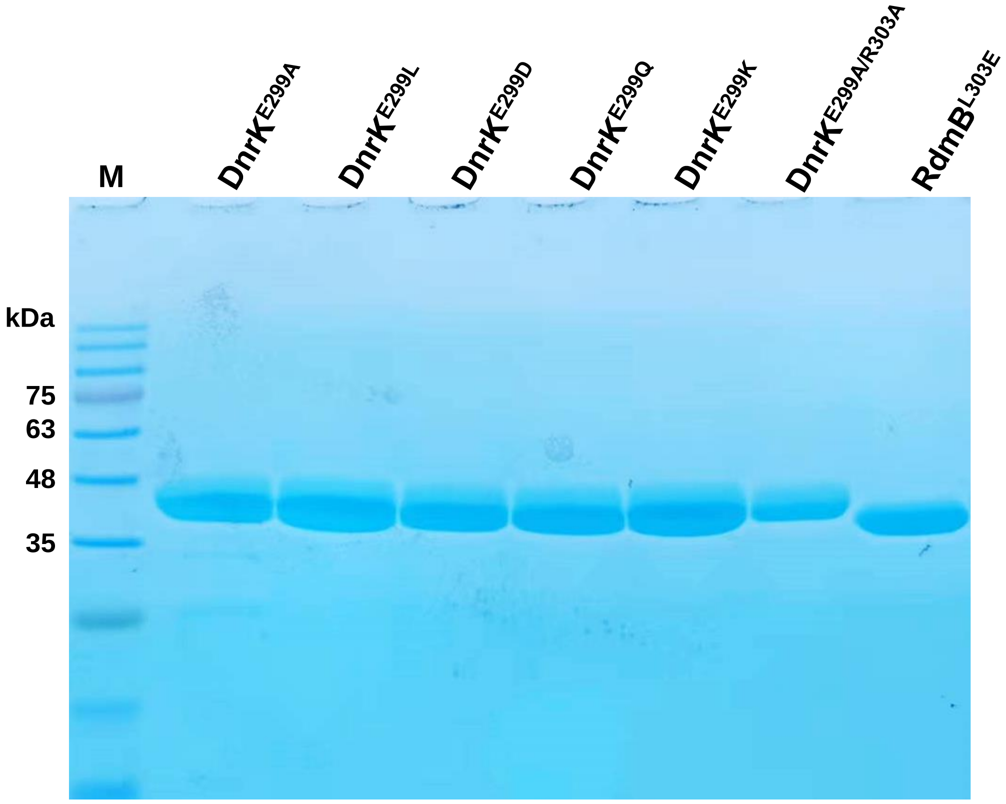

**Figure S27.** SDS-PAGEs analysis of the purified proteins DnrK$^{E299A}$, DnrK$^{E299L}$, DnrK$^{E299D}$, DnrK$^{E299Q}$, DnrK$^{E299K}$, DnrK$^{E299A/R303A}$, RdmB$^{L303E}$; M (Marker 180).

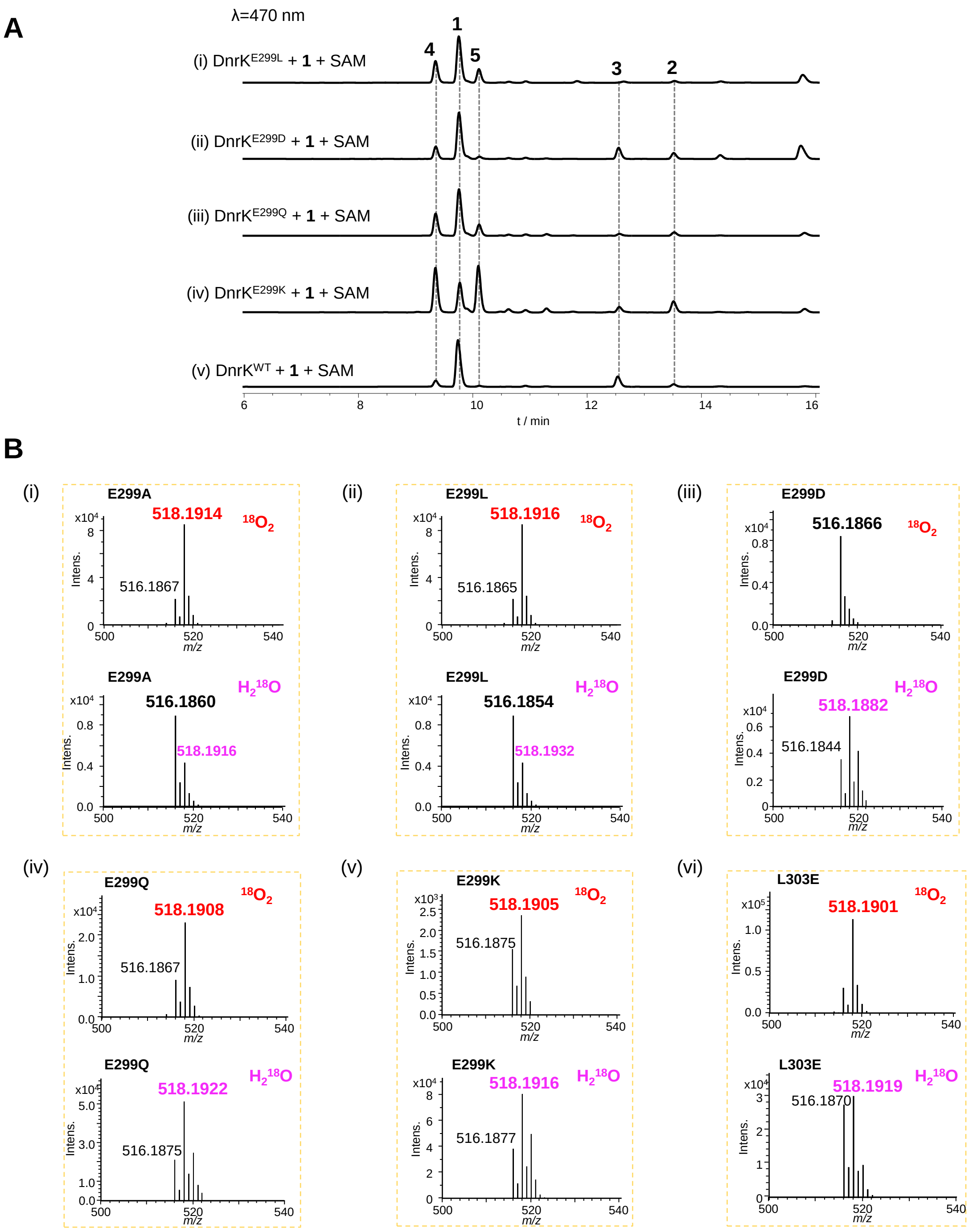

**Figure S28.** The enzymatic conversion of **1** catalyzed by DnrK$^{E299}$ mutants *in vitro* and isotopic labeling investigation of product **5**. (A) HPLC analysis of the conversion of **1** catalyzed by DnrK$^{E299}$ and its mutants (i) DnrK$^{E299L}$, (ii) DnrK$^{E299D}$, (iii) DnrK$^{E299Q}$, (iv) DnrK$^{E299K}$ and DnrK$^{WT}$ in presence of SAM, respectively. (B) HRESI-LCMS analysis of **5** in positive ion mode produced by (i) DnrK$^{E299A}$, (ii) DnrK$^{E299L}$, (iii) DnrK$^{E299D}$, (iv) DnrK$^{E299Q}$, (v) DnrK$^{E299K}$, and (vi) RdmB$^{L303E}$ mutants in $^{18}O_2$ and $H_2{}^{18}O$, respectively.

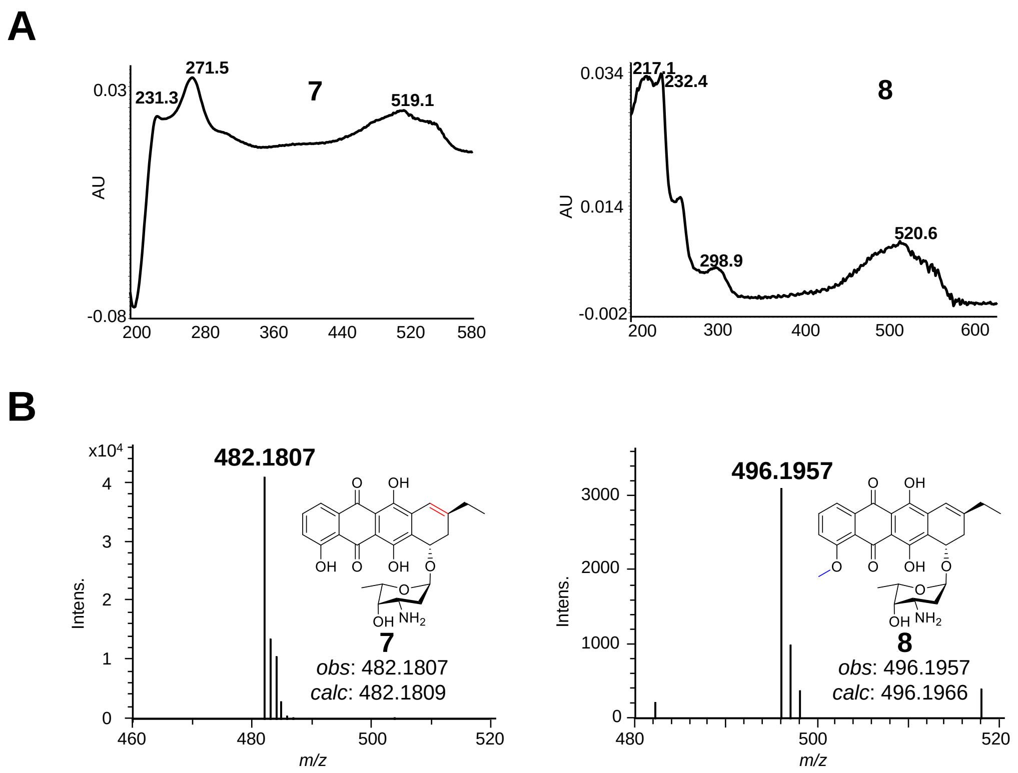

**Figure S29.** UV-Vis (A) and HRESI-LCMS (B) spectra of **7** and **8** in positive ion mode.

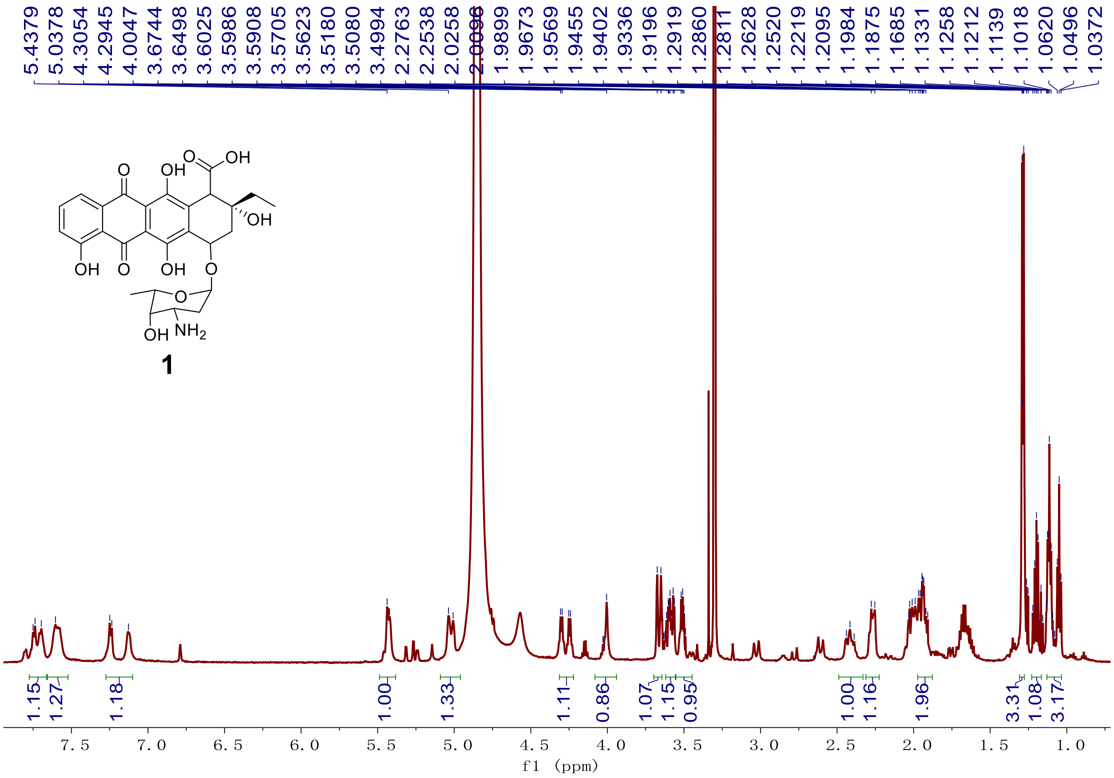

**Figure S30.** [1]H NMR spectrum (600 MHz, CD3OD) of compound **1**.

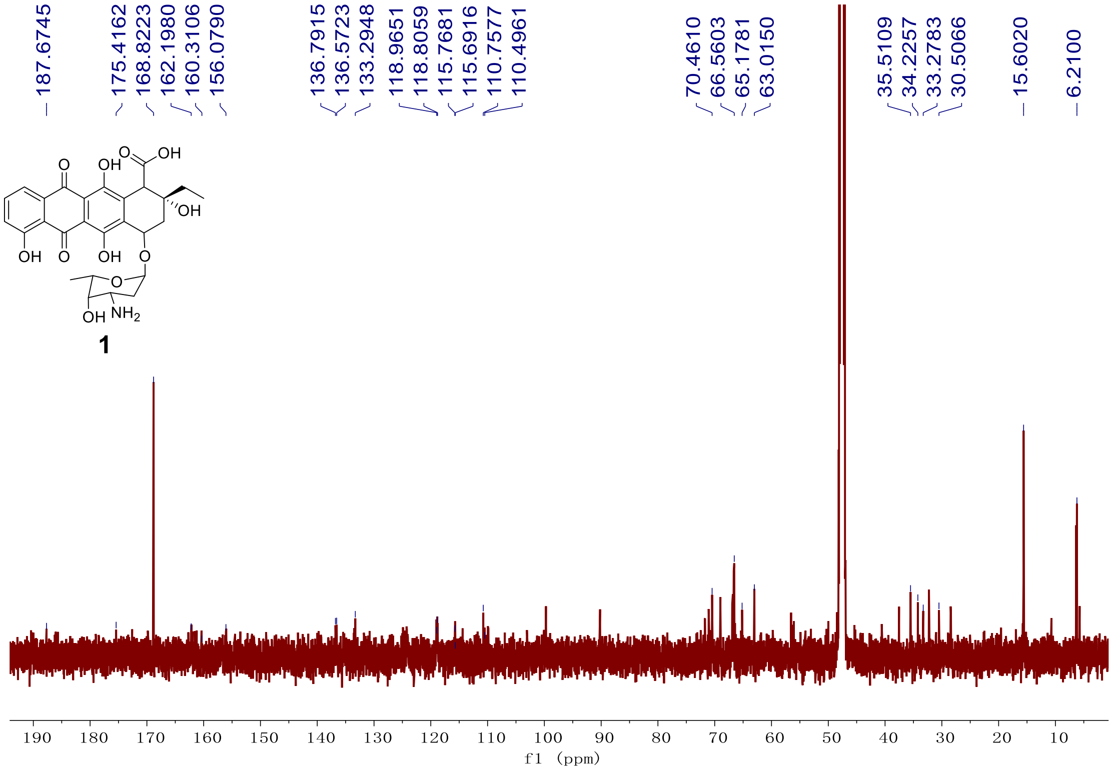

**Figure S31.** [13]C NMR spectrum (151 MHz, CD3OD) of compound **1**.

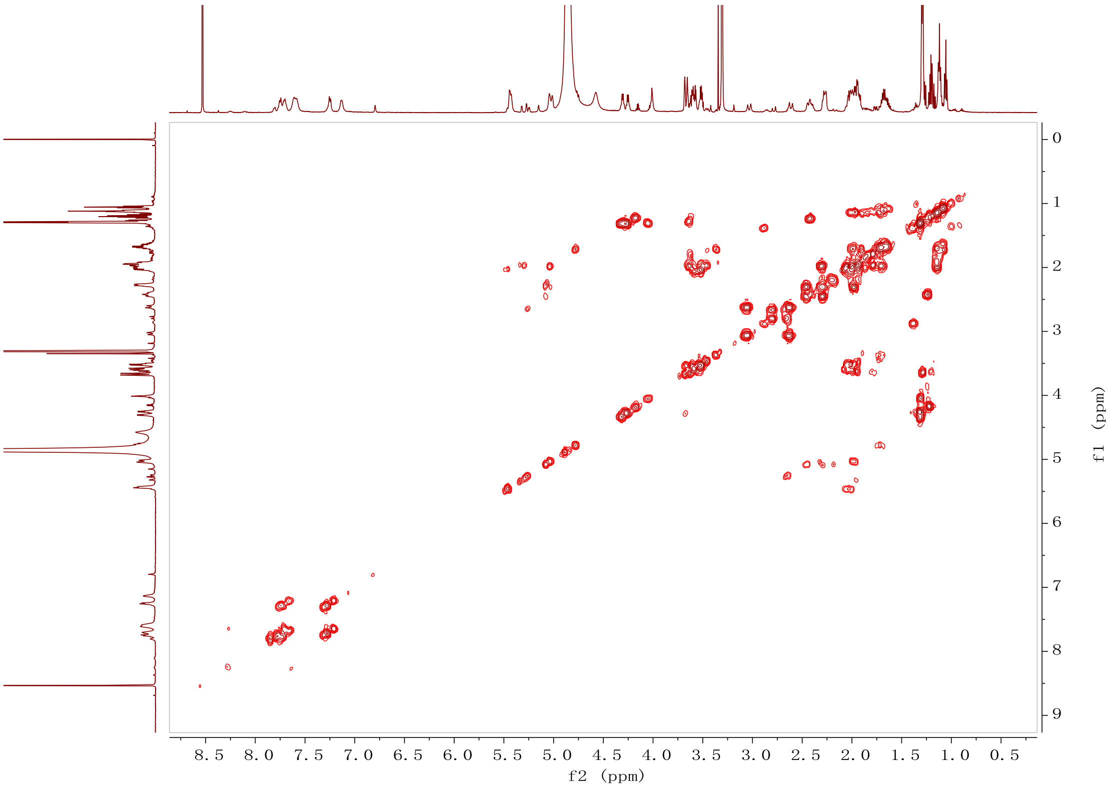

**Figure S32.** [1]H-[1]H COSY spectrum (600 MHz, $CD_3OD$) of compound **1**.

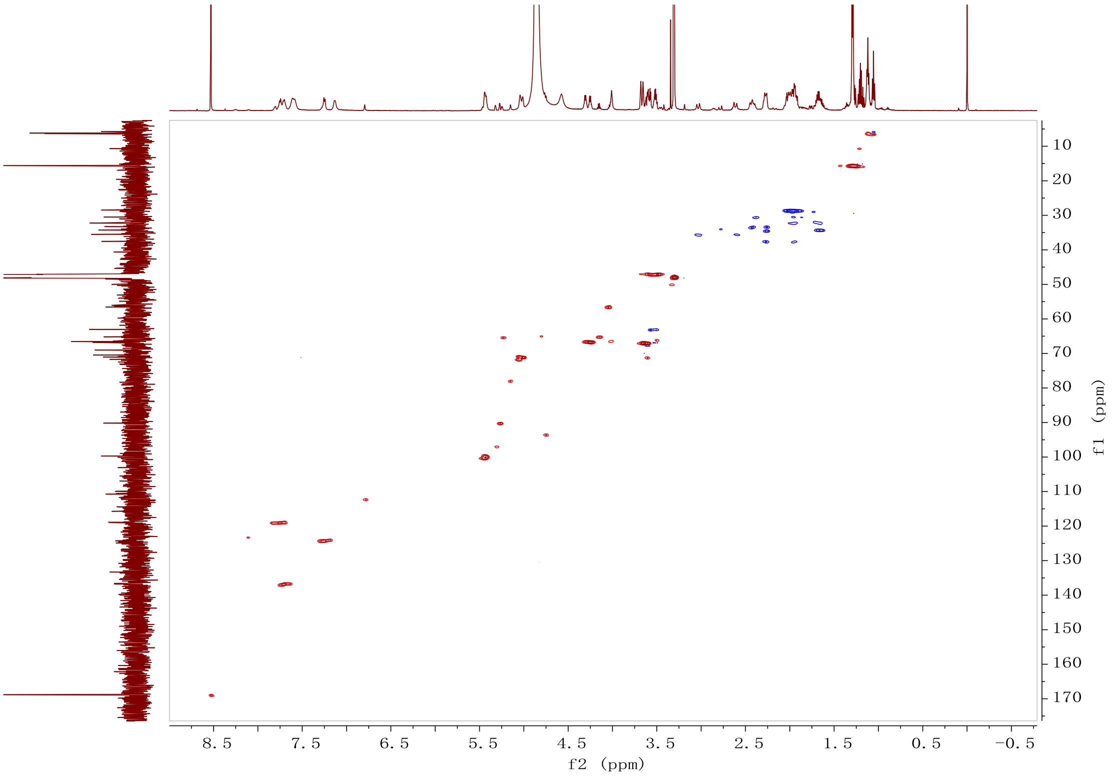

**Figure S33.** HSQC spectrum (600 MHz, $CD_3OD$) of compound **1**.

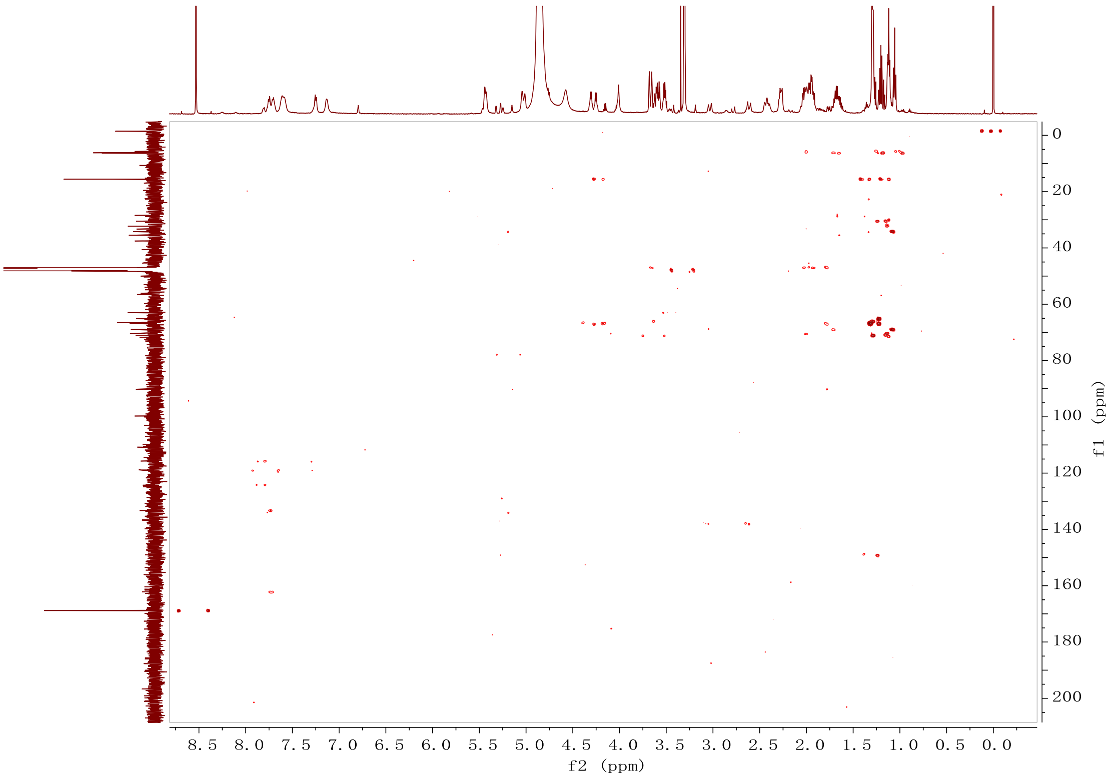

**Figure S34.** HMBC spectrum (600 MHz, CD3OD) of compound **1**.

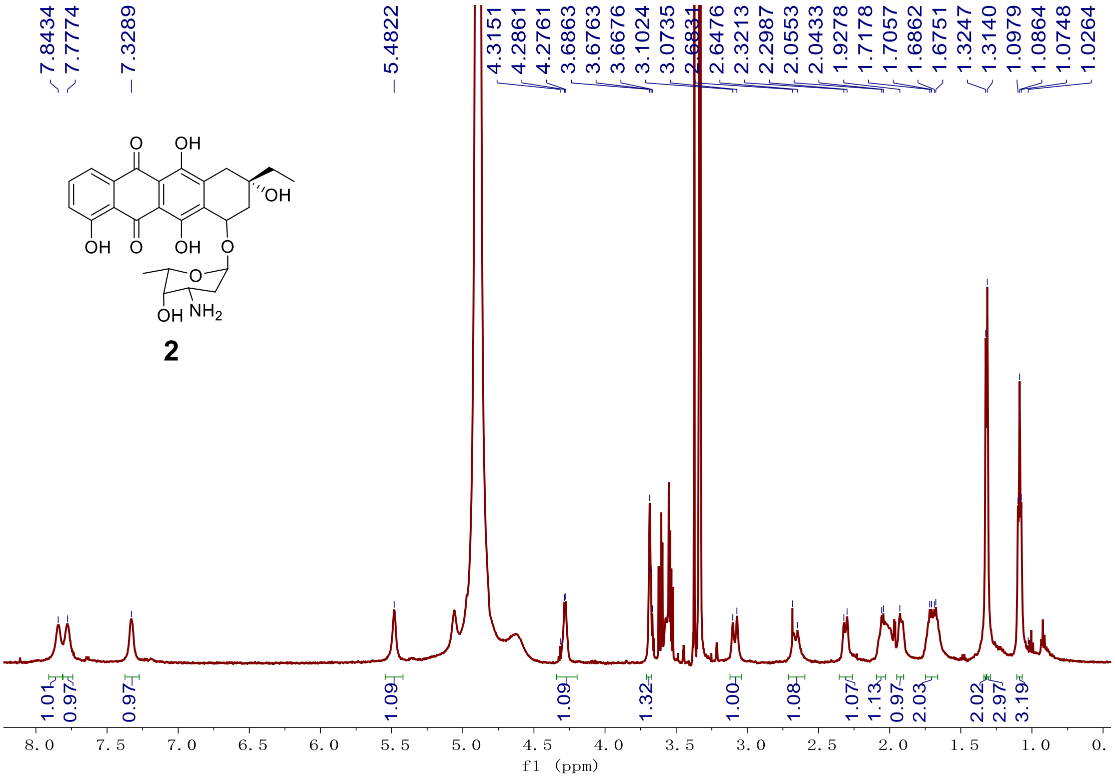

**Figure S35.** [1]H NMR spectrum (600 MHz, CD3OD) of compound **2**.

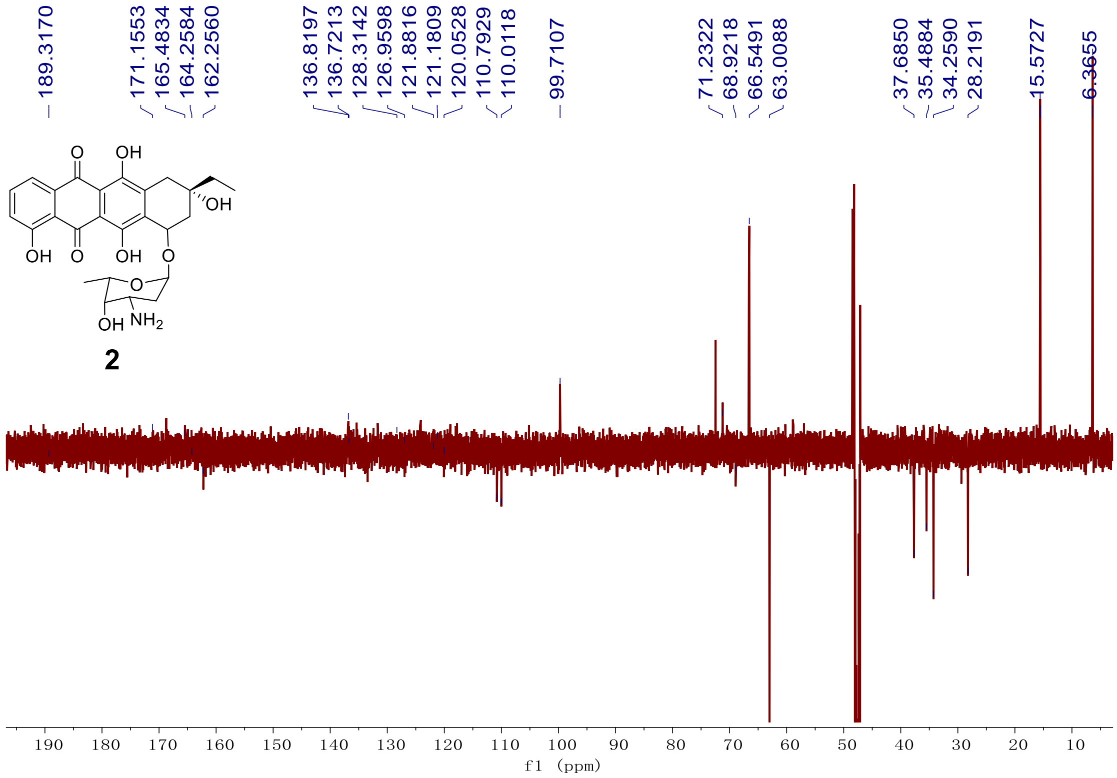

**Figure S36.** DEPT spectrum (151 MHz, $CD_3OD$) of compound **2**.

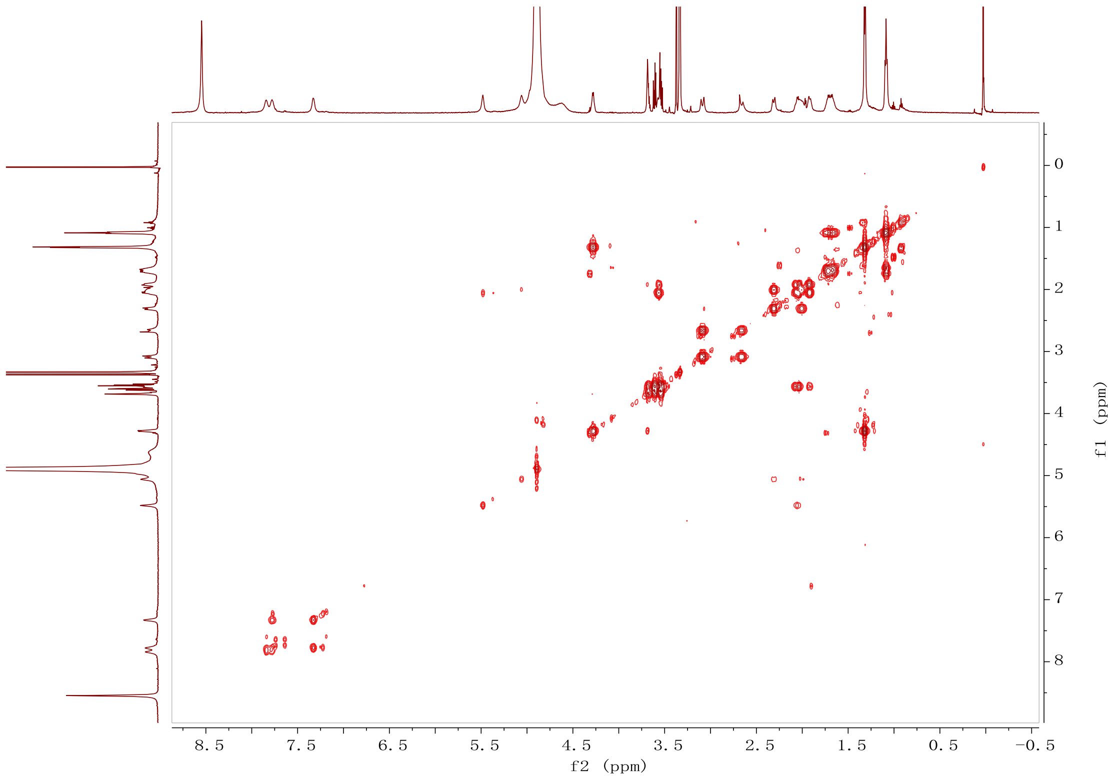

**Figure S37.** $^{1}H$-$^{1}H$ COSY spectrum (600 MHz, $CD_3OD$) of compound **2**.

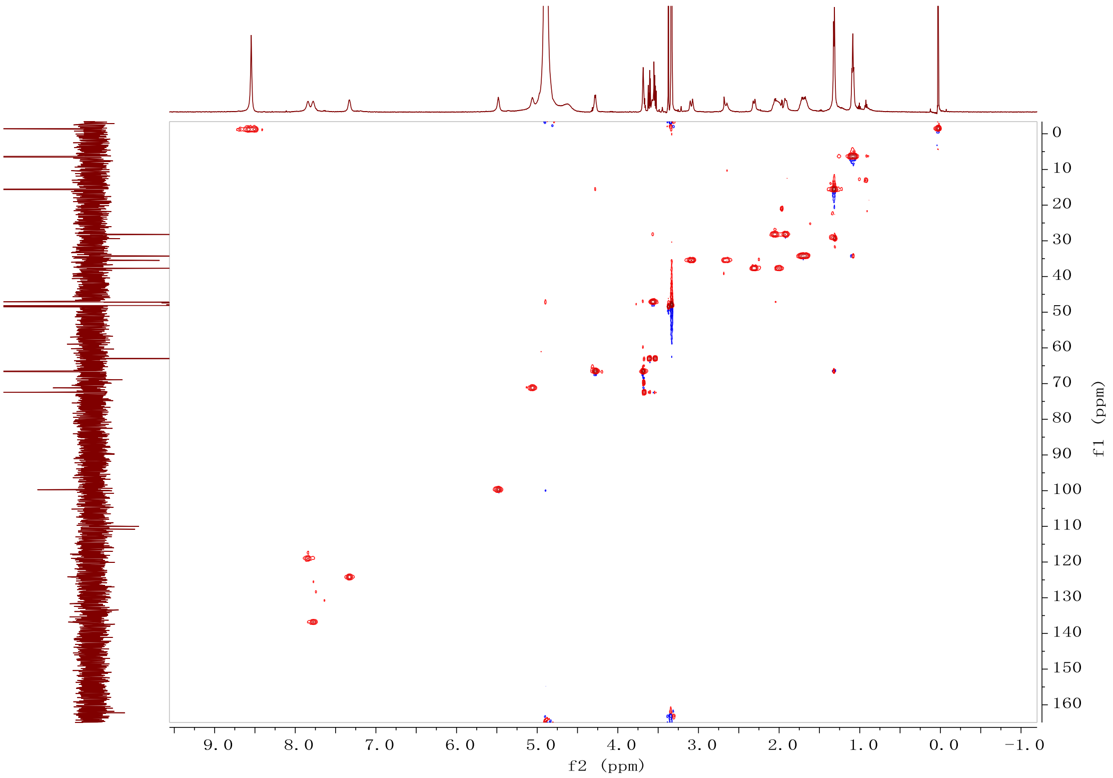

**Figure S38.** HSQC spectrum (600 MHz, $CD_3OD$) of compound **2**.

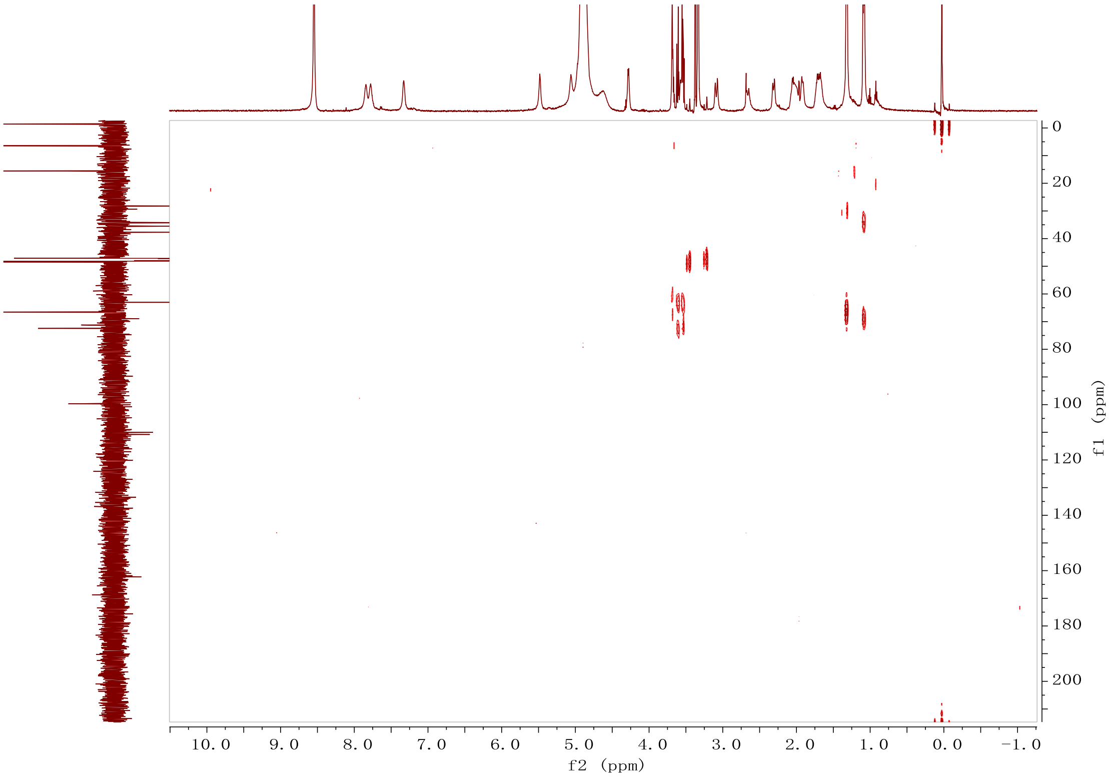

**Figure S39.** HMBC spectrum (600 MHz, $CD_3OD$) of compound **2**.

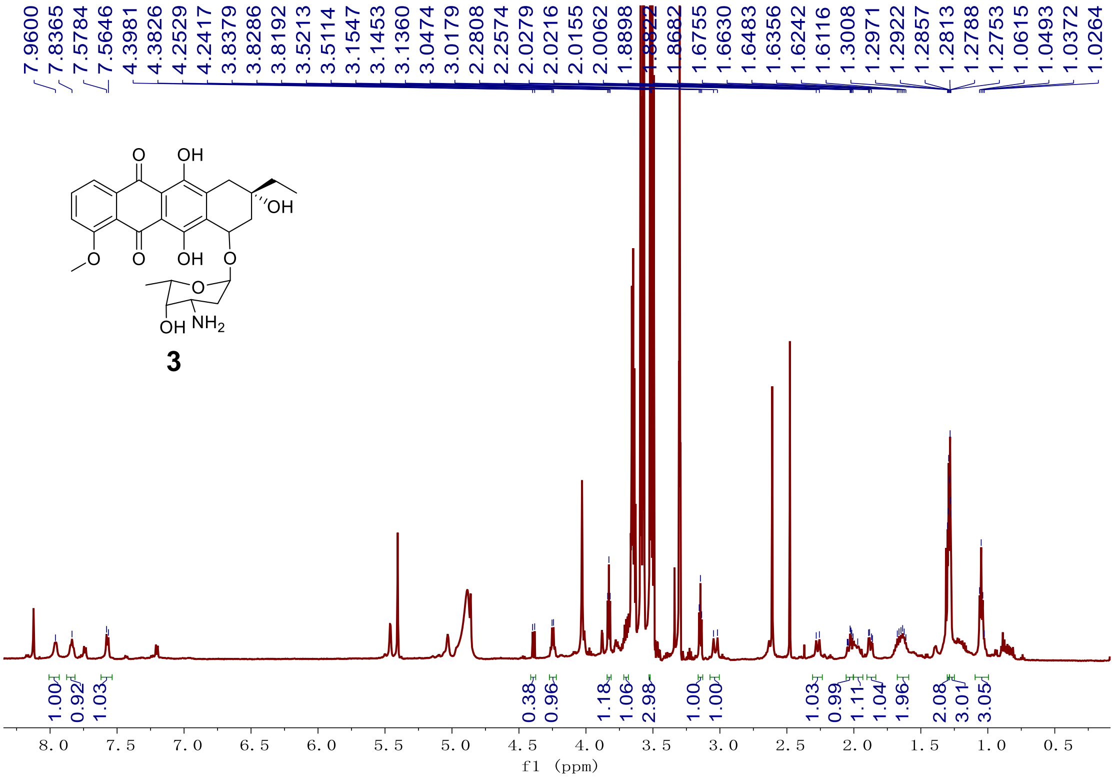

**Figure S40.** $^{1}$H NMR spectrum (600 MHz, CD3OD) of compound **3**.

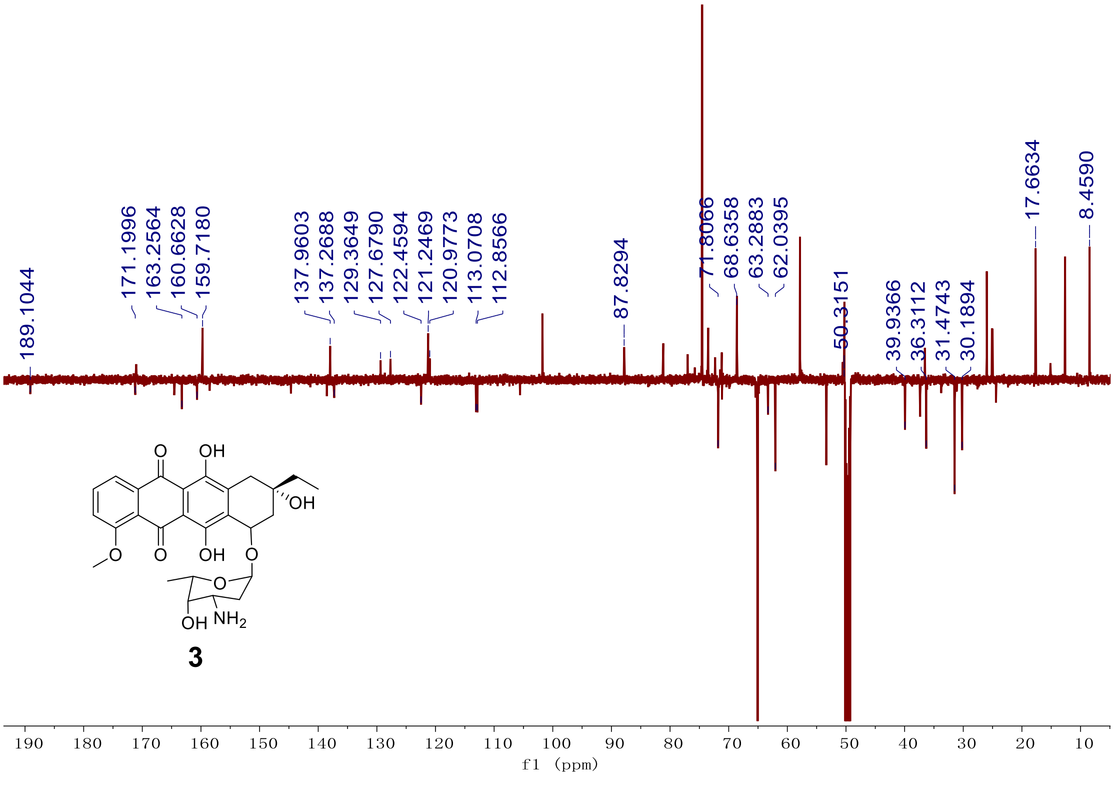

**Figure S41.** DEPT spectrum (151 MHz, CD3OD) of compound **3**.

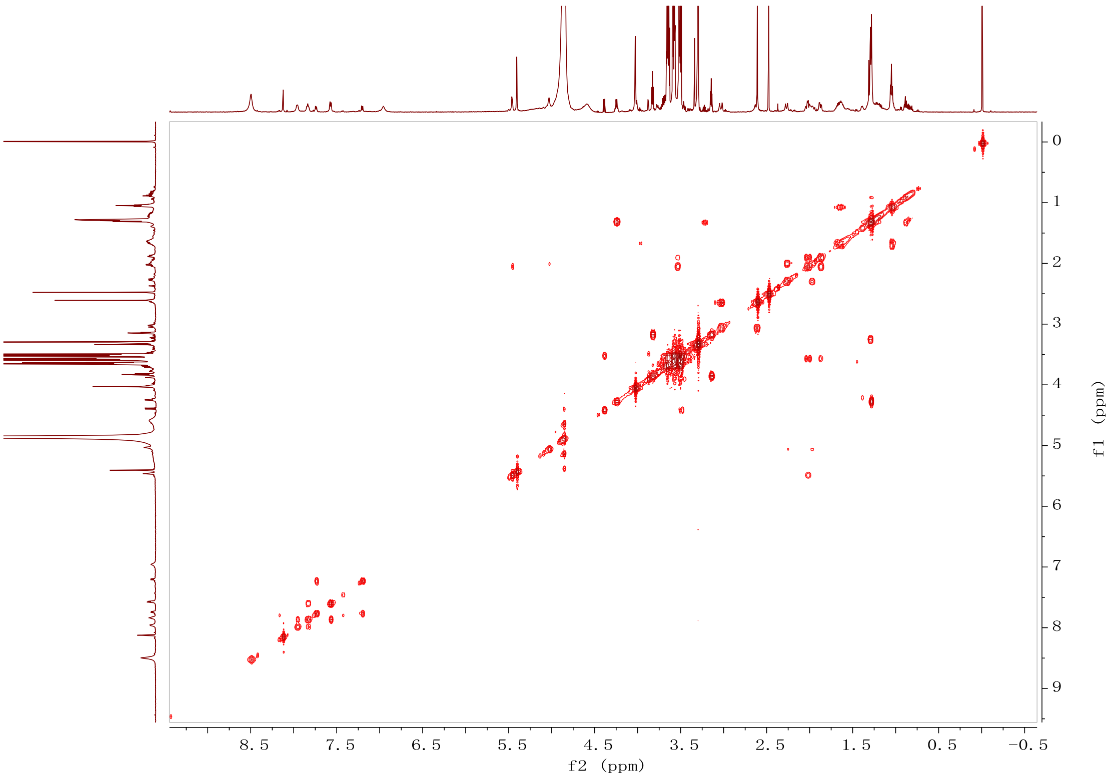

**Figure S42.** [1]H-[1]H COSY spectrum (600 MHz, $CD_3OD$) of compound **3**.

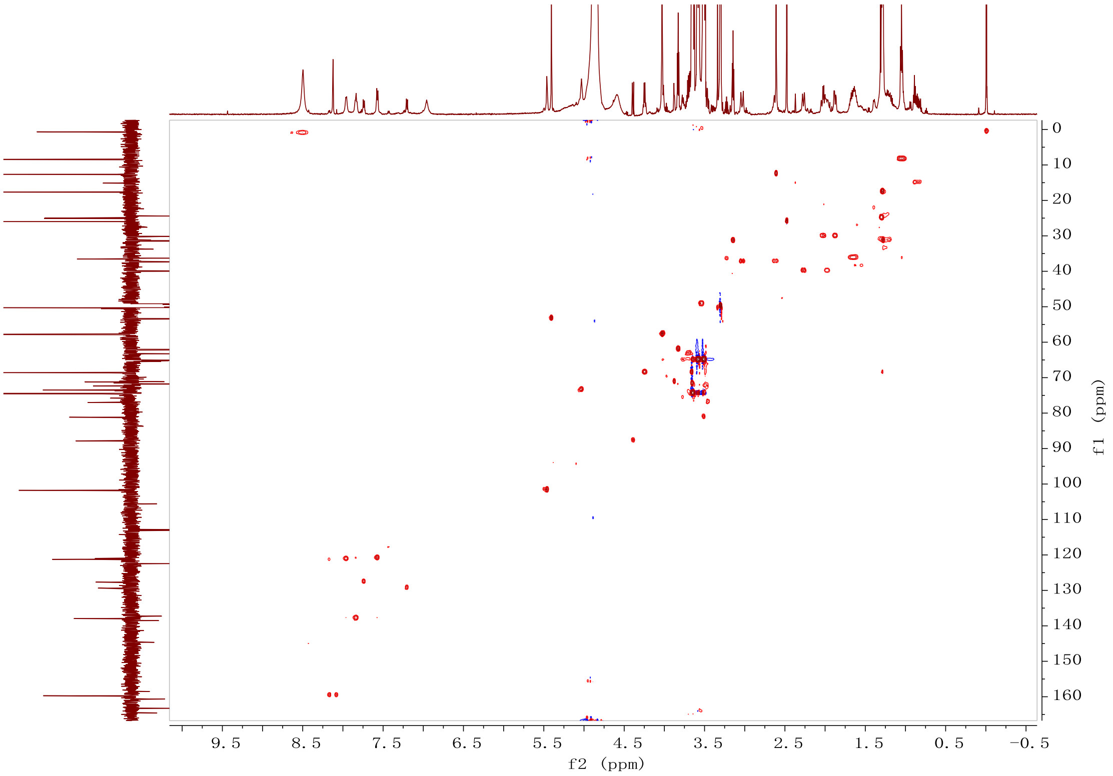

**Figure S43.** HSQC spectrum (600 MHz, $CD_3OD$) of compound **3**.

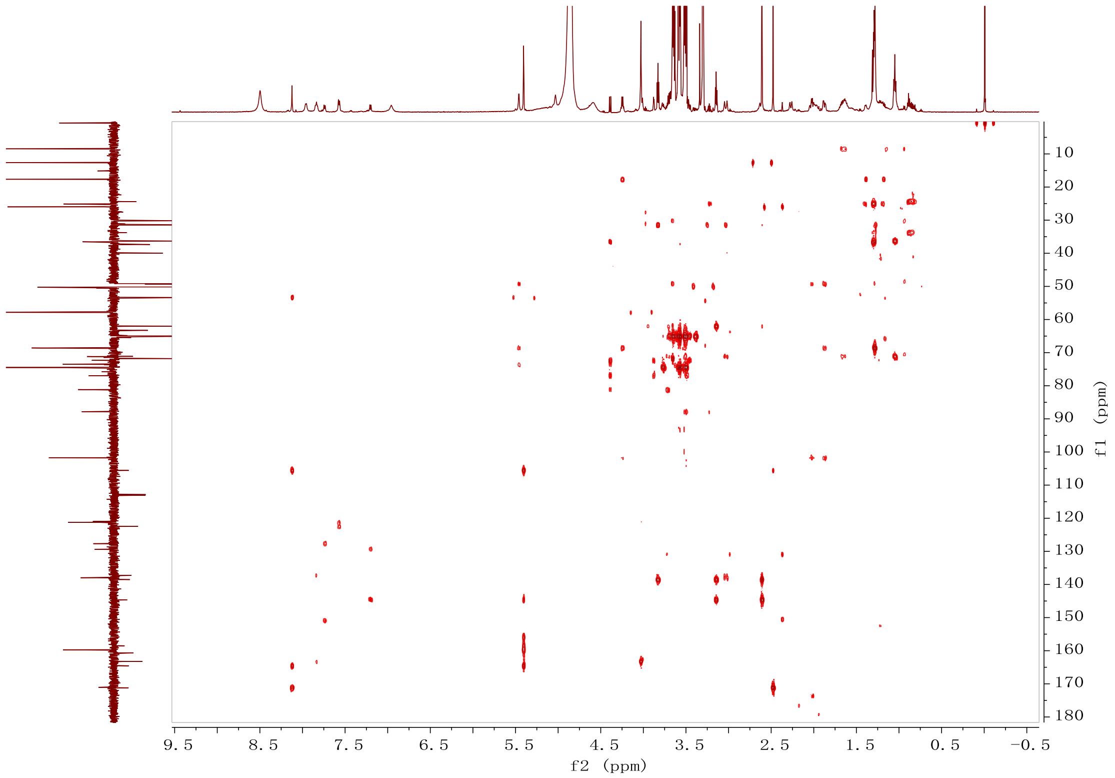

**Figure S44.** HMBC spectrum (600 MHz, CD3OD) of compound **3**.

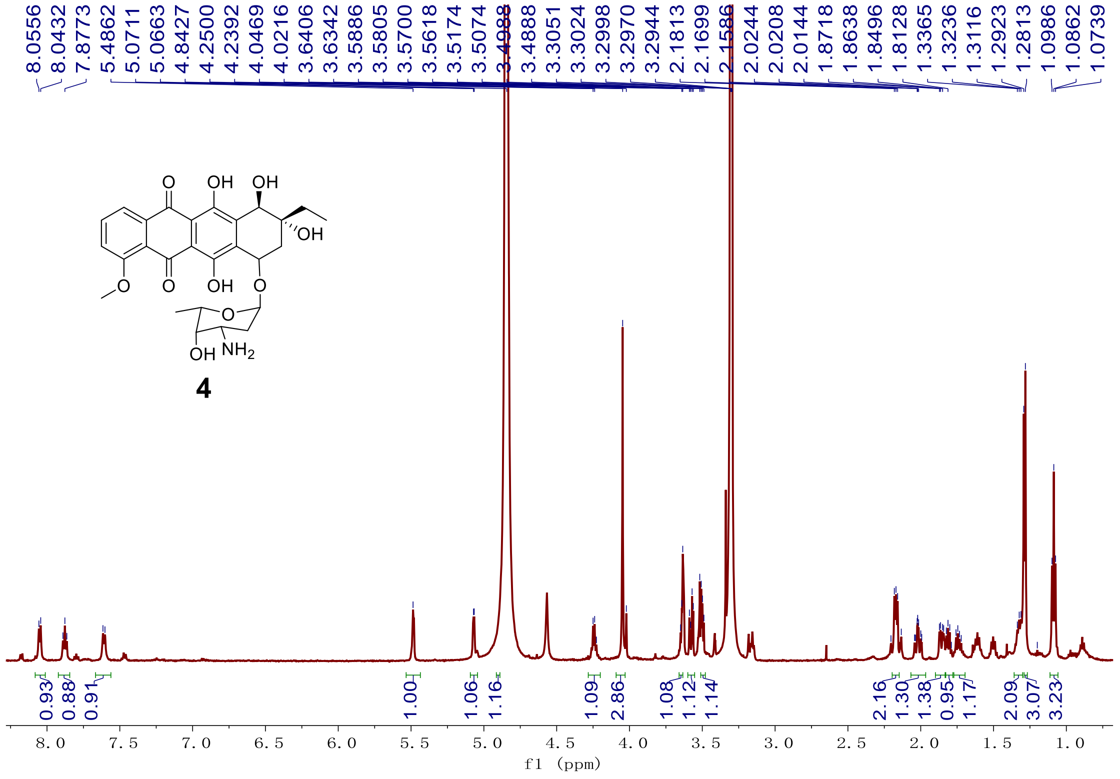

**Figure S45.** [1]H NMR spectrum (600 MHz, CD3OD) of compound **4**.

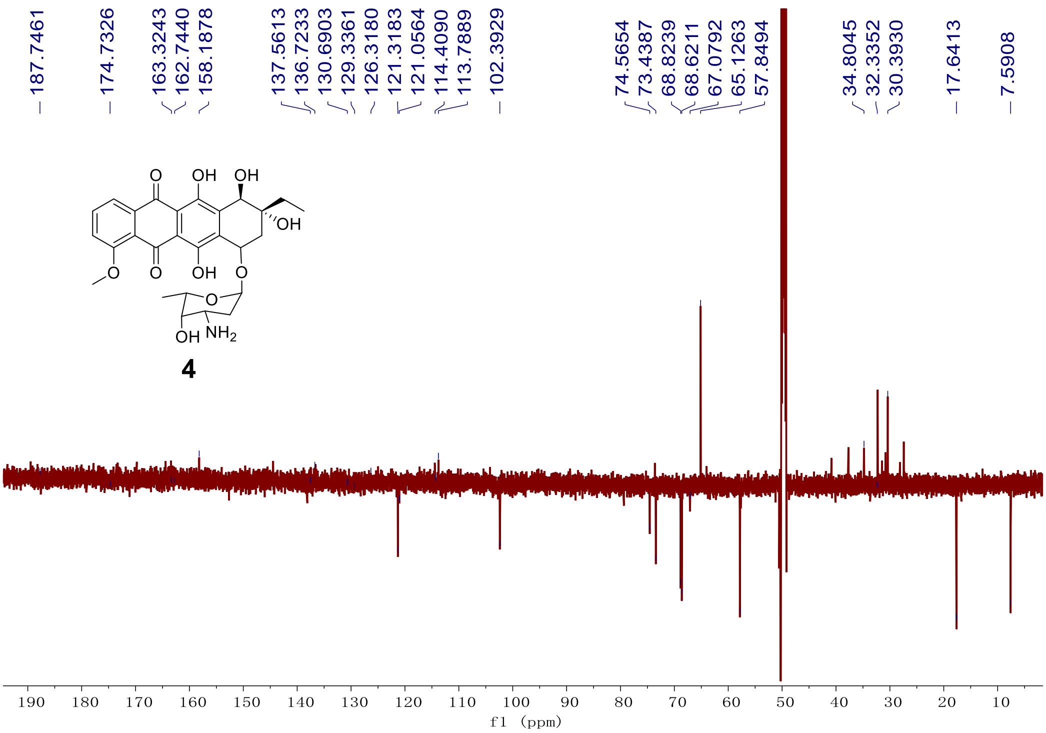

**Figure S46.** DEPT spectrum (151 MHz, $CD_3OD$) of compound **4**.

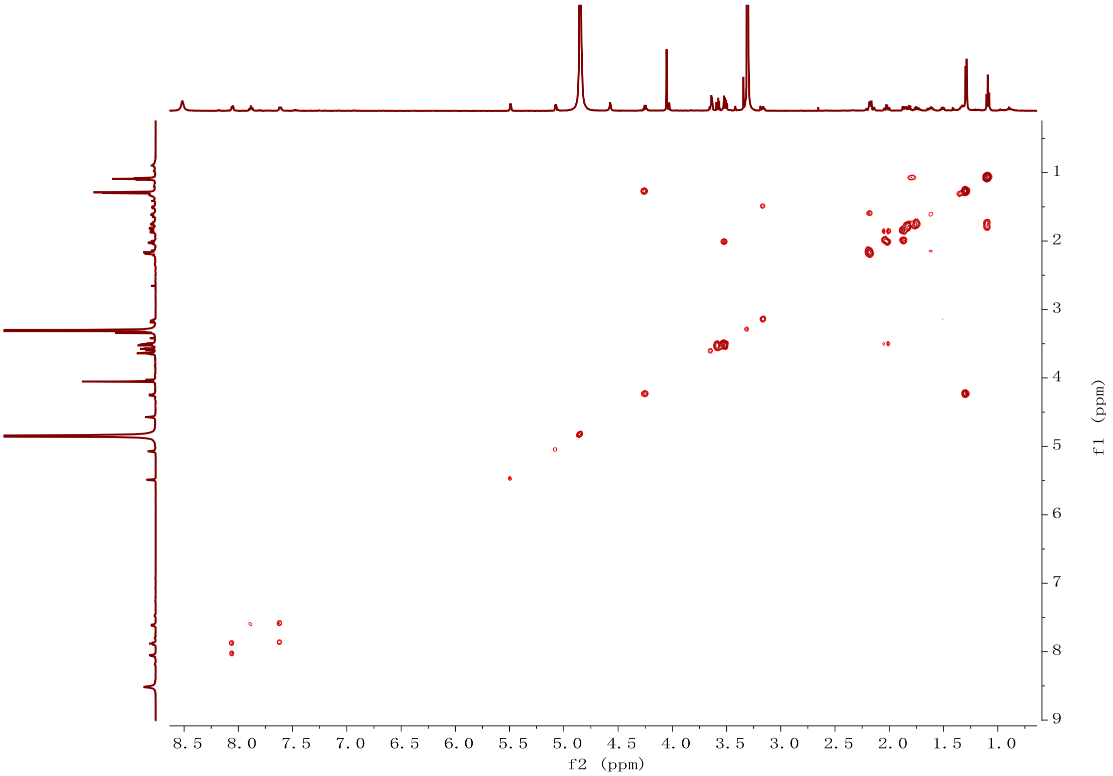

**Figure S47.** $^{1}H$-$^{1}H$ COSY spectrum (600 MHz, $CD_3OD$) of compound **4**.

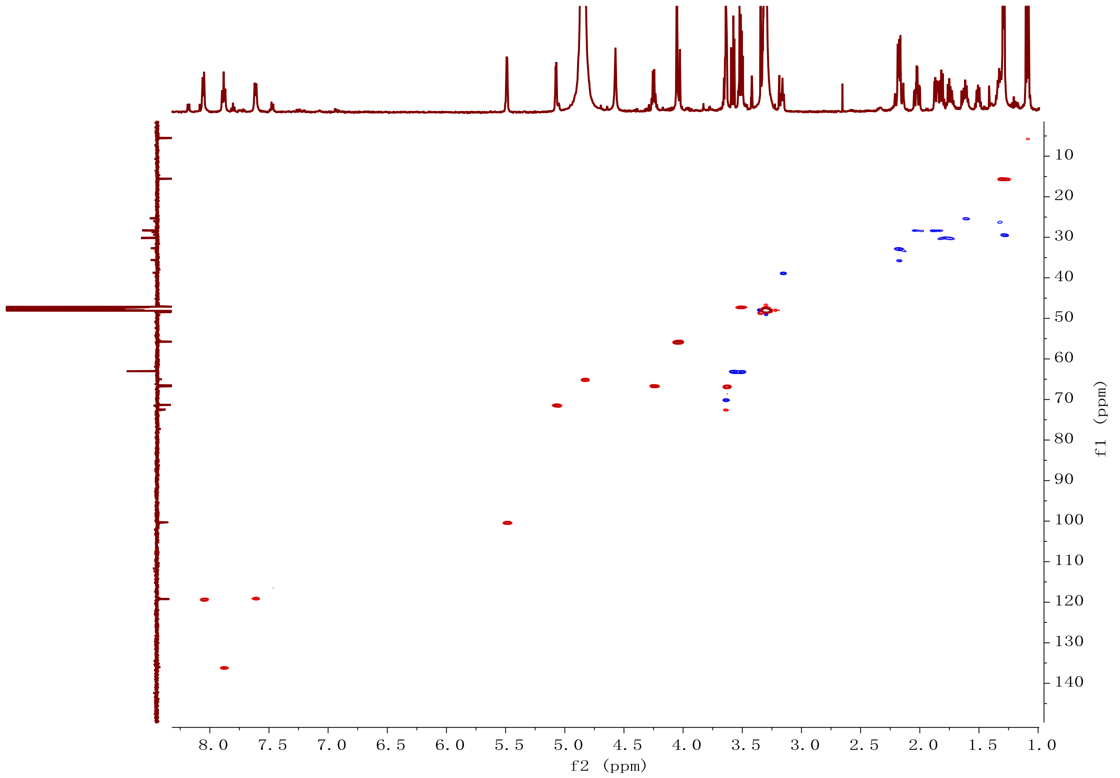

**Figure S48.** HSQC spectrum (600 MHz, $CD_3OD$) of compound **4**.

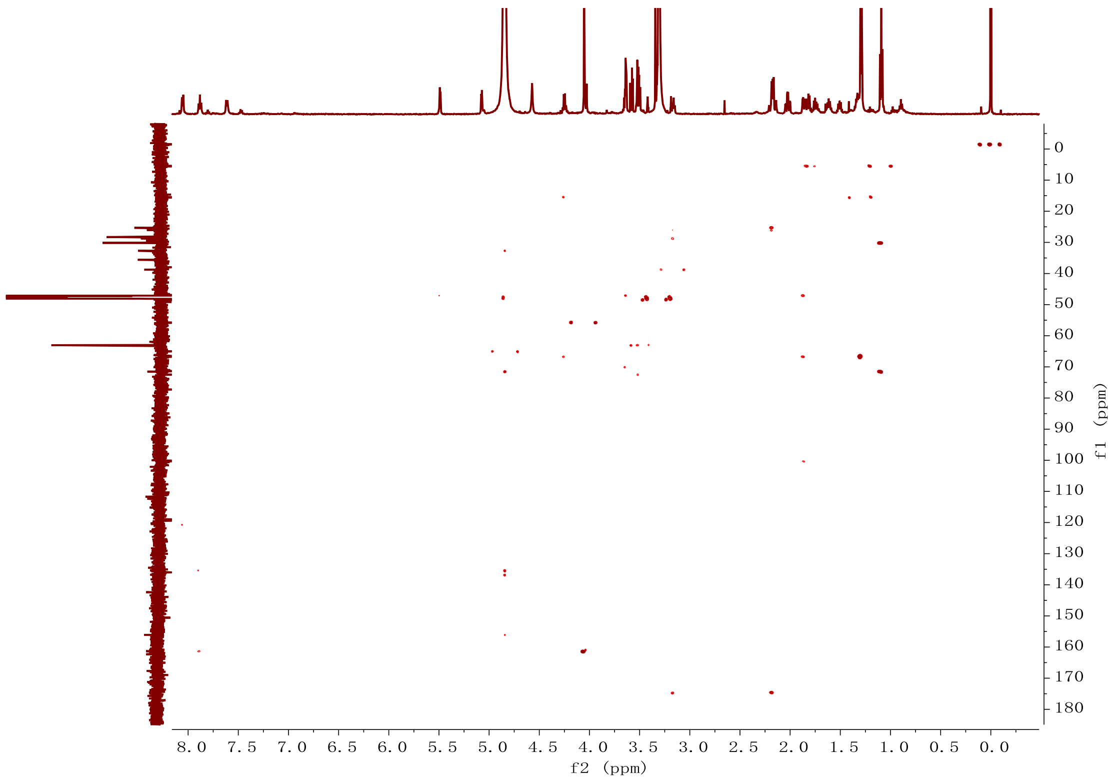

**Figure S49.** HMBC spectrum (600 MHz, $CD_3OD$) of compound **4**.

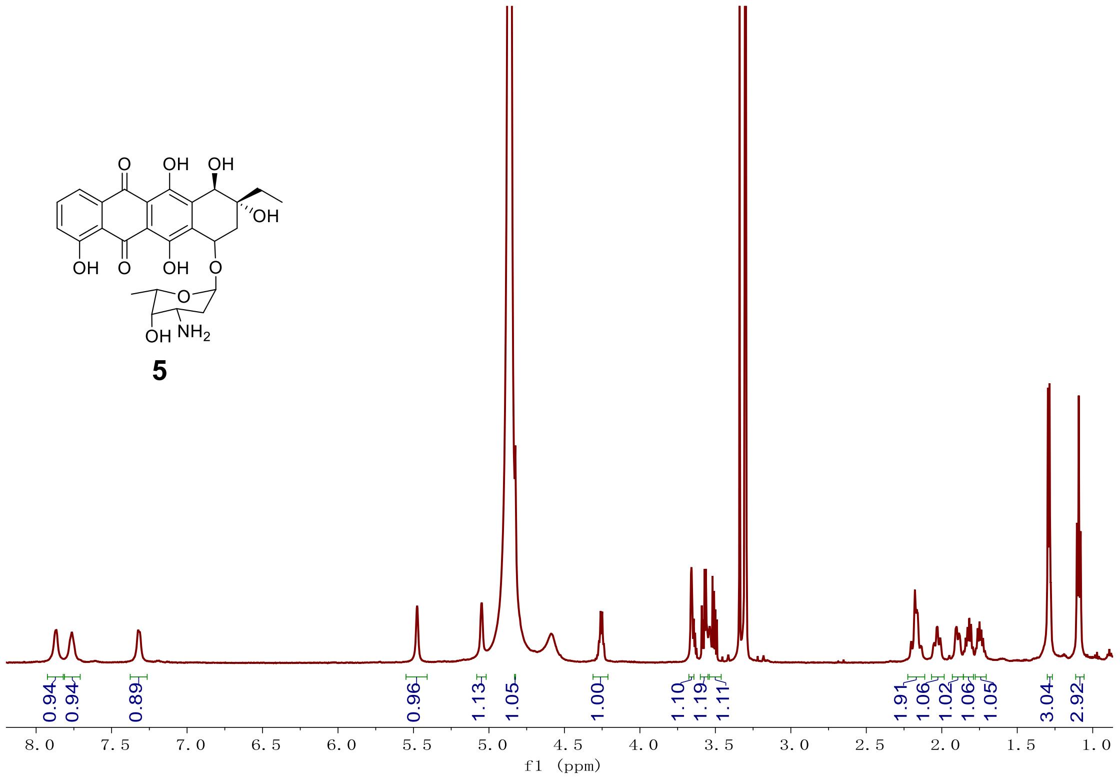

**Figure S50.** $^{1}$H NMR spectrum (600 MHz, $CD_3OD$) of compound **5**.

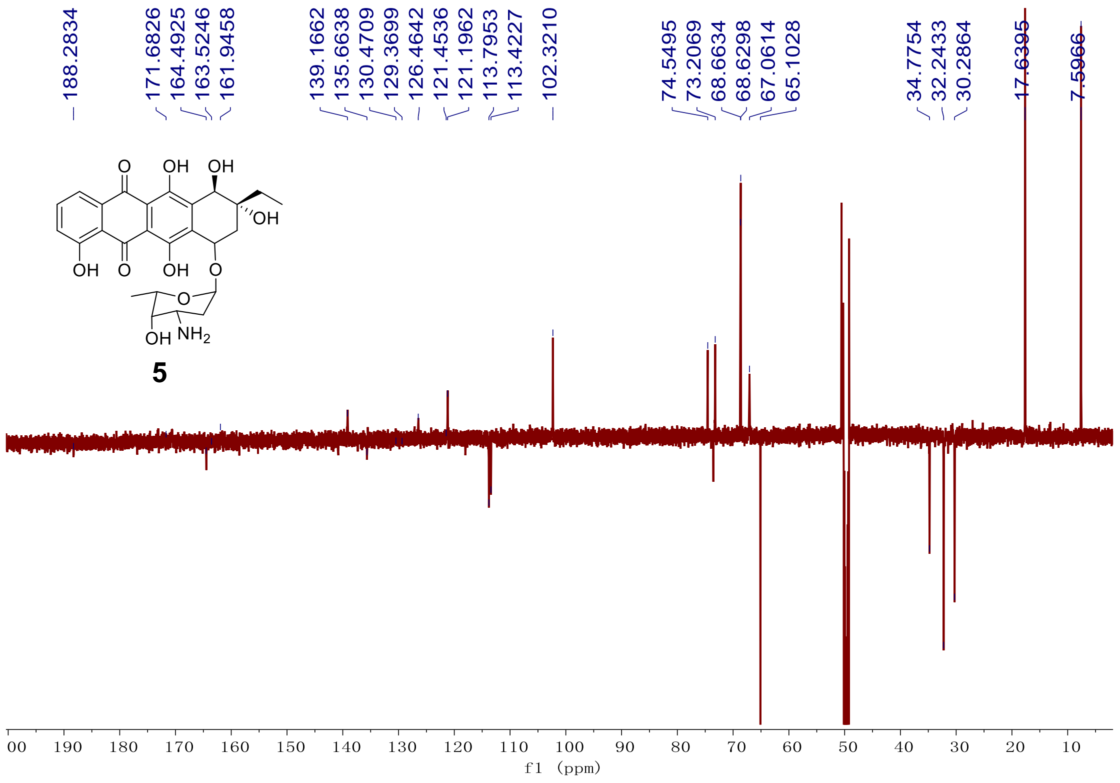

**Figure S51.** DEPT spectrum (151 MHz, $CD_3OD$) of compound **5**.

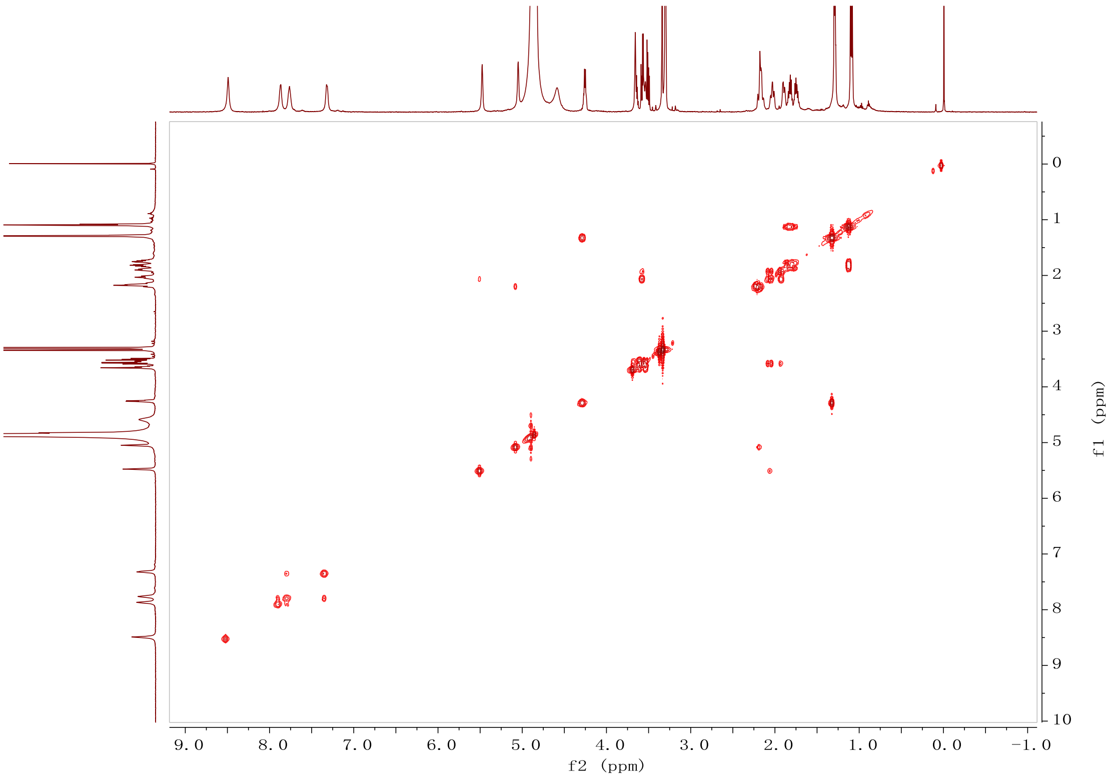

**Figure S52.** [1]H-[1]H COSY spectrum (600 MHz, $CD_3OD$) of compound **5**.

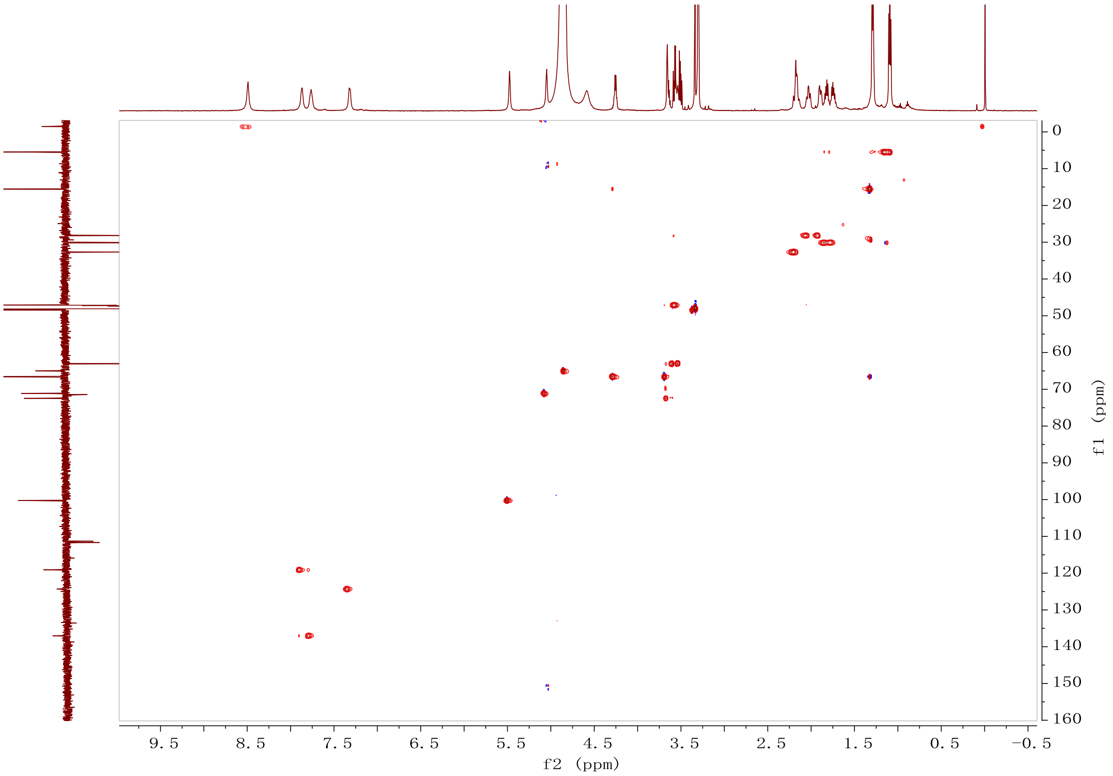

**Figure S53.** HSQC spectrum (600 MHz, $CD_3OD$) of compound **5**.

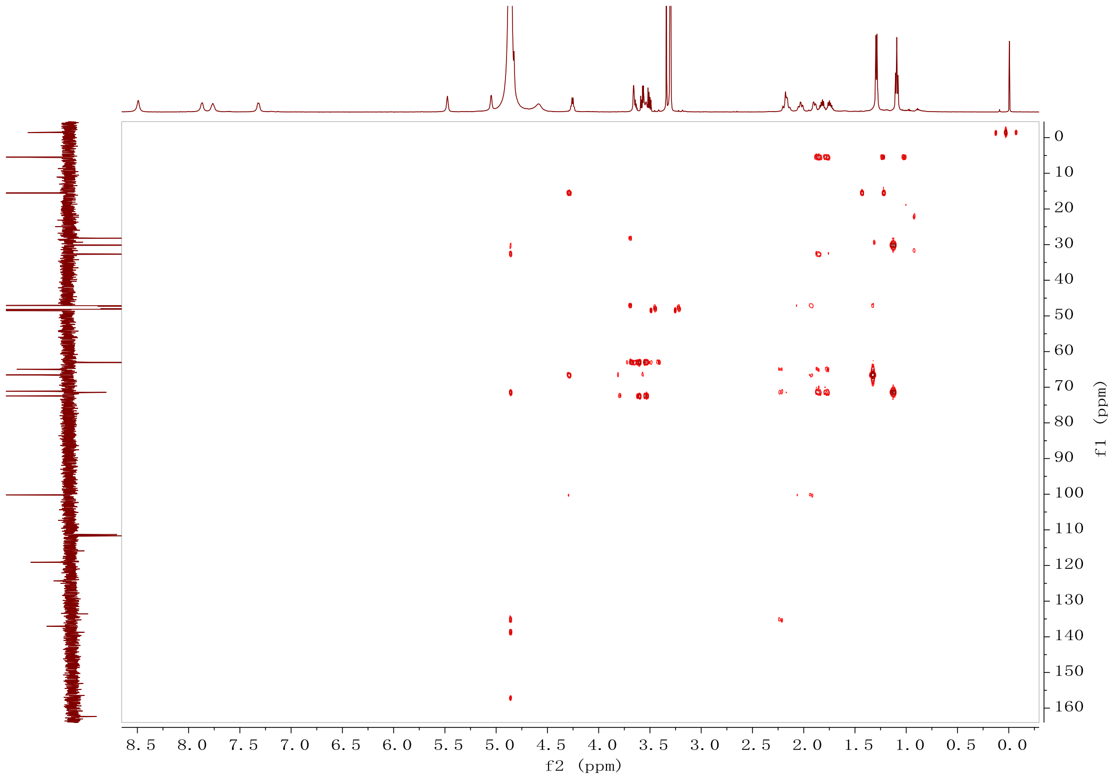

**Figure S54.** HMBC spectrum (600 MHz, $CD_3OD$) of compound **5**.
